# YAP1-2α–TAZ heterodimerization promotes BRD4-associated LLPS-like assembly at candidate super-enhancers and cancer cell malignancy

**DOI:** 10.1101/2025.01.28.635202

**Authors:** Takuya Ooki, Takeru Hayashi, Yui Yamamoto, Masanori Hatakeyama

## Abstract

YAP1 is a transcriptional coactivator expressed as multiple splice isoforms, but how individual isoforms specify malignant cell states remains unclear. Here, we identify YAP1-2α as an isoform that drives a distinct transcriptional state in cancer cells. The YAP1-2α–TAZ heterodimer, together with TEAD, forms BRD4-associated nuclear assemblies with LLPS-like properties. These assemblies are enriched at candidate super-enhancer-associated regions and promote transcription. Disrupting YAP1-2α–TAZ heterodimerization, TEAD binding, or BRD4 interaction impairs assembly formation and associated phenotypes. Exon-6 exclusion promotes YAP1-2α-associated nuclear puncta, invasion, and therapeutic resistance in high-grade serous ovarian cancer (HGSOC), and enhances intraperitoneal tumor progression in gastric cancer xenografts. Gastric cancer cells selected through peritoneal dissemination exhibit increased YAP1-2α, nuclear YAP1 puncta, and therapeutic resistance. In HGSOC specimens, YAP1-2α abundance, YAP1–BRD4 proximity, and nuclear YAP1 puncta are associated with advanced stages. Thus, YAP1-2α drives a BRD4-associated condensate-like transcriptional state linked to malignant progression across multiple cancer contexts.

## INTRODUCTION

Alternative splicing expands proteomic diversity by generating multiple protein isoforms from a single gene. Its dysregulation is common during cancer progression and can influence cell proliferation, apoptosis, invasion, metastasis, therapeutic resistance, and immune evasion.^1–3^ Defining the functions of cancer-associated splice isoforms may therefore reveal mechanisms of tumor progression and therapeutic vulnerabilities.

YAP1 and its paralog TAZ are transcriptional coactivators that lack intrinsic DNA-binding activity and regulate gene expression primarily through TEAD family transcription factors (TEAD1–4).^4,5^ YAP1/TEAD and TAZ/TEAD complexes integrate mechanical, metabolic, and cell-density signals to regulate cell proliferation, differentiation, and tumor progression.^6,7^ In cancer, YAP1 and TAZ can promote epithelial-to-mesenchymal transition (EMT), stem-like phenotypes, invasion, and resistance to anticancer therapies.^8–11^ Although YAP1 and TAZ share many functions, they can occupy distinct genomic sites and regulate partially overlapping transcriptional programs.^12,13^ Unlike TAZ, YAP1 undergoes complex alternative splicing and produces at least eight human isoforms.^14–16^ However, the structural basis by which individual YAP1 isoforms generate distinct transcriptional outputs remains incompletely understood. Alternative splicing of exons 4, 5, and 6 generates YAP1 isoforms with distinct domain compositions. Exon 4 encodes the second WW domain (WW2), distinguishing YAP1-1, which contains only the first WW domain (WW1), from YAP1-2, which contains both WW1 and WW2. WW domains canonically recognize PPxY motifs and can modulate YAP1 regulation and subcellular localization.^17,18^ Each YAP1-1 and YAP1-2 subgroup comprises four variants, α, β, γ, and δ, generated by alternative splicing within the transcriptional activation domain. Exon 5 has an alternative splice donor site that produces the 3′-extended exon 5.2, encoding a 4-amino-acid β segment (VRPQ), whereas exon 6 encodes a 16-amino-acid γ segment. The γ isoforms contain exon 5.1 and exon 6, whereas the δ isoforms contain exon 5.2 together with exon 6. The β isoforms contain exon 5.2 but lack exon 6, whereas the α isoforms contain exon 5.1 and lack exon 6. Thus, only the α isoforms retain an intact leucine zipper within the transcriptional activation domain, which mediates protein–protein interactions through a coiled-coil region.^19^ The biological functions of individual YAP1 isoforms in cancer remain poorly defined.

Biomolecular condensates are dynamic, membraneless assemblies formed through multivalent interactions among proteins, nucleic acids, and chromatin-associated factors.^20,21^ In the nucleus, transcription factors and coactivators can assemble into condensate-like structures at highly active regulatory regions, including super-enhancers, thereby facilitating transcriptional control.^22–24^ Aberrant enhancer-associated transcriptional programs contribute to malignant cell states, including invasion, drug resistance, and tumor progression.^25–29^

In this study, we identify YAP1-2α as a splice isoform that heterodimerizes with TAZ through its leucine zipper and cooperates with TEAD and BRD4 to form dynamic, liquid-liquid phase separation (LLPS)-like nuclear condensates. These assemblies are associated with candidate super-enhancer regulatory regions, local chromatin interactions, and activation of EMT- and stemness-associated genes. Functionally, YAP1-2α promotes sphere formation, invasion, therapeutic resistance, and intraperitoneal tumor progression. Our findings identify YAP1 splicing as a potential regulator of condensate-associated transcriptional states and suggest that YAP1-2α production or YAP1-2α/TAZ/TEAD/BRD4-associated interactions may represent therapeutic vulnerabilities in selected tumors.

## RESULTS

### YAP1-2α uniquely forms dynamic nuclear condensates with liquid-like properties

To compare the eight known YAP1 splice isoforms (Figure 1A; four YAP1-1 and four YAP1-2 isoforms), we generated *YAP1*-knockout UWB1.289 human high-grade serous ovarian cancer (HGSOC) cells using CRISPR–Cas9. In parental UWB1.289 cells, YAP1-2 isoforms were detected (Figure S1A). Immunoblotting confirmed loss of endogenous YAP1 and comparable expression of the eight C-terminal mVenus-tagged isoforms (YAP1-mVenus) after transient transfection (Figure S1A), enabling direct comparison of their subnuclear behaviors.

**Figure 1.**
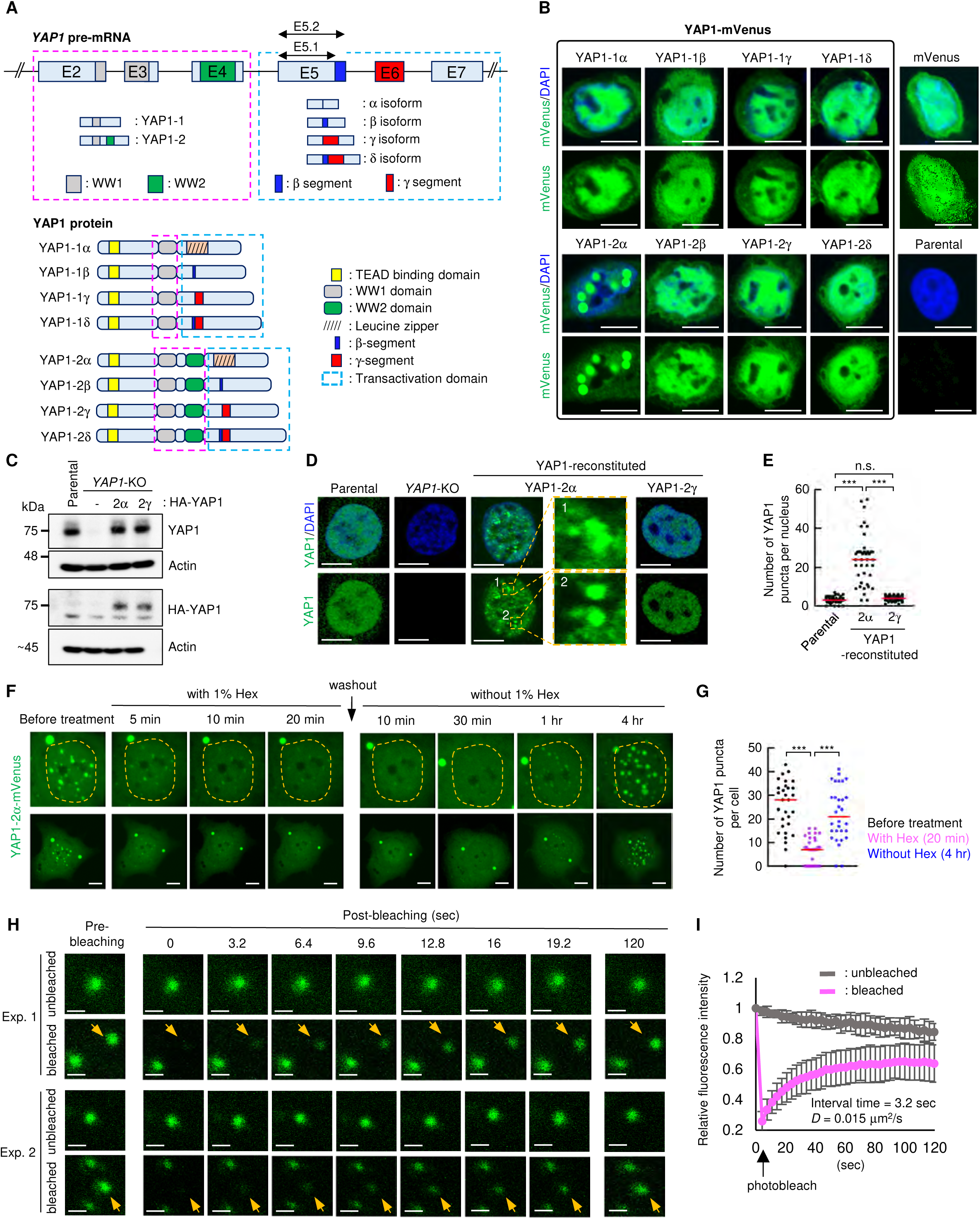
YAP1-2α uniquely forms dynamic nuclear puncta with LLPS-like properties among YAP1 splice isoforms. (**A**) Alternative splicing of *YAP1* pre-mRNA and the eight resulting protein isoforms. (**B**) Representative images of *YAP1*-KO UWB1.289 cells expressing C-terminal mVenus-tagged YAP1 splice isoforms (mVenus-YAP1). See Figure S1A for protein expression and quantification. See Figure S3F for protein expression. (**C–E**) Clonal *YAP1*-KO UWB1.289 cells reconstituted with YAP1-2α or YAP1-2γ were analyzed by immunoblotting (C), YAP1 immunofluorescence (D), and quantification of nuclear puncta (E). (**F** and **G**) Live-cell imaging and quantification of YAP1-2α-mVenus puncta before, during, and after 1% Hex treatment. (**H** and **I**) FRAP analysis of YAP1-2α-mVenus puncta. Arrows indicate bleached puncta; unbleached puncta served as controls (H). Quantification shows fluorescence recovery and diffusion coefficients (I). Data are mean ± SD. \*\*\**p* < 0.001; n.s., not significant; Mann-Whitney U test. *n* = 40 nuclei per group (E), 30 cells from three independent biological replicates (G), and 12 puncta per condition (I). Scale bars, 10 μm (B, D, and F) and 1 μm (H). The same cell lysates were resolved on separate SDS-PAGE gels to detect endogenous YAP1 and HA-tagged proteins, with their respective loading controls (C). See also Figures S1 and S2.

Among the eight isoforms, YAP1-2α alone consistently formed prominent nuclear puncta under identical imaging conditions; the other seven exhibited predominantly diffuse nuclear localization (Figures 1B and S1A). This difference was not explained by gross variation in protein abundance or overt cytoplasmic redistribution. To exclude tag-related effects, we examined N-terminal HA- or mVenus-tagged YAP1-2 isoforms. Only YAP1-2α formed nuclear puncta, whereas YAP1-2β, YAP1-2γ, and YAP1-2δ remained diffusely distributed (Figures S1B–S1E), indicating an intrinsic isoform-selective property of YAP1-2α.

We therefore compared YAP1-2α with YAP1-2γ, the predominant endogenous YAP1-2 isoform in parental UWB1.289 cells (Figure S1F). In *YAP1*-knockout cells reconstituted with HA-tagged YAP1-2α or YAP1-2γ at near-endogenous protein levels (Figure 1C), YAP1-2α, but not YAP1-2γ, formed discernible nuclear condensates (Figures 1D and 1E). Thus, this differential behavior was maintained at expression levels comparable to endogenous YAP1.

Because biomolecular condensates often display dynamic and liquid-like properties,^22–24^ we next characterized the physical behavior of YAP1-2α puncta. Treatment with 1,6-hexanediol markedly reduced their number (Figures 1F and 1G), and time-lapse imaging showed fusion and coalescence (Figure S2A). Fluorescence recovery after photobleaching (FRAP)^30^ demonstrated recovery of mVenus-YAP1-2α fluorescence within bleached puncta (Figures 1H, 1I, and S2B–S2G), with kinetics and apparent diffusion coefficients consistent with molecular exchange in relatively constrained nuclear assemblies. Although these properties do not alone establish liquid–liquid phase separation, they show that YAP1-2α puncta are dynamic and exhibit liquid-like behavior. Together, these findings identify YAP1-2α as a unique YAP1 splice isoform that selectively forms dynamic nuclear condensates, suggesting an isoform-specific mechanism of YAP1-dependent transcriptional regulation.

### YAP1-2α/TAZ/TEAD assembly is required for efficient condensate formation

Previous work showed that YAP1-2α and TAZ interact through coiled-coil leucine zippers to form heterodimers.^19^ Consistently, TAZ colocalized with YAP1-2α puncta in UWB1.289 cells (Figure 2A). To test whether this association contributes to condensate formation, we generated YAP1-2α(LZM), in which L311, L318, L325, L332, and L339 were replaced with alanine. This mutation reduced YAP1-2α-mediated TEAD reporter activation (Figure S3A). Proximity ligation assays (PLA) confirmed nuclear YAP1-2α–TAZ association; TAZ interacted with the YAP1-2α(S94A), which lacks TEAD binding,^4^ but did not efficiently with YAP1-2α(LZM), YAP1-2α(LZM/S94A), or YAP1-2γ (Figures S3B and S3C). Conversely, YAP1-2α, YAP1-2α(LZM), and YAP1-2γ retained TEAD interaction, whereas the S94A-containing variants did not (Figures S3B and S3D). These results separated the effects of disrupting YAP1-2α–TAZ heterodimerization from those of disrupting YAP1–TEAD binding.

**Figure 2.**
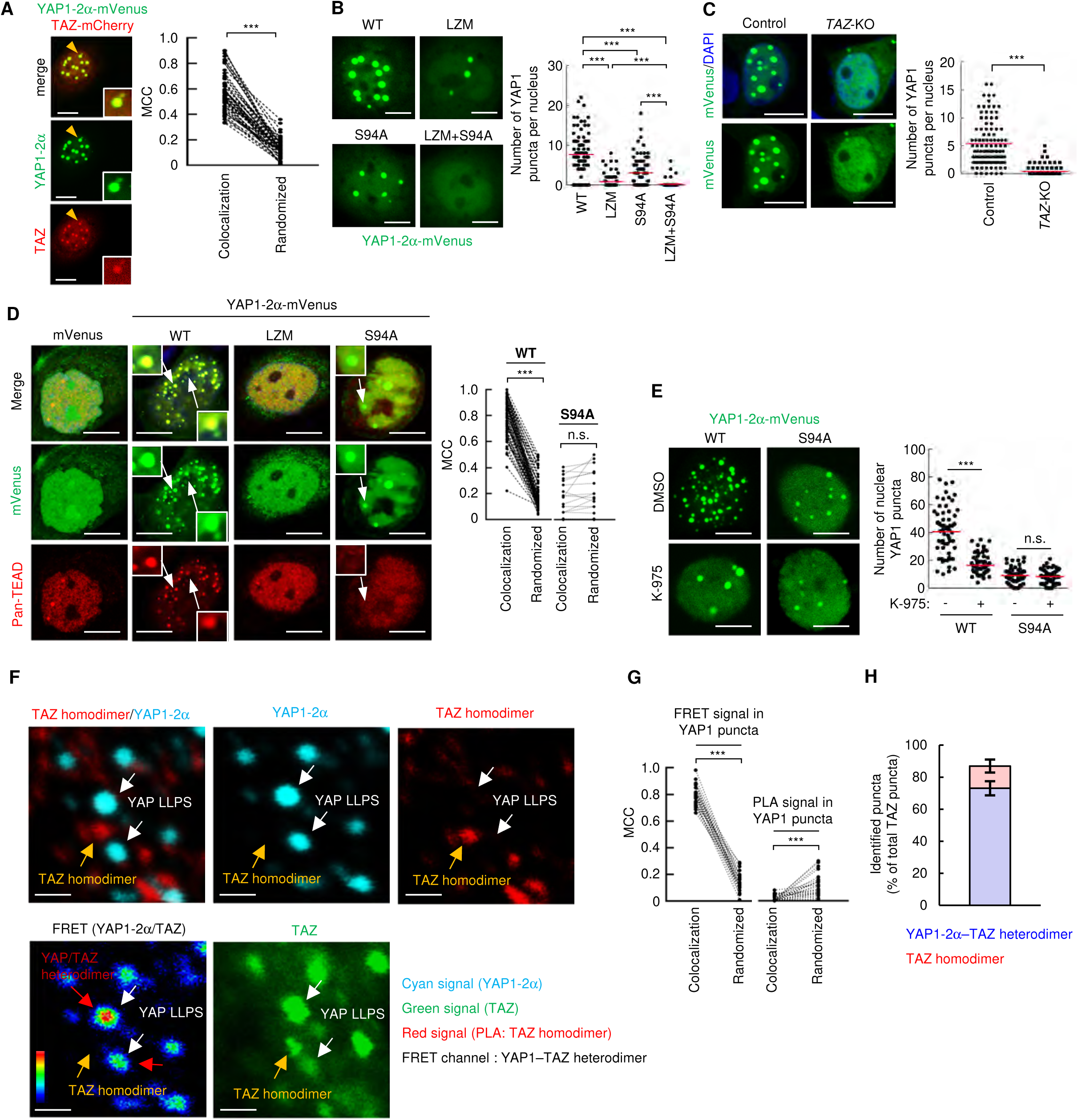
YAP1-2α–TAZ heterodimerization and TEAD association promote LLPS-like nuclear condensates. (**A**) Colocalization of YAP1-2α-mVenus and C-terminally mCherry-tagged TAZ (TAZ-mCherry) in UWB1.289 cells. Manders’ colocalization coefficient (MCC) was compared with randomized controls. (**B**) Nuclear puncta formed by the indicated mVenus-tagged YAP1-2α mutants in *YAP1*-KO UWB1.289 cells. See Figure S3F for protein expression. (**C**) YAP1-2α puncta in parental and *TAZ*-KO UWB1.289 cells. *TAZ* knockout is shown in Figure S3G. (**D**) Colocalization of the indicated YAP1-2α variants with pan-TEAD. Enlarged images show representative puncta (arrows). (**E**) Effects of K-975 treatment on puncta formed by YAP1-2α(WT) and YAP1-2α(S94A). (**F–H**) Combined FRET/PLA analysis of YAP1-2α/TAZ-associated and TAZ-homodimer-associated puncta. FRET signals indicate YAP1-2α/TAZ-associated puncta, whereas PLA signals indicate TAZ-homodimer-associated puncta (F). Arrows indicate representative puncta. Colocalization with YAP1-2α puncta (G) and the proportions of classified puncta (H) were quantified. See Figures S5A–S5D for the assay schematic, controls, co-immunoprecipitation, and low-magnification images. Data are mean ± SD; red lines indicate means. \*\*\**p* < 0.001; n.s., not significant. Paired *t* test (A, D, and G) or Mann–Whitney U test (B, C, and E). *n* = 60 cells (A), 75 cells per group (B), 100 cells per group (C), 75 and 17 cells for WT and S94A, respectively (D), 60 cells per group (E), 30 cells (G), and 434 puncta (H), from three independent biological replicates unless indicated otherwise. Scale bars, 10 μm. See also Figures S6–S8.

In *YAP1*-knockout UWB1.289 cells and MDA-MB-436 triple-negative breast cancer cells, YAP1-2α(LZM), YAP1-2α(S94A), and YAP1-2α(LZM/S94A) formed fewer nuclear condensates than wild-type YAP1-2α (Figures 2B and S3E–S3F). Endogenous TAZ deletion abolished YAP1-2α-induced condensate formation in both cell lines (Figures 2C and S3G–S3J). In parental UWB1.289 cells, wild-type YAP1-2α colocalized with TEAD proteins in nuclear puncta, whereas LZM and S94A mutants lacked comparable TEAD enrichment (Figure 2D). K-975,^31^ which disrupts YAP1/TAZ–TEAD interactions, abolished YAP1-2α-induced puncta without detectable cytotoxicity (Figures 2E, S4A and S4B).

Although YAP1-2α(S94A) formed a limited number of TAZ-containing puncta, these were less abundant than wild-type puncta, lacked significant Pol II colocalization, and showed reduced FRAP recovery, consistent with a less dynamic, predominantly solid-like state (Figures 2E and S4C–S4E). Thus, TEAD binding is dispensable for low-level YAP1-2α–TAZ clustering but contributes to efficient formation or stabilization of dynamic, Pol II-enriched nuclear condensates.

TAZ can form leucine zipper-dependent homodimeric condensates independently of YAP1 and TEAD.^32^ Accordingly, TAZ puncta formed similarly in parental and *YAP1*-knockout UWB1.289 cells (Figure S4F). FRET and PLA showed that YAP1-2α-induced condensates preferentially contained YAP1-2α–TAZ heterodimers, which accounted for approximately 73% of TAZ-containing condensates (Figures 2F–2H and S5A–S5D). Together, these findings indicate that stable YAP1-2α/TAZ/TEAD assembly is required for efficient formation of dynamic nuclear condensates.

### BRD4 multivalent interactions promote YAP1-2α/TAZ/TEAD-associated condensate formation

Leucine zipper-mediated YAP1-2α–TAZ heterodimerization alone could not explain the unique condensate-forming activity of YAP1-2α. YAP1-1α retains an intact leucine zipper and interacts with TAZ but did not form nuclear condensates under the same conditions (Figures 1A, 1B, and S6A). Unlike YAP1-1α, YAP1-2α contains two WW domains, suggesting a contribution of additional WW-domain-mediated interactions. Wild-type YAP1-2α/TAZ puncta colocalized with Pol II, whereas YAP1-2α(S94A)/TAZ puncta showed markedly reduced Pol II association (Figures S4C and S6B). FRAP analyses in *YAP1*-knockout UWB1.289 and MDA-MB-436 cells yielded apparent diffusion coefficients of 0.01–0.02 μm²/s, compatible with limited mobility within transcription-associated assemblies (Figures 1I, S2E, and S2G)^22,33^. YAP1-2α puncta did not colocalize with PML bodies, nuclear speckles, or nucleoli (Figures S6C and S6D).

BRD4 is enriched at active enhancers and super-enhancers and can contribute to transcription-associated condensate assembly through bromodomain-dependent recognition of acetylated chromatin.^22,34–36^ BRD4 contains LPxY (LPDY) and PPxY (PPTY) motifs,^17,37^ candidate ligands for YAP1 WW domains (Figure 3A). Co-immunoprecipitation and PLA confirmed YAP1-2α–BRD4 interactions in UWB1.289 and MDA-MB-436 cells; mutation of either motif weakened this interaction, whereas dual mutation abolished it (Figures 3B, 3C, S7A, and S7B). BRD4 depletion or competitive inhibition with a BRD4-derived PPTY peptide suppressed YAP1-2α/TAZ puncta (Figures 3D, 3E, and S7C–S7E). In BRD4-depleted cells, shRNA-resistant wild-type BRD4 restored condensate formation, whereas BRD4 proteins lacking either one or both PPxY/LPxY motifs did not (Figures 3F, S7F, and S7G).

**Figure 3.**
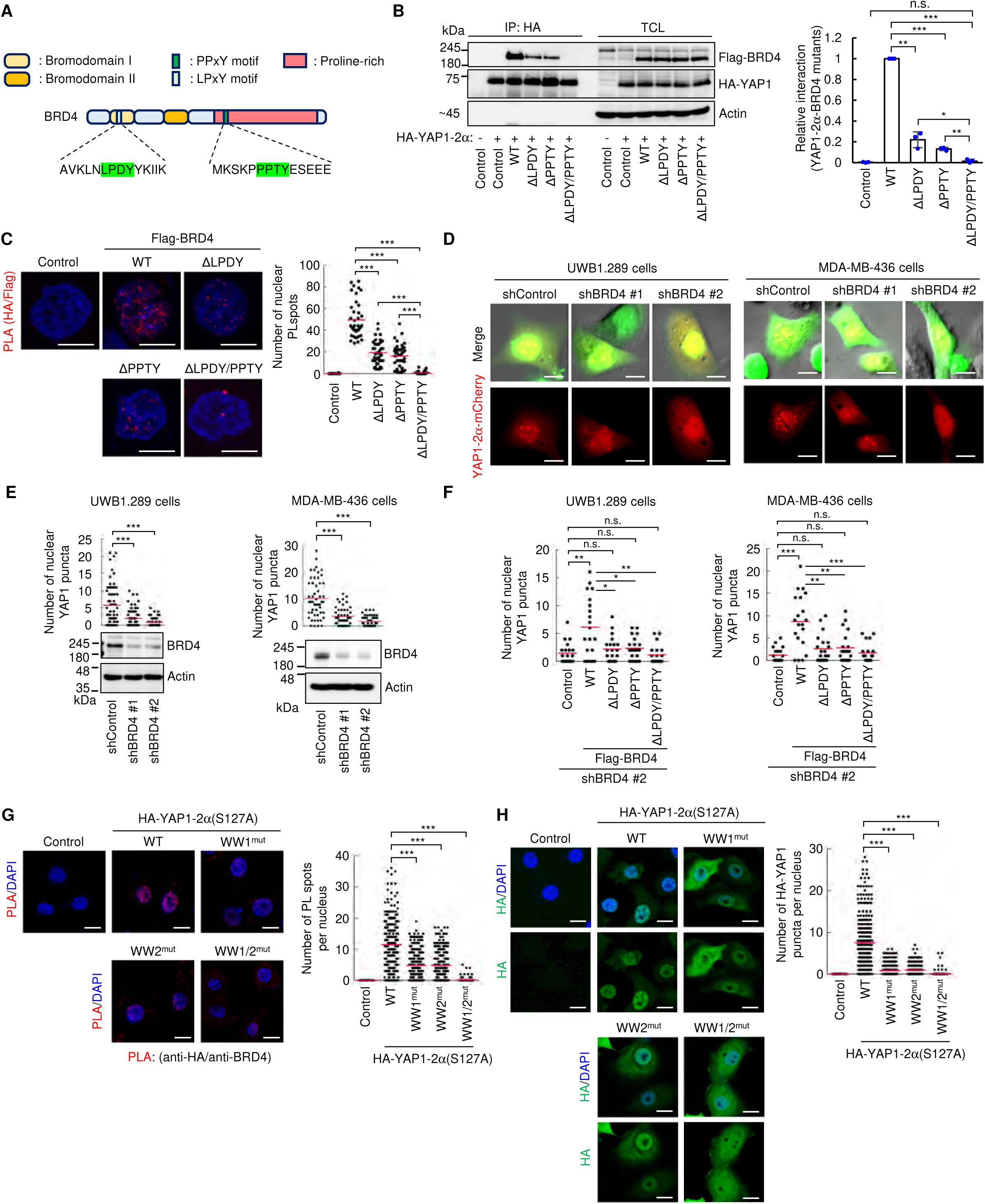
YAP1 WW domain-dependent association with BRD4 supports multivalent interactions in YAP1-2α/TAZ/TEAD LLPS-like condensates. (**A**) Schematic of BRD4 WW domain-binding motifs. (**B** and **C**) Association of HA-YAP1-2α with the indicated Flag-BRD4 mutants, assessed by co-immunoprecipitation in MDA-MB-436 cells (B) and PLA in UWB1.289 cells (C). Relative co-immunoprecipitation was normalized to wild-type BRD4. See Figures S7A and S7B for expression and additional co-immunoprecipitation data. (**D** and **E**) Effects of BRD4 knockdown on YAP1-2α-mCherry puncta in UWB1.289 and MDA-MB-436 cells. shBRD4-expressing cells are GFP positive; BRD4 depletion was confirmed by immunoblotting. (**F**) Rescue of shBRD4-mediated reductions in YAP1-2α puncta by the indicated shBRD4-resistant BRD4 constructs. (**G** and **H**) Effects of YAP1 WW-domain mutations on BRD4-associated PLA signals (G) and YAP1-2α nuclear puncta (H). See Figures S8C and S8D for mutant expression and co-immunoprecipitation with endogenous BRD4. Data are mean ± SD; red bars indicate means. \**p* < 0.05, \*\**p* < 0.01, \*\*\**p* < 0.001; n.s., not significant. Student’s *t* test (B) or Mann–Whitney U test (C and E–H). *n* = 3 biological replicates (B), 40 nuclei per group (C), 60 and 50 nuclei per group in UWB1.289 and MDA-MB-436 cells, respectively (E), 20 nuclei per group (F), and 300 nuclei per group (G and H), from three independent biological replicates unless indicated otherwise. Scale bars, 10 μm. See also Figure S6–S8.

To assess the two YAP1-2α WW domains, we introduced WW-disrupting mutations. Because YAP1-2α WW-domain mutants generated in the wild-type background accumulated predominantly in the cytoplasm,^18^ we introduced the WW-disrupting substitutions into the YAP1-2α(S127A) background^4^ to enable evaluation of nuclear condensate formation (Figures S8A and S8B). Disruption of either WW domain reduced BRD4 binding and nuclear condensate formation (Figures 3G, 3H, S8C, and S8D). Consistently, YAP1-1α bound BRD4 less efficiently than YAP1-2α and did not form nuclear puncta (Figures 1B and S8E–S8G). Together, these findings support a model in which multivalent interactions between the two YAP1-2α WW domains and the PPxY/LPxY motifs of BRD4 cooperate with YAP1-2α–TAZ heterodimerization and TEAD binding to promote transcription-associated nuclear condensates.

### YAP1-2α, TAZ, and TEAD converge at candidate super-enhancer-associated regulatory regions

Because YAP1-2α/TAZ/TEAD condensates were BRD4-associated and Pol II-enriched, we examined canonical TEAD target genes. YAP1-2α and YAP1-2γ similarly increased *CCN2* expression in UWB1.289 cells (Figures S9A and S9B), whereas YAP1-2α induced *CCN1* more strongly than other YAP1 isoforms (Figure 4A). TAZ deletion abolished the additional *CCN1* induction by YAP1-2α but had little effect on *CCN2* induction (Figures 4B, S9C, and S9D). YAP1-2α(LZM) retained partial *CCN1*-inducing activity in UWB1.289 cells (Figures 4C, and S9E), indicating that YAP1-2α/TEAD can activate *CCN1* independently of TAZ, whereas YAP1-2α/TAZ complex formation augments this response.

**Figure 4.**
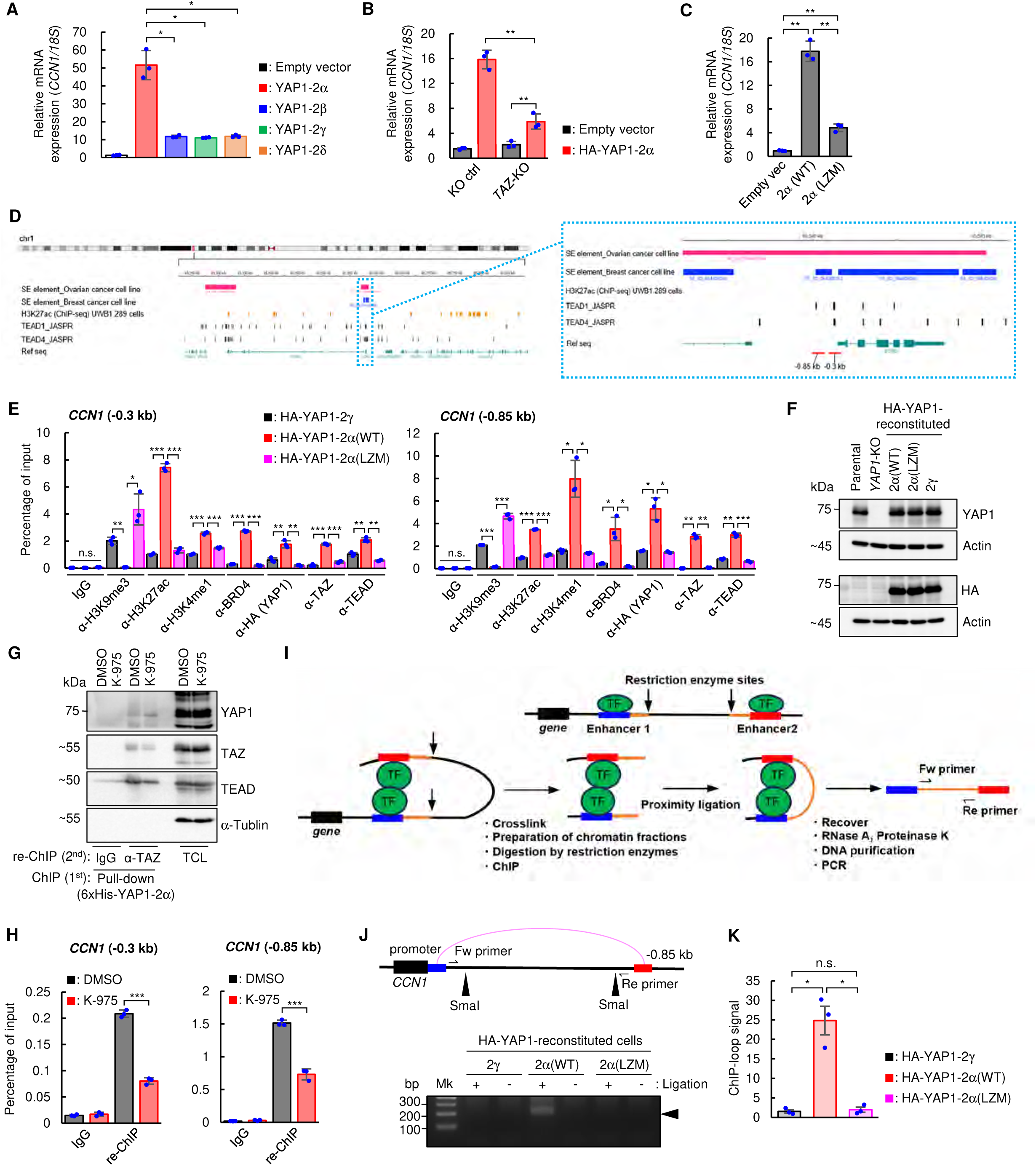
YAP1-2α, TAZ, TEAD, and BRD4 are enriched at selected candidate super-enhancer-associated regulatory regions. (**A–C**) *CCN1* expression in UWB1.289 cells expressing the indicated YAP1 isoforms (A), HA-YAP1-2α in parental or *TAZ*-KO cells (B), or HA-YAP1-2α(WT) and HA-YAP1-2α(LZM) (C). See Figures S9A, S9C, and S9E for protein expression. (**D**) Public H3K27ac datasets and predicted TEAD motifs nominate two candidate super-enhancer-associated regulatory regions upstream of *CCN1*. (**E** and **F**) CUT&RUN-qPCR analysis of chromatin marks and factor enrichment at the indicated candidate regions in UWB1.289 cells reconstituted with the indicated YAP1 variants. H3K9me3, H3K27ac, H3K4me1, BRD4, HA-YAP1, TAZ, and TEAD were examined; IgG served as a negative control (E). YAP1 replacement is shown in (F). (**G** and **H**) ChIP-re-ChIP analysis of YAP1-2α/TAZ/TEAD-associated chromatin at the candidate regions in 6×His-YAP1-2α-expressing *YAP1*-KO UWB1.289 cells. Effects of K-975 on YAP1-2α/TAZ/TEAD-associated protein complex formation (G) and chromatin enrichment (H) are shown. See Figure S9G for analyses at TEAD motif-negative loci. (**I–K**) ChIP-loop analysis of chromatin contact between the −0.85-kb candidate region and the *CCN1* promoter in YAP1-reconstituted UWB1.289 cells. Assay schematic (I). Representative PCR products amplified from ChIP-loop-purified DNA fragments (J). Quantification (K). Data are mean ± SD (A–C, F, and H) or mean ± SEM (K). \**p* < 0.05, \*\**p* < 0.01, \*\*\**p* < 0.001; n.s., not significant; Welch’s *t* test. *n* = 3 technical replicates (A–C, F, and H) or three biological replicates (K). Duplicate aliquots of the same cell lysates were resolved on the same SDS-PAGE gel, and the transferred membrane was cut to allow separate detection of endogenous YAP1 and the HA-tagged protein (F). See also Figure S9.

Using SEdb3.0 and public H3K27ac datasets^38,39^, we identified TEAD motif-containing regulatory regions upstream of *CCN1*, but not *CCN2*, predicted as candidate super-enhancer-associated elements in serous ovarian and breast cancer cells (Figures 4D and S9F). Because these annotations were derived from external datasets, we refer to these elements as “candidate super-enhancer-associated regions” and do not infer a genome-wide super-enhancer landscape in UWB1.289 cells.

In *YAP1*-knockout UWB1.289 cells reconstituted with YAP1 isoforms, wild-type YAP1-2α increased BRD4, H3K27ac, H3K4me1, YAP1, TAZ, and TEAD enrichment at regions 0.3 and 0.85 kb upstream of the *CCN1* transcription start site relative to YAP1-2γ or YAP1-2α(LZM) (Figures 4E and 4F). Conversely, H3K9me3 was more enriched with YAP1-2γ or YAP1-2α(LZM), consistent with a more active chromatin state in the presence of intact YAP1-2α/TAZ assembly. Public H3K27ac ChIP-seq data^40^ showed limited enrichment at these elements in parental UWB1.289 cells (Figure 4D).

ChIP-re-ChIP demonstrated TEAD-dependent enrichment of the YAP1-2α/TAZ complex at the candidate *CCN1* regulatory regions (Figures 4G, 4H, and S9G). ChIP-loop assays showed greater association between the region 0.85 kb upstream of *CCN1* and the *CCN1* promoter in YAP1-2α-reconstituted cells than in YAP1-2α(LZM)- or YAP1-2γ-reconstituted cells (Figures 4I–4K). JQ1^41^ suppressed YAP1-2α-induced *CCN1* expression (Figure S9H). Together, these findings indicate that YAP1-2α/TAZ/TEAD complexes and BRD4 converge at candidate super-enhancer-associated regulatory regions upstream of *CCN1* and promote their association with the *CCN1* promoter.

### IDR-dependent YAP1-2α condensates promote EMT- and stemness-associated transcriptional programs

To identify transcriptional signatures associated with *YAP1* splice variation in human tumors, we analyzed TCGA HGSOC RNA-seq and SpliceSeq data.^42–45^ Tumors were classified as *YAP1-1/2α–, -β–, -γ–, or -δ–*dominant according to percent-spliced-in values for exon 5.2 and exon 6. Compared with *YAP1-1/2γ–*dominant tumors, *YAP1-1/2α–*dominant tumors showed 754 upregulated and 146 downregulated genes (Figure S10A). Hallmark gene set analysis linked the upregulated genes to EMT and angiogenesis (Figure 5A). *SPARC, ALDH1A3, and SNAI2,* which have been implicated in EMT, stemness, invasion, and drug resistance,^8,46–52^ were more highly expressed in *YAP1-1/2α*–dominant tumors despite similar *YAP1* and *TAZ* mRNA levels across groups (Figure S10B).

**Figure 5.**
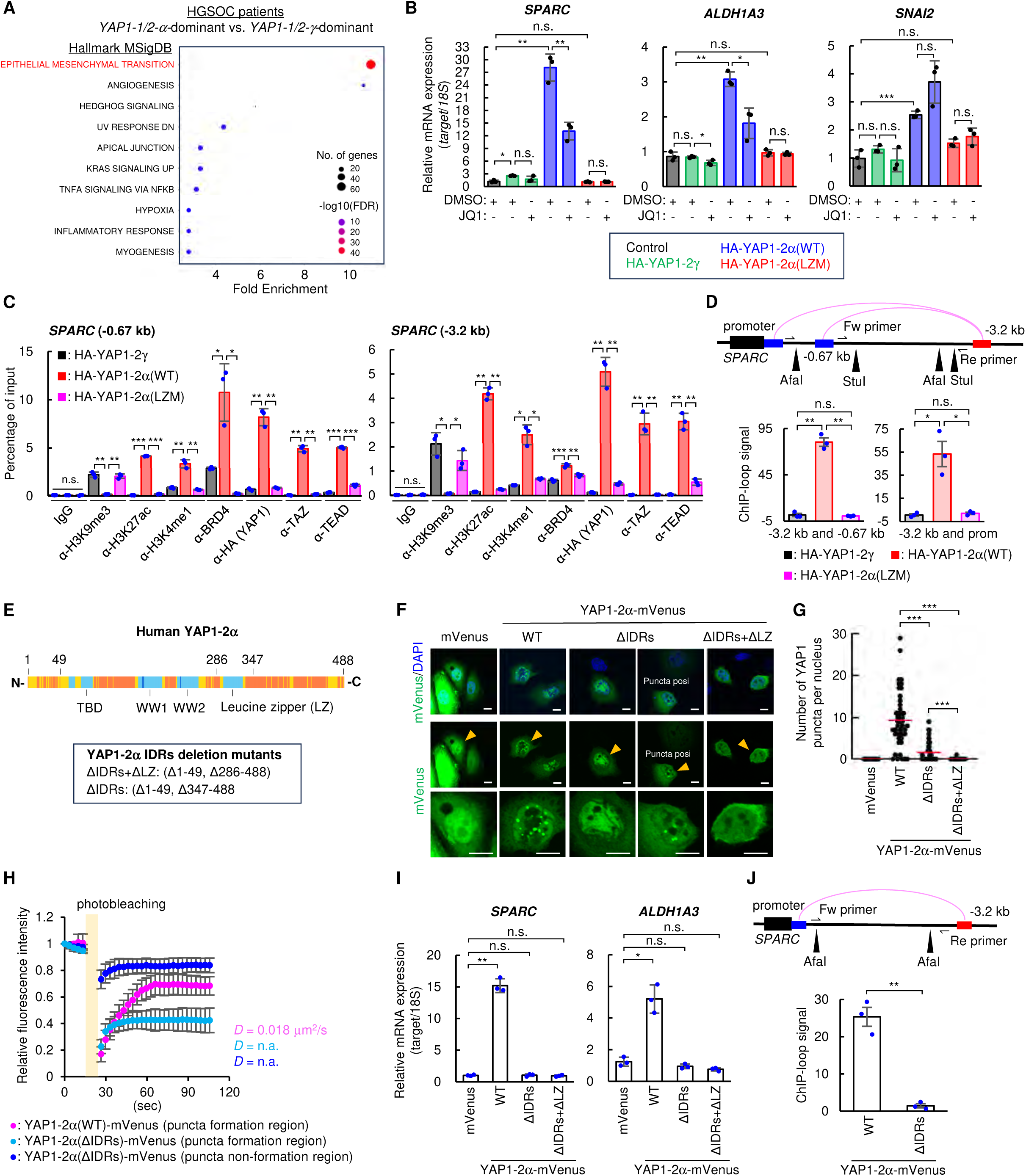
IDR integrity contributes to YAP1-2α-associated nuclear puncta and expression of EMT- and stemness-associated genes. (**A**) Gene set enrichment analysis of genes upregulated in YAP1-1/2α-dominant versus YAP1-1/2γ–dominant HGSOC. Circle size indicates gene-set size, and color indicates −log10(FDR). See Figure S10A for differential-expression analysis. (**B**) Expression of *SPARC*, *ALDH1A3*, and *SNAI2* in UWB1.289 cells expressing the indicated YAP1 variants and treated with JQ1 or DMSO. See Figure S10C for protein expression. (**C**) CUT&RUN-qPCR analysis of chromatin marks and factor enrichment at candidate super-enhancer-associated regions linked to SPARC in UWB1.289 cells reconstituted with the indicated YAP1 variants. H3K9me3, H3K27ac, H3K4me1, BRD4, HA-YAP1, TAZ, and TEAD were examined; IgG served as a negative control. See Figure S11A for the analyzed regions. (**D**) ChIP-loop analysis of chromatin contacts among the indicated SPARC candidate regions and the promoter in YAP1-reconstituted UWB1.289 cells. (**E**) Predicted disorder of YAP1-2α based on AlphaFold pLDDT scores. (**F–H**) Effects of N- and C-terminal IDR deletions on YAP1-2α-mVenus puncta (F and G) and FRAP recovery (H) in *YAP1*-KO UWB1.289 cells. See Figure S12B for representative whole-nucleus images. (**I** and **J**) Effects of the indicated YAP1-2α constructs on *SPARC* and *ALDH1A3* expression (I) and contact between the −3.2-kb *SPARC* candidate region and promoter (J). Data are mean ± SD (B, C, H, and I) or mean ± SEM (D and J); red bars indicate means. \**p* < 0.05, \*\**p* < 0.01, \*\*\**p* < 0.001; n.s., not significant; Mann–Whitney U test (G) or Welch’s *t* test (B–D and I–J). n = 3 technical replicates (B, C, and I), 45 cells (G), 10 cells per group (H), or three biological replicates (D and J). Scale bars, 10 μm (F). See also Figure S10–S12.

Given the selective ability of YAP1-2α to form nuclear puncta among the eight isoforms tested, we next examined whether YAP1-2α promotes expression of these genes in UWB1.289 cells. YAP1-2α increased *SPARC, ALDH1A3,* and *SNAI2* expression more strongly than YAP1-2γ or YAP1-2α(LZM) (Figures 5B and S10C). JQ1, a BRD4-derived PPTY peptide, or TAZ deletion reduced YAP1-2α condensates and attenuated *SPARC* and *ALDH1A3* induction (Figures S7C–S7E and S10D–S10F), whereas TAZ overexpression alone was insufficient to activate these genes (Figures S10G–S10H). These results support a requirement for coordinated YAP1-2α/TAZ/BRD4 activity rather than increased TAZ abundance alone. Candidate regulatory regions upstream of *SPARC* and *ALDH1A3* showed greater enrichment of H3K27ac, H3K4me1, BRD4, TAZ, TEAD, and YAP1 in YAP1-2α(WT)-reconstituted cells than in cells reconstituted with YAP1-2γ or YAP1-2α(LZM) (Figures 5C, S11A, and S11B). ChIP-loop assays further revealed increased interactions between these regions and their cognate promoters (Figures 5D and S11C–S11E).

To examine the contribution of IDR-dependent assembly to YAP1-2α-mediated gene induction, we generated two deletion mutants guided by the YAP1-2α sequence and AlphaFold-derived pLDDT scores.^53^ YAP1-2α(ΔIDRs) lacked residues 1–49 and 347–488 but retained the leucine zipper (residues 286–331), whereas YAP1-2α(ΔIDRs+ΔLZ) lacked residues 1–49 and 286–488, including the leucine zipper (Figure 5E). Both mutants were markedly impaired in condensate formation (Figures 5F, 5G, and S12A). Residual puncta formed by YAP1-2α(ΔIDRs) exhibited reduced fluorescence recovery after photobleaching, consistent with reduced molecular exchange (Figures 5H and S12B). Although YAP1-2α(ΔIDRs) retained TEAD-dependent reporter activity and BRD4 binding (Figures S12C–S12E), it was less effective at inducing *SPARC* and *ALDH1A3* expression and failed to maintain the increased interactions between the candidate regulatory regions and their cognate promoters (Figures 5I, 5J, and S12F–S12H). Together, these findings support a role for IDR-dependent YAP1-2α assembly in promoting *SPARC* and *ALDH1A3* expression and maintaining interactions between the candidate regulatory regions and their cognate promoters.

### YAP1-2α promotes malignant phenotypes and intraperitoneal tumor progression

Because the YAP1-2α-associated transcriptional program included EMT- and stemness-related genes, we examined malignant cellular phenotypes.^54–57^ In two-dimensional culture, YAP1-2α- and YAP1-2γ-expressing UWB1.289 cells showed similarly modest increases in proliferation relative to controls (Figure S13A). In contrast, YAP1-2α preferentially enhanced sphere formation in three-dimensional culture (Figure S13A). This phenotype was reduced by K-975, JQ1, or YAP1-2α leucine zipper mutation (Figures 6A and 6B), linking sphere formation to the YAP1-2α/TAZ/TEAD/BRD4-associated condensate state. YAP1-2α also enhanced chemotactic migration, extracellular-matrix invasion, and ferroptosis resistance (Figures 6C, 6D, and S13B). Ferrostatin-1 rescued the cytotoxic effect of the ferroptosis inducer in parental cells (Figure S13C), supporting ferroptosis as the relevant mode of cell death. TEAD or BRD4 inhibition and leucine zipper mutation attenuated these phenotypes. Wild-type YAP1-2α also increased resistance to trametinib, sorafenib, cisplatin, and olaparib; these effects were reduced by K-975, JQ1, or leucine zipper mutation (Figures 6E and S13D). Thus, YAP1-2α promotes malignant phenotypes through an intact YAP1-2α–TAZ interaction and TEAD- and BRD4-dependent activity.

**Figure 6.**
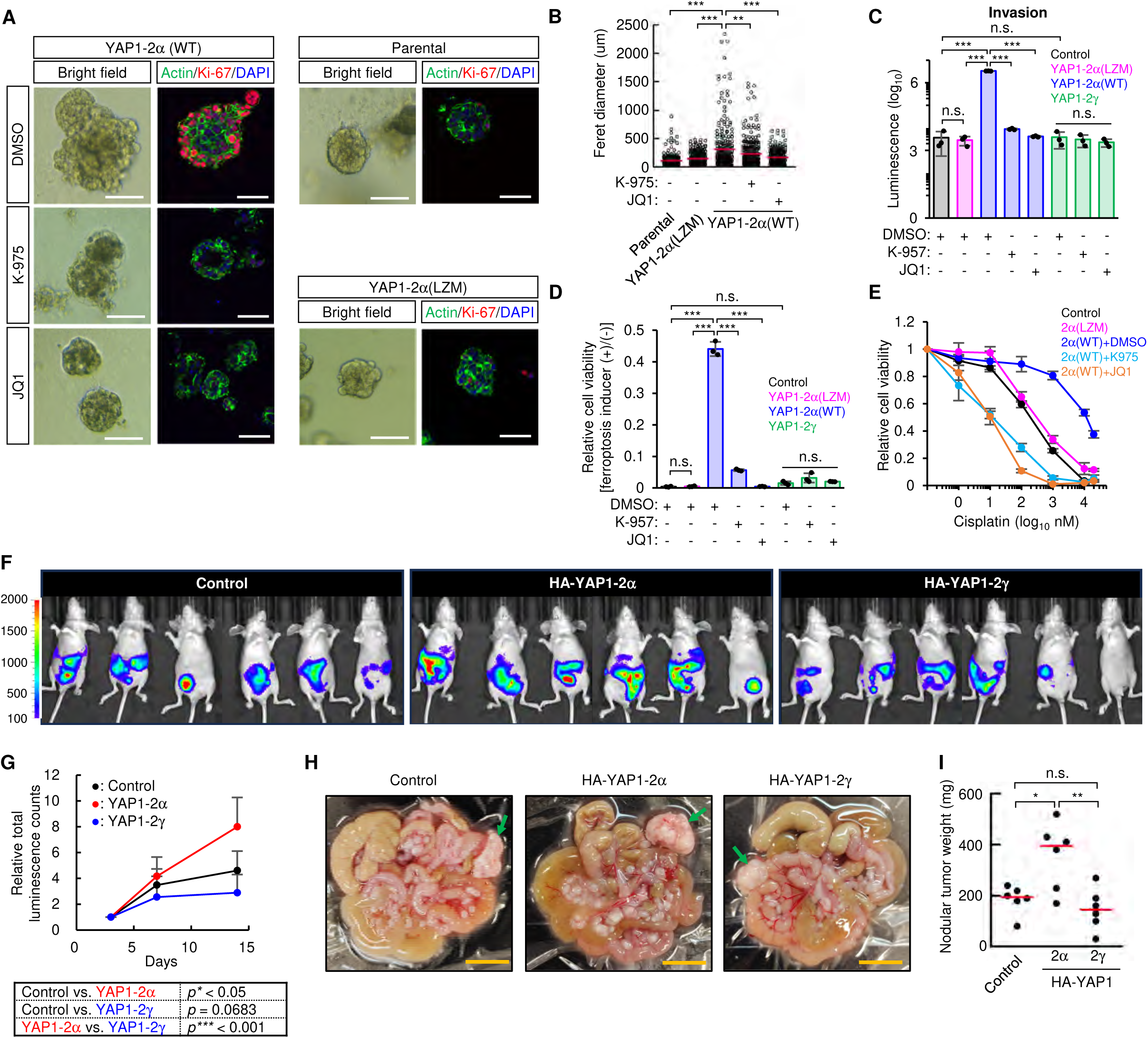
YAP1-2α-associated LLPS-like condensates promote malignant phenotypes and intraperitoneal tumor growth. (**A** and **B**) Effects of K-975 and JQ1 on spheroids formed by control, YAP1-2α(WT)-expressing UWB1.289 cells. YAP1-2α(LZM) was used as a nuclear condensate-defective control. Representative bright-field and Ki-67 images (A) and Feret diameters (B) are shown. (**C–E**) Effects of YAP1-2α(WT), YAP1-2α(LZM), and YAP1-2γ expression, with K-975, JQ1, or DMSO treatment, on matrix invasion (C), ferroptosis-associated cell death (D), and cisplatin sensitivity (E) in UWB1.289 cells. See Figure S13C for ferrostatin-1 rescue. (**F–I**) Intraperitoneal xenograft analysis of luciferase-expressing MKN45 cells transduced with the indicated HA-YAP1 isoforms. Representative IVIS images at day 14 (F). Longitudinal total intraperitoneal bioluminescence normalized to day 3 (G). Representative peritoneal and omental tumor nodules collected at day 14; arrows indicate omental nodules (H). Weights of resected tumor nodules (I). See Figures S14B–S14E for construct expression, luciferase activity, and tumor qPCR analyses. Data are mean ± SD; red bars indicate means. \**p* < 0.05, \*\**p* < 0.01, \*\*\**p* < 0.001; n.s., not significant; Student’s *t* test (B) or Welch’s *t* test (C–E, G, and I). *n* > 230 spheroids per group (B), three technical replicates (C–E), or six mice per group (G and I). Scale bars, 100 μm (A) and 1 cm (H). See also Figures S13 and S14.

We next assessed the in vivo consequences of YAP1 isoform expression using firefly luciferase-expressing MKN45 (MKN45-Fluc) gastric cancer cells,^58^ because UWB1.289 cells do not reproducibly engraft in nude mice.^59^ YAP1-2α, but not YAP1-2γ, formed nuclear condensates in MKN45-Fluc cells (Figure S14A), indicating that the isoform-specific phenotype was retained in this gastric cancer. MKN45-Fluc cells that stably express HA-YAP1-2α or HA-YAP1-2γ at comparable levels and exhibit comparable luciferase activities were injected intraperitoneally into nude mice (Figures S14B and S14C). Mice intraperitoneally inoculated with MKN45-Fluc cells expressing HA-YAP1-2α developed greater longitudinal bioluminescence signals and heavier intestinal tumor nodules than mice inoculated with parental or HA-YAP1-2γ-expressing cells (Figures 6F–6I). The recovered tumors retained ectopic YAP1 expression and showed increased *SPARC* and *ALDH1A3* expression in the YAP1-2α group (Figures S14D and S14E). These findings show that elevated YAP1-2α is sufficient to promote intraperitoneal tumor progression in this model, but do not establish an in vivo requirement for endogenous YAP1-2α.

### Endogenous *YAP1* exon 6 exclusion promotes a YAP1-2α-associated condensate and malignant program

We next examined whether modulation of endogenous *YAP1* pre-mRNA splicing promotes a YAP1-2α-associated condensate state and malignant phenotypes. Exon 6 exclusion can generate YAP1-2α and/or YAP1-2β, whereas exon 6 inclusion generates YAP1-2γ and/or YAP1-2δ. We therefore used an antisense oligonucleotide targeting *YAP1* exon 6 (YAP1-E6 ASO)^60^ to promote exon 6 exclusion (Figure S15A). In parental UWB1.289 cells, YAP1-2γ was the predominant YAP1-2 isoform, whereas exon 6-lacking isoforms were scarce (Figures S1A and S1F). YAP1-E6 ASO markedly increased exon 6-lacking transcripts without altering the total abundance of *YAP1-1* and *YAP1-2* transcripts (Figures S15B and S15C). Sequencing of the splice products, together with the substantially higher expression level (approximately 10-fold) of YAP1-2 than YAP1-1 in UWB1.289 cells (Figure S1A: lane 1), indicated that this intervention predominantly favored a YAP1-2α-associated splicing program. YAP1-E6 ASO also induced endogenous nuclear YAP1 condensates containing BRD4, increased YAP1–TAZ heterodimerization, enhanced TAZ-dependent expression of *SPARC* and *ALDH1A3*. In parallel, exon 6 exclusion promoted TAZ-dependent sphere formation, invasion, and therapeutic resistance (Figures 7A and S15D–S15L). Thus, modulation of endogenous *YAP1* splicing toward exon 6 exclusion was sufficient to induce a BRD4-associated, YAP1-2α-linked condensate state and associated malignant phenotypes without ectopic YAP1-2α expression.

**Figure 7.**
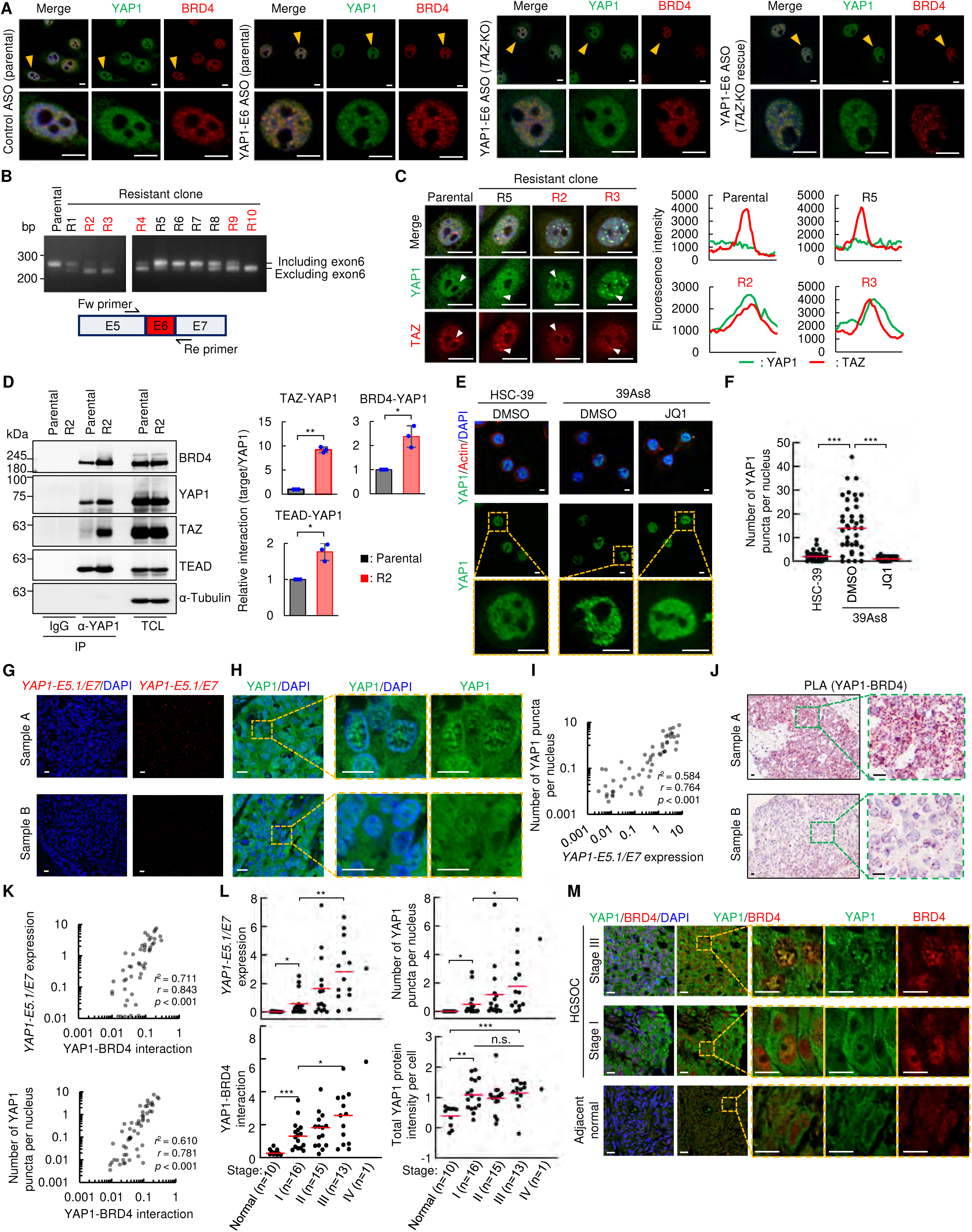
Endogenous YAP1 exon 6 exclusion and a YAP1-2α-linked nuclear punctate state are associated with stress adaptation and advanced HGSOC. (**A**) Effects of E6-ASO-mediated *YAP1* exon 6 exclusion on endogenous nuclear YAP1 puncta in parental, *TAZ*-KO, and TAZ-rescued UWB1.289 cells. Scrambled ASO was used as a negative control. See Figures S15A–S15L for ASO validation and related analyses. (**B–D**) *YAP1* exon 6 splicing in parental UWB1.289 cells and cisplatin-resistant clones (R1–R10) (B), endogenous YAP1 or TAZ puncta, and YAP1 association with TAZ (C), TEAD, and BRD4 (D) in R2 cells. Scanning fluorescence intensity of YAP1 (red) or TAZ (green) within a single punctum indicated by the white arrow in the left panels is shown in the right panels (C). See Figures S16A–S16I for related analyses. (**E** and **F**) Effect of JQ1 on endogenous nuclear YAP1 puncta (E) and their quantification (F) in 39As8 cells, an HSC-39-derived highly aggressive subline. See Figures S17A–S17C for related analyses. (**G–L**) Analysis of HGSOC clinical specimens. H&E staining of samples A and B is shown in Figure S18C. Representative BaseScope images show high expression of *YAP1-E5.1/E7* transcripts in sample A (upper; signal score 3) and low expression in sample B (lower; signal score 0). Red spots indicate BaseScope signals (G). Representative images show endogenous nuclear YAP1 puncta (green) in sample A (upper) but not in sample B (lower) (H). Correlation between *YAP1-E5.1/E7* transcript levels and nuclear YAP1 puncta formation (I). Representative PLA images show endogenous YAP1–BRD4 proximity in samples A and B (J). Correlation of YAP1–BRD4 PLA signals with *YAP1-E5.1/E7* transcript levels (top) and nuclear YAP1 puncta (bottom) (K). Associations of FIGO stage with *YAP1-E5.1/E7* transcript signals, nuclear YAP1 puncta, YAP1–BRD4 PLA signals, and total YAP1 intensity in the discovery cohort (L). (**M**) Representative images of YAP1 and BRD4 colocalization in adjacent normal, stage I, and stage III HGSOC clinical specimens. See Figures S19A–S19H for related analyses. Data are mean ± SEM; red bars indicate means. \**p* < 0.05, \*\**p* < 0.01, \*\*\**p* < 0.001; n.s., not significant; Welch’s *t* test (D and L) or Mann–Whitney U test (F). Pearson’s correlation coefficients are shown in (I and K). *n* = three biological replicates (D) or 40 cells per group (F). Scale bars, 10 μm. See also Figures S15–S19.

To test whether a YAP1 exon 6-excluded, YAP1-2α-associated nuclear state can arise during stress adaptation, we generated independently derived cisplatin-resistant UWB1.289 clones (R1 to R10) by repeated exposure to sublethal cisplatin. Of these, five showed increased exon 6 exclusion, whereas the remainder showed no detectable change (Figures 7B and S16A). In exon 6-excluded R2 and R3, but not in exon 6-included R5, K-975 or JQ1 restored cisplatin sensitivity (Figures S16B). *YAP1* depletion also sensitized R2 and R3 cells, whereas shRNA-resistant YAP1-2α, but not YAP1-2γ, restored resistance (Figures S16C and S16D). R2 and R3 contained abundant endogenous YAP1 condensates colocalized with TAZ; YAP1 association with TAZ, TEAD, and BRD4 was increased in R2 cells (Figures 7C and 7D). *TAZ* knockout reduced YAP1 condensates and restored cisplatin sensitivity in R2 cells, whereas TAZ overexpression alone did not alter parental-cell sensitivity (Figures S16E–S16I).

We also examined the metastatic subclone 39As8, derived from HSC-39 diffuse-type gastric cancer cells after in vivo peritoneal selection.^61^ Relative to parental HSC-39 cells, the peritoneal dissemination-derived subclone 39As8 showed increased exon 6 exclusion, BRD4-associated nuclear YAP1 condensates, and reduced cisplatin sensitivity; both K-975 and JQ1 restored cisplatin sensitivity (Figures 7E, 7F, and S17A–S17C). Together, these findings indicate that a *YAP1* splicing shift can contribute to stress adaptation and drug resistance in selected cancer-cell populations, rather than representing a universal resistance mechanism.

### YAP1-2α expression and YAP1/TAZ/BRD4-associated condensates correlate with HGSOC progression

Finally, we assessed the clinical relevance of YAP1-2α-associated condensates in human HGSOC specimens using a BaseScope probe targeting the *YAP1* exon 5.1–exon 7 junction. We collectively refer to *YAP1* mRNAs containing this junction as *YAP1-E5.1/E7*; these can encode YAP1-1α or YAP1-2α. Probe specificity was validated in *YAP1*-knockout UWB1.289 cells expressing individual YAP1 isoforms (Figure S18A). Although *YAP1-E5.1/E7* includes transcripts encoding both isoforms, only YAP1-2α formed discernible nuclear YAP1 puncta in our isoform-reconstitution system.

In HGSOC specimens, *YAP1-E5.1/E7* abundance correlated with nuclear YAP1 condensates (Figures 7G–7I and S18B–S18C). YAP1–BRD4 PLA signals correlated with both *YAP1-E5.1/E7* abundance and nuclear YAP1 condensates (Figures 7J, 7K, and S18D). *YAP1-E5.1/E7* levels, nuclear YAP1 condensates, and YAP1–BRD4 PLA signals were higher in FIGO stage III than in stage I tumors, whereas total YAP1 protein intensity was not associated with stage (Figure 7L). Thus, the nuclear punctate YAP1 state and its association with BRD4, rather than total YAP1 abundance alone, were linked to advanced disease.

In TCGA HGSOC data, lower inclusion of exons 5.2 and 6 was associated with poorer overall survival (Figure S18E). However, neither exon 5.2 nor exon 6 inclusion was an independent prognostic factor in multivariable Cox models adjusted for age, FIGO stage, and YAP1 expression (Figure S18F).

In an independent HGSOC cohort, associations among *YAP1-E5.1/E7* abundance, YAP1–BRD4 interaction, nuclear YAP1 condensates, and FIGO stage were reproduced (Figures S19A–S19G). BRD4 and TAZ colocalized with nuclear YAP1 condensates in tumor tissues (Figures 7M, S19C, and S19H). Collectively, these correlative analyses link *YAP1-E5.1/E7* expression and YAP1-2α/TAZ/BRD4-associated nuclear condensates with advanced HGSOC.

## DISCUSSION

We identify YAP1-2α as a YAP1 splice isoform with a distinctive capacity to form dynamic, condensate-like nuclear puncta and to support a BRD4- and TEAD-associated transcriptional state linked to malignant phenotypes. At representative candidate super-enhancer-associated regulatory regions, YAP1-2α, TAZ, TEAD, and BRD4 co-enriched for active chromatin features and showed increased local chromatin interactions. These findings support an alternative splicing-dependent model in which YAP1-2α organizes a transcriptionally active nuclear state that promotes EMT, stemness, invasion, sphere formation, and therapeutic resistance. Consistent with this model, a recent comprehensive analysis of YAP1 splice isoforms reported that YAP1-2α has transcriptional properties distinct from those of other YAP1 isoforms and enhances expression of TEAD target genes.^16^

Our findings provide a mechanistic framework for functional diversity among the eight YAP1 splice isoforms in cancer cells. In a comprehensive isoform screen in high-grade serous ovarian cancer (HGSOC) cells, YAP1-2α was the only isoform that consistently formed prominent nuclear puncta under identical experimental conditions. YAP1-1α, despite retaining the leucine zipper required for TAZ interaction, did not form YAP1 nuclear puncta. A plausible explanation is that the additional WW domain in YAP1-2α increases interaction valency with TAZ, BRD4, and other nuclear partners, thereby favoring the formation or stabilization of a BRD4-associated nuclear assembly. This interpretation is supported by impaired BRD4 interaction and puncta formation after disruption of the WW2 domain in YAP1-2α or the relevant WW-domain-binding motifs in BRD4. Thus, the second WW domain of YAP1-2α is a key multivalent feature that promotes formation of a BRD4-associated nuclear assembly.

Heterodimerization with TAZ, TEAD engagement, and IDR-mediated multivalency further cooperate to generate the transcriptionally competent YAP1-2α state. The YAP1-2α(S94A) mutant, which cannot bind TEAD, retained limited TAZ-containing nuclear clustering, indicating that TEAD engagement is not absolutely required for the initial clustering propensity of YAP1-2α. The persistence of residual S94A puncta further suggests that additional TEAD-independent mechanisms may contribute to YAP1-2α clustering, although the molecular basis and functional significance of these structures remain unclear and warrant further investigation. Nevertheless, compared with wild-type YAP1-2α, the S94A mutant formed fewer puncta, showed markedly reduced colocalization with RNA polymerase II, and displayed slower, less complete fluorescence recovery after photobleaching. These findings distinguish low-level, TEAD-independent YAP1-2α–TAZ clustering from the BRD4- and RNA polymerase II-enriched puncta formed by wild-type YAP1-2α. Experiments with YAP1-2α IDR-deletion mutants further support a role for multivalent, condensate-associated assembly in YAP1-2α-dependent transcriptional regulation. A YAP1-2α mutant, in which a substantial portion of IDRs were deleted, retained interactions with BRD4, TAZ, and TEAD and preserved canonical TEAD-dependent activity, but formed fewer and less dynamic puncta and failed to efficiently activate target genes linked to candidate super-enhancer-associated regulatory regions, including *SPARC* and *ALDH1A3*, or to support the local chromatin interactions examined. Conversely, leucine-zipper disruption abolished TAZ binding and strongly impaired puncta formation, chromatin association, transcriptional activation, and malignant phenotypes. Together, these data support a model in which YAP1-2α–TAZ heterodimerization, TEAD engagement, BRD4 interaction, and IDR-mediated multivalency cooperate to promote YAP1-2α nuclear organization and transcriptional output.

The relevance of the YAP1-2α-associated punctate program beyond HGSOC is supported by findings in triple-negative breast cancer (TNBC) and gastric cancer models. The puncta-formation patterns of wild-type YAP1-2α and its leucine-zipper- and TEAD-binding-defective mutants were recapitulated in TNBC cells. These findings extend condensate-associated phenotypes beyond HGSOC cells. Although both the HGSOC and TNBC cell lines used for the mechanistic experiments harbor *BRCA1* defects,^62^ elevated expression of YAP1-2α, but not YAP1-2γ, also induced nuclear YAP1 puncta in *BRCA*-proficient gastric cancer cells. Moreover, an in vivo-selected *BRCA*-proficient gastric cancer subclone with increased potential for intraperitoneal expansion^61^ exhibited exon 6 skipping, formed YAP1-associated nuclear puncta, and showed reduced cisplatin sensitivity. Complementing these observations, elevated YAP1-2α expression in another *BRCA*-proficient gastric cancer cell line potentiated peritoneal dissemination and increased intraperitoneal tumor burden relative to parental cells and a YAP1-2γ-expressing isoform control. Together, these findings demonstrate the pro-tumorigenic capacity of YAP1-2α in a *BRCA*-proficient gastric cancer model and extend the potential relevance of this mechanism beyond *BRCA*-deficient contexts. Nevertheless, these findings do not directly establish the applicability of this mechanism to *BRCA*-proficient HGSOC or isolate the contribution of *BRCA* status from other differences between the models. Validation in *BRCA*-proficient HGSOC and genetically matched models will be needed to address these questions.

The occurrence of exon 6 skipping and YAP1-associated nuclear puncta in the peritoneally selected gastric cancer subclone raised the idea that this program may be linked to cancer cell adaptation to circumstantial/microenvironmental changes/pressure. Consistent with this possibility, our ASO-based experiments demonstrated that enforced exon 6 skipping of *YAP1* pre-mRNA in HGSOC cells preferentially increased YAP1-2α expression and induced an endogenous YAP1-associated punctate state. This splicing switch also increased BRD4-associated nuclear puncta, enhanced YAP1–TAZ heterodimerization, stimulated YAP1/TAZ/TEAD-dependent expression of candidate target genes, and promoted cell-intrinsic malignant phenotypes. The cisplatin-resistant clone panel further supports a role for the *YAP1* pre-mRNA splicing switch in therapeutic adaptation. A subset of cisplatin-resistant clones displayed increased exon 6 skipping and the emergence of nuclear YAP puncta; treatment with the TEAD inhibitor K-975 or the BET/BRD4 inhibitor JQ1 restored cisplatin sensitivity in these clones. In contrast, a resistant clone that retained exon 6 inclusion showed neither YAP1 puncta formation nor sensitivity to TEAD inhibition under the conditions tested.

The emergence of *YAP1* exon 6 skipping in selected cisplatin-resistant clones and in peritoneally selected gastric cancer cells suggests that the YAP1-2α-associated punctate state may be an adaptive strategy cancer cells use to withstand unfavorable conditions. By promoting a BRD4- and TEAD-associated transcriptional program linked to stemness, invasion, and therapeutic resistance, this state may enable a subset of cancer cells to persist under adverse, stressed conditions. We propose that a stress-associated splicing switch toward YAP1-2α increases its availability and favors puncta formation, rather than assuming that detectable puncta are constitutively present under basal conditions. Whether this switch is required for stress-associated puncta formation remains to be directly established. For instance, in the presence of DNA-damaging chemotherapeutic agents, DNA-damage-sensing signaling and stress-responsive RNA splicing regulators are plausible upstream mechanisms that could favor *YAP1* exon 6 skipping and YAP1-2α production. However, the upstream signaling pathways and specific splicing factors responsible for *YAP1* exon 6 skipping remain unknown. Our study addressed the biological consequences of isoform switching rather than the regulatory mechanisms that initiate it. Identifying these regulators will be important for understanding how YAP1 isoform switching is associated with adaptation to therapeutic and microenvironmental stress. Consistent with this possibility, anti-PD-1 therapy has been reported to promote IFN-γ-dependent YAP1 nuclear translocation and phase separation in tumor cells, with hydrophobic interactions within the YAP1 coiled-coil domain contributing to puncta induction and C-terminal IDR-mediated weak interactions promoting puncta formation^63^. Elucidating the upstream mechanisms that induce YAP1 exon 6 skipping and nuclear puncta formation may reveal opportunities to limit the emergence of this adaptive, therapy-refractory state of cancer cells.

Clinical analyses provide exploratory support for the relevance of this mechanism in human tumors. In HGSOC specimens, higher *YAP1-E5.1/E7* transcript levels were associated with increased nuclear YAP puncta and greater YAP1–BRD4 proximity, and these measures were elevated in advanced-stage tumors. Clinical stage was associated with puncta number but not with integrated nuclear YAP1 signal per cell, suggesting that tumor progression may be more closely linked to punctate nuclear organization than to total nuclear YAP1 abundance. Exploratory TCGA analyses further suggested that a splicing pattern characterized by preferential use of *YAP1* exon 5.1 and exclusion of exon 6, consistent with YAP1-2α production, was associated with poorer overall survival. Together, these observations link the *YAP1* exon 6-skipping-associated punctate program to aggressive features of human HGSOC and provide a rationale for further isoform-resolved clinical validation.

Our findings further suggest that the YAP1-2α-associated TAZ/TEAD/BRD4 transcriptional program may represent a therapeutic vulnerability in selected tumor contexts. Selected YAP1-2α-associated HGSOC and gastric cancer models responded to K-975 and JQ1 at concentrations that did not cause marked nonspecific loss of cell viability. In addition, the inflammatory-response signature observed in YAP1-2α–dominant HGSOC tumors suggests that this program may also influence the tumor immune microenvironment. These findings provide a rationale for investigating whether YAP1 splicing toward YAP1-2α, nuclear YAP1 puncta, or downstream pathway activity could inform therapeutic strategies targeting TEAD- or BRD4-dependent transcription.

### Limitations of the study

This study has several limitations. First, the dynamic behavior of YAP1-2α puncta and the effects of IDR- and leucine-zipper-directed perturbations support a condensate-associated transcriptional state but do not definitively establish liquid–liquid phase separation. The complete molecular composition of these assemblies, the contributions of YAP1 post-translational modifications such as phosphorylation, and their genome-wide chromatin organization remain incompletely characterized. Quantitative biophysical studies, compositional analyses, and nascent-transcription profiling will be needed to refine this model. Second, our chromatin-related analyses focused on candidate regulatory regions rather than on comprehensive mapping and characterization of the full repertoire of super-enhancers potentially regulated by the proposed YAP1-2α–TAZ–TEAD axis. Public epigenomic datasets, chromatin interaction analyses, and functional transcriptional studies supported the identification and investigation of candidate super-enhancer-associated targets, but we did not perform H3K27ac, H3K4me1, BRD4, or H3K4me3 ChIP-seq or CUT&Tag profiling in the experimental models examined. Accordingly, this study does not establish the genome-wide distributions or changes in these chromatin features, nor does it comprehensively define the associated super-enhancer landscape. Systematic chromatin profiling will be required to test the proposed chromatin mechanisms more directly and to define their relationship to the biological phenotypes examined. Third, direct identification of endogenous YAP1-2α within individual nuclear puncta remains unresolved. To our knowledge, no validated imaging strategy currently enables selective visualization of endogenous YAP1-2α with sufficient specificity and sensitivity to distinguish it from other relevant YAP1 splice isoforms. Conventional tagging at an N- or C-terminus shared by multiple isoforms would not provide this selectivity. Moreover, detectable puncta under basal conditions cannot be assumed. Our working model proposes that a stress-associated splicing switch toward YAP1-2α increases its availability and favors puncta formation, but neither the requirement for this switch nor the presence of YAP1-2α within individual endogenous puncta has been directly established. Testing this model will require an isoform-selective detection strategy applicable to the relevant cellular state and validated to preserve endogenous splicing, expression, and puncta formation. The present findings therefore support a proposed model but do not constitute direct visualization of endogenous YAP1-2α condensates. Fourth, the upstream signals and splicing regulators responsible for *YAP1* exon 6 skipping remain unidentified. Our investigation focused on the biological consequences of isoform switching rather than the regulatory machinery that initiates it. Observations in selected resistant or disseminating populations do not establish whether stress directly induces the switch or selects cells with a pre-existing splicing state. Defining the causal links among stress-responsive signals, splicing regulators, and *YAP1* exon 6 selection will require a dedicated mechanistic investigation. Fifth, the molecular basis and biological significance of the residual puncta formed by TEAD-binding-deficient YAP1-2α(S94A) remain unresolved. Their persistence indicates that TEAD binding is not absolutely required for all YAP1-2α puncta formation. However, these structures should not be assumed to share the molecular composition, physical properties, or biological functions of puncta formed by wild-type YAP1-2α. Further work is needed to determine their formation mechanism and potential functions. Finally, the *BRCA*-proficient gastric cancer models provide complementary evidence in a tissue context distinct from HGSOC, but they do not directly establish applicability to *BRCA*-proficient HGSOC or isolate the contribution of *BRCA* status from other differences between the models. Validation in *BRCA*-proficient HGSOC and genetically matched models will be needed to address these questions. More broadly, the resistant-clone, xenograft, pharmacological, and exploratory clinical findings do not establish how frequently this program operates across tumor types, its prognostic value, or the extent to which tumors depend on it. Larger, clinically annotated cohorts, endogenous loss-of-function models, patient-derived and immunocompetent systems, and longitudinal treatment studies will be needed to define the translational relevance of the YAP1-2α-associated TAZ/TEAD/BRD4 transcriptional program.

## RESOURCE AVAILABILITY

### Lead contact

Further information and requests should be addressed to the lead contact, Masanori Hatakeyama.

### Materials availability

The materials generated in this study are available upon request to the lead contact.

### Data and code availability

All other data supporting the findings of this study are available within the article and its supplementary information files. This study did not generate the original code.

## ACKNOWLEDGMENTS

We thank Dr. Naoko Murata-Kamia for valuable discussions. This work was supported by a JSPS Grant-in-Aid for Scientific Research from the Japan Society for the Promotion of Science Grant numbers 21H04804 (M.H.) 24H00618 (M.H.), JP21K15526 (T.O.), and JP26K10118 (T.O.); P-PROMOTE from AMED Grant Number JP23ama221219h0001 (M.H.); and the Microbial Chemistry Research and Development Grant, funded by grant number 23A02 (T.O.) from the Microbial Chemistry Research Foundation.

## AUTHOR CONTRIBUTIONS

T.O. and M.H. conceptualization; T.O. and T.H. methodology; T.O., T.H., and Y.Y. formal analysis; T.O., T.H., and Y.Y. investigation; T.O. and M.H. writing-original draft; T.O., T.H., and M.H. writing-review & editing; M.H. supervision; M.H. project administration.

## Declaration of generative AI and AI-assisted technologies in the writing process

During the preparation of this work, the authors used Perplexity AI and Microsoft Copilot to assist with literature and journal-policy searches and to improve the manuscript’s language, readability, and organization. These tools were not used to generate, analyze, or interpret experimental data or to create figures. The authors reviewed, verified, and edited all content as needed after using these tools and take full responsibility for the published article.

## SUPPLEMENTAL INFORMATION

Document S1. Figures S1–S19.

## EXPERIMENTAL MODEL AND STUDY PARTICIPANT DETAILS

### Cell lines

UWB1.289 cells (ATCC, CRL-2945) were cultured in a 1:1 mixture of mammary epithelial cell growth medium (MEGM, Lonza, CC-3150) without GA-1000 (gentamicin-amphotericin B mixture) and RPMI-1640 (Wako, 189-02025), supplemented with 3% FBS (Gibco), 10 mM HEPES, and 1 mM sodium pyruvate. MDA-MB-436 (ATCC, HTB-130), MKN45-Luc (JCRB, JCRB1945), HSC-39 (JCRB, JCRB1864), and 39As8 (JCRB, JCRB1886) cells were cultured in RPMI-1640 supplemented with 10% FBS (Gibco), 10 mM HEPES, and 1 mM sodium pyruvate. Lenti-X 293T cells (Clontech Laboratories, 632180) were cultured in D-MEM (Wako, 044-29765) supplemented with 10% FBS (Gibco) and 1 mM sodium pyruvate. Cells were incubated at 37 °C in 5% CO_2_. To generate gene knockout (KO) cell lines, cells were transfected with the pX330-sgYAP1 or pX330-sgTAZ vector and a blasticidin S or puromycin resistance gene. After transfection, cells were selected with blasticidin S (10 μg/ml), or puromycin (1 μg/ml) and cloned by limiting dilution. Ten independent cisplatin-resistant clones were established from UWB1.289 cells after three months of culture under stepwise increases in cisplatin (MedChemExpress) concentration, ranging from 100 pM to 100 nM. All cell lines used in this study were obtained from the commercial sources listed in the Key Resources Table. Cell line authentication was not independently performed in our laboratory. Mycoplasma contamination was routinely monitored using the MycoAlert Mycoplasma Detection Kit (Lonza, #LT07-318), and all cell lines were confirmed to be negative for mycoplasma contamination before use in experiments.

### Human cancer patients data

PSI [percent-spliced-in index: the ratio of normalized read counts indicating inclusion of a transcript element over the total normalized reads for that event (both inclusion and exclusion reads)] data for *YAP1* mRNA splicing variants containing the γ-segment (referred to as *YAP1-E6*) and *YAP1* mRNA splicing variants containing the β-segment (referred to as *YAP1-E5.2*) were downloaded from the TCGA SpliceSeq (https://bioinformatics.mdanderson.org/TCGASpliceSeq/singlegene.jsp).^45^ RNA-seq data and corresponding clinical information for the TCGA-HGSOC cohort were downloaded from the cBioportal for Cancer Genomics (https://www.cbioportal.org/).^42–44^ The TCGA datasets were subgrouped and analyzed (see the METHOD DETAILS section for experimental details). A human ovarian cancer tissue array was purchased from US biomax (see the Key resource table).

### Mice

BALB/cAJcl-nu/nu female mice at five weeks of age were purchased from CLEA Japan and were used for xenograft assays. The mouse room was quarantined by airflow and kept at a constant temperature and humidity under specific pathogen-free (SPF) conditions. A humane endpoint was predefined as a loss of more than 20% of the initial body weight. All animal experiments were conducted in accordance with protocols approved by the Ethics Committee for Animal Experiments of the Institute of Microbial Chemistry, Microbial Chemistry Research Foundation (Animal Experiment Permit No. 2026-041) (see the METHOD DETAILS section for experimental details).

## METHOD DETAILS

### Three-dimensional (3D) overlay culture

UWB1.289 cells were suspended in growth medium supplemented with 2% Matrigel (CORNING, #356231), hereafter referred to as assay medium, and seeded onto Matrigel-coated 8-well chamber slides at 4 × 10^3^ cells/well. Eight days after seeding, spheroids were treated with 100 nM JQ1 (MedChemExpress, HY-13030), 1 μM K-975 (MedChemExpress, HY-138565), or DMSO as a mock control for 6 days. During cultivation, the assay medium was replaced every two days. After treatment, spheroids were fixed with 2% paraformaldehyde for 20 min and collected using a cell-recovery solution (CORNING, 354253). Fixed spheroids were permeabilized with 0.5% Triton X-100 in PBS for 20 min at room temperature and then subjected to immunostaining. Feret diameters of spheroids were measured from images captured 14 days after seeding using Fiji/ImageJ. For spheroid growth analysis, see the “Three-dimensional Cell Viability Assay” section.

### Expression vectors

All expression plasmids listed in the Key Resources Table were generated by subcloning cDNAs encoding the indicated proteins into the pCDH-CMV-MCS-EF1-Puro expression vector. Base substitution and deletion mutants were generated using a KOD-plus-Mutagenesis kit (TOYOBO, SMK-101) according to the manufacturer’s instructions. To generate shRNA expression vectors, the pLKO.5-U6-MCS-CMV-TurboGFP-hPGK-Puro vector was derived from pLKO.5-U6-CMV-TurboGFP-hPGK-Puro by introducing AgeI and EcoRI restriction sites using KOD-Plus Mutagenesis. ^64^ Annealed oligonucleotides encoding shRNAs targeting human YAP1 or BRD4 were inserted into this vector. For sgRNA expression vectors, annealed oligonucleotides encoding sgRNAs targeting human YAP1 or TAZ were inserted into the pX330 vector. The nucleotide sequences of all constructed vectors were confirmed by Sanger sequencing. See the Key resource table for RNAi and sgRNA target sequences.

### Preparation of lentivirus and infections

Lentivirus was generated by transfecting Lenti-X 293T cells with the pLKO.5 shRNA vector or the pCDH cDNA expression vector, along with the packaging plasmids psPAX2 and pMD2.G, using Lipofectamine 2000 (Invitrogen) according to the manufacturer’s protocol. At 72 hours after transfection, culture media containing recombinant lentiviruses were collected and concentrated using PEG-it (SBI). Lentivirus particle titers were determined using Lenti-X qRT-PCR Titration Kit (Clontech Laboratories) according to the manufacturer’s instructions. Lentivirus (MOI: 0.5 to 10) was added to cell culture medium with 10 μg/ml polybrene.

### Live-cell imaging

Cells transfected with mVenus-fused YAP1 isoforms were analyzed to confirm liquid-like properties (puncta fusion, disruption of puncta formation by Hex treatment, and FRAP). To assess the fusion behavior of YAP1-2α puncta, time-lapse fluorescence imaging was performed on cells expressing mVenus-fused proteins for 120 s using a FLUOVIEW FV1200 confocal microscope (Olympus). To observe disruption of YAP1-2α puncta formation by Hex treatment, transfected cells were treated with 1% Hex for 20 min. Twenty minutes after Hex treatment, the cells were washed and incubated for an additional 4 h. Fluorescence signals were detected at the indicated time points. To evaluate the fusion dynamics of YAP1-2α puncta, time-lapse fluorescence imaging was performed, and the aspect ratio (major axis/minor axis) of individual puncta was quantified using Fiji/ImageJ. Fusion events were identified as the coalescence of two condensates into a single structure, and temporal changes in aspect ratio were monitored throughout the fusion process. A decrease in the aspect ratio toward 1.0 after fusion was used as an indicator of liquid-like relaxation.

### Fluorescence recovery after photobleaching (FRAP)

The mVenus-fused YAP1-2α nuclear puncta and adjacent non-punctate regions were defined as ROIs and photobleached by laser irradiation. After photobleaching, time-lapse fluorescence images of the bleached ROIs were acquired for 120 s. Unbleached puncta served as controls. Images were acquired at 3.2 second intervals using a 40× objective lens. The fluorescence intensities of puncta from the time-lapse series were analyzed using FLUOVIEW FV1200 software. The fitting equation used to determine the diffusion coefficient “*D*” is shown below

*D* = (0.88・r^2^)/4t_1/2_

r: radius of the bleached area

t_1/2_: half-time of recovery

### Immunofluorescence staining

Cells were fixed with 2% or 4% paraformaldehyde in PBS for 15 min at room temperature or 37°C, permeabilized with 0.25% Triton X-100 in PBS for 10 min at room temperature, and blocked in blocking solution (5% BSA, 0.1% Tween 20, 0.05% NaN3 in PBS) for 60 min at room temperature. For spheroid staining, fixed and collected spheroids were permeabilized with 0.5% Triton X-100 in PBS for 20 min at room temperature. Blocked cells/spheroids were incubated with primary antibodies against YAP1 (CST, 12395, lot 3, 1:500; CST, 14074, lot 5, 1:500), HA-tag (CST, 2367, lot 5, 1:500 or CST, 3724, lot 13, 1:500), TAZ (CST, 83669, lot 2, 1:500), pan-TEAD (CST, 13295, lot 2, 1:500), Flag-tag (SIGMA, F3165, lot SLCP4941, 1:1000), BRD4 (CST, 13440, lot 9, 1:500), ki-67 (Thermo Fisher Scientific, 14-5698-80, lot 2047169, 1:500), Pol II (Santa Cruz Biotechnology, sc-47701, lot 1123, 1:500), SRRM2 (CST, 19700, lot 1, 1:500), PML (CST, 69789, lot 1, 1:500), NPM1 (CST, 92825, lot 1, 1:200), and Actin (CST, 3700, lot 17, 1:500). After primary antibody incubation, samples were incubated with Alexa Fluor-conjugated secondary antibodies (1:1000) and/or Alexa Fluor-conjugated phalloidin (Invitrogen). Nuclei were stained with DAPI (Wako, 043-18804, 1:1000) or DRAQ5 (CST, 4083, 1:1000).

The number of mVenus-fused YAP1 puncta was quantified using Fiji/ImageJ. The parameters used for puncta identification were as follows.

Image > Type > 16-bit; Adjust > Threshold > 50-65535

Analyze > Analyze particles > size (0.1-100), circularity (0.3-1.0)

The number of endogenous or HA-tagged YAP1 puncta was quantified using Fiji/ImageJ. The parameters used for puncta identification were as follows.

Image > Type > 16-bit

Process > Find Maxima > Prominence (10), Single Points

Analyze > Tools > ROI Manager > Add ROI > Measure

Regions of interest (ROIs) were defined based on DAPI or DRAQ5 staining. The parameters used for ROI identification were as follows.

Image > Type > 16-bit, Adjust > Threshold > 30-65535

Analyze > Analyze particles > size (20-infinity), circularity (0.2-1.0)

Cells expressing mVenus or mVenus-fused proteins were identified based on mVenus fluorescence signals. The parameters used for ROI identification were as follows.

Image > Type > 16-bit, Adjust > Threshold > 10-65535

Analyze > Analyze particles > size (50-infinity), circularity (0.0-1.0)

The extent of colocalization between the indicated fluorescent signals was quantified using the Coloc2 plugin in Fiji/ImageJ. Manders’ colocalization coefficients were calculated using Costes-derived automatic thresholds. The significance of colocalization within individual images was assessed with Costes’ randomization test (100 iterations), which compares the observed correlation with that of randomized images. For population-level analysis, Manders’ colocalization coefficients from individual cells were compared between groups, and statistical significance was assessed using a paired two-tailed *t* test.

### Proximity ligation assay (PLA)

Cells were washed with PBS containing 1 mM MgCl₂ and fixed with 4% paraformaldehyde in PBS for 15 min at room temperature, followed by permeabilization with 0.25% Triton X-100 in PBS for 10 min at room temperature. Cells were then incubated overnight at 4°C with primary antibodies against YAP1 (CST, #12395, lot 3, 1:500), HA tag (CST, #2367, lot 5, 1:500; or CST, #3724, lot 13, 1:500), TAZ (CST, #83669, lot 2, 1:500), pan-TEAD (CST, #13295, lot 2, 1:500), FLAG tag (Sigma-Aldrich, #F3165, lot SLCP4941, 1:1000), or BRD4 (CST, #13440, lot 9, 1:500), and were subsequently subjected to proximity ligation assay (PLA). PLA was performed using a Duolink *In situ* PLA kit (SIGMA; Duo92007, Duo92002, Duo92004) according to the manufacturer’s instructions. PLA signals were quantified using Fiji/ImageJ and analyzed with GraphPad Prism software. The criteria and parameters used to identify PLA signals were as follows.

Image > Type > 16-bit

Process > Find Maxima > Prominence (20), Single Points

Analyze > Tools > ROI Manager > Add ROI > Measure

See also our previous STAR Protocols.^65^

### Förster resonance energy transfer (FRET) assay

FRET analysis was performed using a FLUOVIEW FV1200 confocal microscope (Olympus). For sensitized emission FRET, SECFP was excited with a 440 nm laser, and fluorescence was collected at 460–500 nm for SECFP and 520–550 nm for mVenus. Sequential image acquisition was used to minimize spectral cross-talk. Cells expressing SECFP-fused YAP1-2α alone or mVenus-fused TAZ alone served as controls for donor bleed-through and acceptor cross-excitation correction. The specificity of the FRET signal was further validated by photobleaching the donor fluorophore (SECFP), which eliminated the FRET signal and confirmed that the observed energy transfer reflected a specific interaction between the fluorophore-tagged proteins. In the combined FRET/PLA analysis, PLA images used to visualize TAZ homodimers were acquired before FRET imaging. Quantification of YAP1-2α/TAZ heterodimer puncta based on FRET signals, quantification of PLA signals corresponding to TAZ homodimers, and Manders’ colocalization analysis was performed as described in the “Immunofluorescence Staining” and “Proximity Ligation Assay (PLA)” sections of the STAR Methods. The proportions of YAP1-2α/TAZ heterodimer puncta and TAZ/TAZ homodimer puncta among total TAZ puncta were quantified by determining the presence of FRET signals (representing YAP1-2α/TAZ heterodimers) and PLA signals (representing TAZ/TAZ homodimers) within ROIs generated from the fluorescence signals of TAZ puncta. TAZ puncta were identified using the same detection criteria as those applied for mVenus-YAP1-2α puncta. FRET and PLA signals were detected using the following parameters: Image > Type > 16-bit and Process > Find Maxima > Prominence (10 for FRET and 20 for PLA), followed by Analyze > Measure within ROIs defined by total TAZ puncta.

### Immunohistochemistry

The human ovarian cancer tissue arrays (OV1004a, OV0071a, OV242d, OV244, and T112-L47; US Biomax) were deparaffinized and rehydrated. Antigen retrieval was performed by boiling for 15 minutes in 1× Target Retrieval (ACD). To block endogenous peroxidase activity, the tissue array was incubated with 3% hydrogen peroxide in double-distilled water for 10 min at room temperature. The array was then incubated overnight at 4 °C with a primary antibody against YAP1 (CST, 14074, lot 5, 1:400; CST, 12395, lot 3, 1:200), BRD4 (CST, 13440, lot 9, 1:200), and TAZ (CST, 83669, lot 2, 1:200). After the antigen-antibody reaction, the array was incubated for 60 min at room temperature with Alexa Fluor-conjugated secondary antibodies (1:1000). After the secondary antibody reaction, the nuclei were stained with DAPI solution (1:1000) for 20 min at room temperature. Fluorescence images were obtained using the FLUOVIEW FV1200 confocal microscope. H&E staining was performed using the Hematoxylin and Eosin Stain kit (ScyTek Laboratories). For the proximity ligation assay (PLA), antigen-retrieved tissue sections were treated to block endogenous peroxidase activity and incubated overnight at 4 °C with primary antibodies against YAP1 (CST, 14074, lot 5, 1:200) and BRD4 (Santa Cruz, sc-518021, lot E2226, 1:200). PLA was then performed using the Brightfield Duolink *In situ* PLA kit (SIGMA, Duo92012, Duo92002, Duo92004) according to the manufacturer’s instructions. PLA signals were developed for 2 min using the substrate solution, followed by hematoxylin counterstaining. Nuclear PLA signals were quantified using QuPath^66^ under the following analysis conditions.

i. Ref ROI > annotation > Analyze > Estimate stain vectors > Auto
ii. Measure ROI > new annotation > Analyze > Cell detection > positive cell detection > Hematoxylin OD, Background radius (100 px), Sigma (8)
iii. Analyze > Calculate features > add intensity features > Region (ROI), DAB, mean, median
iv. Measure > show annotation measurements

### In situ hybridization (BaseScope)

Deparaffinized and rehydrated tissue arrays were analyzed by BaseScope in situ hybridization using the BaseScope Detection Reagent Kit v2-RED (ACD), following the manufacturer’s instructions. ACD designed the human *YAP1-1α/2α*-isoform (*YAP1-E5.1/E7*)-specific probe (BA-Hs-YAP1-tv3-E5E6). *YAP1-E5.1/E7* mRNA expression levels were quantified by measuring BaseScope signals with Fiji/ImageJ. Expression was scored as follows: 0, no staining or fewer than 1 dot per 20 cells; 1+, 1 dot/cell; 2+, 2-3 dots/cell, with no or very few dot clusters; 3+, 4-10 dots/cell, with dot clusters in fewer than 10% of positive cells; 4+, >10 dots/cell, with dot clusters in more than 10% of positive cells. Staining scores were assigned according to the BaseScope scoring guidelines.

### Immunoprecipitation and immunoblotting

The cells were washed in ice-cold PBS and lysed in lysis buffer (100 mM NaCl, 50 mM Tris-HCl pH 7.5, 5 mM EDTA, 2 mM Na_3_VO_4_, 10 mM NaF, 10 mM β-glycerophosphate, 2 mM PMSF, 10 µg/ml leupeptin, 10 µg/ml trypsin inhibitor, 10 µg/ml aprotinin, and 0.4% NP-40). Cell lysates were incubated with anti-TAZ (CST, 83669, lot 2, 1:500), anti-HA (CST, 3724, lot 13, 1:500), anti-Flag (SIGMA, F3165, lot SLCP4941, 1:1000), anti-YAP1 (CST, 14074, lot 5, 1:500), or isotype control IgG, then incubated with protein G-beads. The beads were washed five times with lysis buffer, and the immunoprecipitates were eluted with SDS sample buffer. The immunoprecipitates and total cell lysates were subjected to SDS-PAGE and immunoblotted. Images were captured using the LAS-4000 imager (FUJIFILM). To compare protein expression levels, protein band intensities from three biological replicates were collected and quantitatively analyzed using the LAS-4000 luminoimage analyzer (FUJIFILM). Exposure time for detection was automatically calculated to stay within the LAS-4000 dynamic range, and signal linearity was maintained within this range.

### Chromatin immunoprecipitation

To evaluate interactions between genomic DNA and target proteins, DNA samples were prepared by CUT&RUN and ChIP-re-ChIP. CUT&RUN was performed using the CUT&RUN Assay Kit (CST) according to the manufacturer’s instructions, employing anti-BRD4 (CST, 13440, lot 9, 1:50), anti-H3K9me3 (CST, 13969, lot 6, 1:50), anti-H3K27ac (CST, 8173, lot 8, 1:50), anti-H3K4me1 (CST, 5326, lot 7, 1:50), anti-HA (CST, 3724, lot 13, 1:50), anti-TAZ (CST, 83669, lot 2, 1:50), or anti-pan-TEAD (CST, 13295, lot 2, 1:50) antibody. Rabbit IgG (CST, 66362, lot 3, 1:20) served as the isotype control. For ChIP-re-ChIP, *YAP1*-KO UWB1.289 cells were transfected with 6xHis-tagged YAP1-2α and treated with K-975 or DMSO. Twenty-four hours after K-975 treatment, formaldehyde cross-linking (CST, 12606) was performed, followed by chromatin isolation and sonication. Cross-linked chromatin was pulled down using Ni Sepharose Excel (Cytiva) and eluted with 500 mM imidazole. The imidazole was replaced with a lysis buffer by centrifugal ultrafiltration (Millipore, UFC903024), and immunoprecipitation was performed using anti-TAZ antibody (CST, 83669, lot 2) or rabbit IgG (CST, 66362, lot 3) as an isotype control. Chromatin bound to the YAP1-2α/TAZ complex was reversed and the DNA fragments were purified using spin columns (CST, 14209). Purified DNA samples were subjected to PCR or qPCR analysis.

### Detection of chromatin interactions (ChIP-loop)

After formaldehyde crosslinking, chromatin was isolated and digested with restriction enzymes [NEB: StuI (R0187), SmaI (R0141); Takara: BciT130I (1232), AfaI (1116)]. Crosslinked chromatin was immunoprecipitated using either an anti-HA antibody (CST, 3724, lot 13, 1:100) or an anti-GFP antibody (MBL, 598, lot 080, 1:50), together with ChIP-grade Protein G agarose beads (CST, 9007, lot 40). The resulting samples were divided into two aliquots. One aliquot was ligated with Ligation High Ver.2 (TOYOBO), whereas the other was processed as a ligation-negative control without ligase. Crosslinks were reversed, and DNA was purified using a spin column (CST, 14209) according to the manufacturer’s instructions. Specific chromatin interactions were detected by PCR. Amplified products were separated by agarose gel electrophoresis, visualized by ethidium bromide staining, and quantified using Fiji/ImageJ.

### Quantitative PCR (qPCR)

Total RNA was extracted from cells using TRIzol reagent (Invitrogen) and reverse-transcribed using the High-Capacity cDNA Reverse Transcription Kit (Thermo Fisher Scientific, 4368814). Relative mRNA expression was quantified using the QuantStudio 3 Real-Time PCR System (Thermo Fisher Scientific) with SYBR *Premix Ex Taq* Tli RNaseH Plus (TaKaRa). *18S* rRNA served as the internal control. Data were analyzed using the comparative Ct (ΔΔCt) method. For ChIP-qPCR analysis, purified DNA fragments from CUT&RUN and ChIP-re-ChIP were quantified using the QuantStudio 3 Real-Time PCR System with KOD-SYBR qPCR Mix (TOYOBO). See the Key resource table for primer sequences.

### Detection of YAP1 splicing isoforms

Total RNA was extracted from the cells using TRIzol reagent (Invitrogen, 15596026) and reverse-transcribed using oligo(dT) primers and SuperScript II Reverse Transcriptase (Invitrogen, 18064014). The resulting cDNA was amplified by PCR to detect the YAP1 exon 6 sequence, which encodes the γ-segment. After PCR, the amplicon was loaded onto a 2% TAE-agarose gel and electrophoresed. Amplicon sequence analysis for YAP1 isoform identification was performed by excising appropriately separated bands following gel electrophoresis, purifying the DNA fragments, performing nested PCR using the purified products as templates, and subjecting the resulting amplicons to Sanger sequencing.

### Luciferase reporter assay

Luciferase reporter assays were performed using the Dual-Luciferase Reporter Assay System (Promega). The pRL-CMV Control plasmid was used to normalize transfection efficiency. The pδ51-LUCII-8×GT-IIC luciferase reporter plasmid was used as a TEAD-responsive reporter, whereas the pδ51-LUCII-empty luciferase reporter plasmid was used as a negative control. To evaluate the effects of YAP1-2α mutants on TEAD transcriptional activity, *YAP1*-knockout UWB1.289 cells were co-transfected with the TEAD reporter plasmid, pRL-CMV Control plasmid, and the indicated mVenus-tagged YAP1-2α expression vector using Lipofectamine 2000 (Invitrogen). Forty-eight hours after transfection, cells were collected in passive lysis buffer (Promega). Luminescence was measured using a Centro XS^3^ LB960 (BERTHOLD). Relative luciferase activity was calculated as a fold change relative to control samples. To confirm protein expression levels of the mVenus-fused YAP1-2α mutants, the remaining cell lysates were subjected to immunoblot analysis.

### Cell viability assay

Sensitivity to anticancer agents and 2D cell proliferation were measured by MTS assay using CellTiter 96 AQueous One Solution Cell Proliferation Assay (Promega, G3580) according to the manufacturer’s instructions. To examine drug sensitivity, 1,000 cells were seeded into each well of 96-well plates and treated with stepwise concentrations of anticancer agents (cisplatin, sorafenib, trametinib) for 72 h. To examine sensitivity to olaparib, 500 cells were seeded into each well of 96-well plates and treated with a stepwise concentration of olaparib for 6 days. After treatment with anticancer agents, cells were incubated with the MTS assay reagent at 37 °C in 5% CO_2_. After 60-90 min of incubation, the absorbance at 490 nm was measured using Multiskan FC (Thermo Fisher Scientific). Relative cell viability was calculated by normalizing to mock-treated cells. In experiments designed to examine the effects of JQ1 or K-975, cells were treated with 100 nM JQ1, 1 μM K-975, or DMSO as a mock control for 3 days, then seeded into a 96-well plate.

### Three-dimensional cell viability assay

To assess anchorage-independent growth, cells were seeded at 500 cells per well in an Ultra-Low Attachment 96-well plate (CORNING) and cultured for 12 days. The medium was replaced every 2 days. Spheroids were lysed using CellTiter-Glo 3D Cell Viability Assay Reagent (Promega, G9683). Immediately after lysis, 200 μl of cell lysate from each well was transferred to a white 96-well plate (Greiner), and luminescence was measured using a Centro XS^3^ LB960. Dilutions were performed as needed to ensure that the luminescence signal remained within the linear range.

### Ferroptosis assay

Cells were treated with 100 nM JQ1, 1 μM K-975, or DMSO as a mock control for 3 days, then seeded into a 96-well plate (Greiner) and further treated with chlorido[N,N’-Disalicylidene-1,2-Phenylenediamine]Iron(III) (Cayman Chemical Company, 28788) or DMSO as a mock control for 48 h. Cell viability was assessed using the CellTiter-Glo 3D Cell Viability Assay (Promega) according to the manufacturer’s instructions. Luminescence was measured using a Centro XS^3^ LB960 luminometer. To confirm that the observed cell death was attributable to ferroptosis, cells were co-treated with ferrostatin-1 (Selleck, S7243), a ferroptosis inhibitor.

### Migration and invasion assay

After treatment with 100 nM JQ1, 1 μM K-975, or DMSO as a mock control for 3 days, cells were seeded into Transwell inserts (Costar, 3464) placed in 24-well plates (Costar, 3526). Transwell inserts coated with type-I-A collagen gel (Wako), prepared according to the manufacturer’s protocols, were used to examine the ability of the cells to infiltrate the extracellular matrix (invasion assay). Cells were then cultured in medium containing 0.1% FBS for 48 h (chemotaxis assay) or 2 weeks (invasion assay). The bottom wells were filled with medium containing 10% FBS. After incubation for the appropriate period, cells that had migrated or invaded into the lower wells were lysed with CellTiter-Glo 3D Cell Viability Assay Reagent. Cell number was estimated by measuring luminescence using Centro XS^3^ LB960.

### Analysis of HGSOC RNA-seq data

For subgrouping the TCGA cohorts, *YAP1* isoform expression was classified as “*YAP1-1/2*-*α*–dominant” [PSI (*YAP1-exon 6*) < median, PSI (*YAP1-exon 5.2*) < median], “*YAP1-1/2-β*–dominant” [PSI (*YAP1-exon 6*) < median, PSI (*YAP1-exon 5.2*) > median], “*YAP1-1/2-γ*–dominant” [PSI (*YAP1-exon 6*) > median, PSI (*YAP1-exon 5.2*) < median], or “*YAP1-1/2-δ*–dominant” [PSI (*YAP1-exon 6*) > median, PSI (*YAP1-exon 5.2*) > median]. HGSOC patients (n = 425) were divided into the “*YAP1-1/2*-*α*–dominant” (n = 131), “*YAP1-1/2-β*–dominant” (n = 83), “*YAP1-1/2-γ*–dominant” (n = 78), and “*YAP1-1/2-δ*–dominant” (n = 133). TPM counts for “*YAP1-1/2*-*α*–dominant” and “*YAP1-1/2-γ*–dominant” were analyzed using iDEP v2.4.0 web tool (https://bioinformatics.sdstate.edu/idep/)^67^ to perform DEG analysis. Genes with increased expression in “*YAP1-1/2*-*α*–dominant” vs. “*YAP1-1/2-γ*–dominant” (FDR < 0.1, FC >1.5) were further subjected to gene enrichment analysis (Hallmark MSigDB) using the ShinyGO gene-set enrichment tool (https://bioinformatics.sdstate.edu/go/).^68^ Multivariable Cox proportional hazards regression analysis was used to assess the association of *YAP1* mRNA expression and alternative splicing with overall survival (OS). Covariates included age at diagnosis, FIGO stage, *YAP1* mRNA expression [transcripts per million (TPM)], and the percent spliced-in (PSI) values for *YAP1* exon 5.2 and exon 6. Hazard ratios (HRs) and 95% confidence intervals (CIs) were estimated for each variable. Kaplan-Meier survival analysis was performed to evaluate overall survival in the TCGA-HGSOC cohort. Patients were stratified into *YAP1* exon 5.2/exon 6-skipped (exon 5.2 PSI < 0.35 and exon 6 PSI < 0.65, n = 75) and exon 5.2/exon 6-included (exon 5.2 PSI > 0.35 and exon 6 PSI > 0.65, n = 74) splicing subgroups based on PSI values. Differences in survival distributions between the two groups were assessed using the Gehan-Breslow-Wilcoxon test.

### Prediction of candidate super-enhancer-associated regulatory regions

ROSE-identified super-enhancer elements in serous ovarian cancer cells (GSM3328052: PEO1)^38^ and triple-negative breast cancer cells (GSM6132824: MDA-MB-436)^39^ were obtained from SEdb 3.0 (http://www.licpathway.net/sedb). TEAD1 and TEAD4 binding motifs were obtained from JASPAR (https://jaspar.elixir.no/).^69^ The downloaded data were visualized using IGV (https://igv.org/app/).^70^

### Tumor xenograft assay

Firefly luciferase-expressing MKN45 cells (MKN45-Fluc) infected with lentiviruses expressing HA-tagged YAP1-2α, HA-tagged YAP1-2γ, or an empty control vector were selected with puromycin. Puromycin-resistant cells were suspended in PBS at 2.5 x 10^7^ cells/ml and intraperitoneally injected into 5-week-old female nude mice (5 x 10^6^ cells in 0.2 ml per mouse). Mice were maintained under specific pathogen-free conditions for 14 days. A humane endpoint was predefined as a loss of more than 20% of the initial body weight. Bioluminescence imaging was performed using an in vivo imaging system (IVIS) on days 3, 7, and 14 after tumor cell implantation. For imaging, mice were anesthetized with isoflurane (dial setting, 2.5; O_2_ flow rate, 2 L/min). D-luciferin (Revvity) was administered intraperitoneally at a concentration of 30 mg/ml in a volume of 150 μl per mouse. Bioluminescence images were acquired 5 min after D-luciferin administration using an exposure time of 10 s. Relative total intraperitoneal bioluminescence signals on days 7 and 14 were normalized to the day 3 post-implantation signal, which was set to 1. Mice were euthanized 14 days after implantation, and the intestines and peritoneal tissues, including the omentum, were collected. Nodular tumors arising on the peritoneum and omentum were excised and weighed. For total RNA extraction, nodular tumor tissues were homogenized in TRIzol reagent, and total RNA was purified according to the manufacturer’s instructions. All animal experiments were conducted in accordance with protocols approved by the Ethics Committee for Animal Experiments of the Institute of Microbial Chemistry, Microbial Chemistry Research Foundation (Animal Experiment Permit No. 2026-041).

## QUANTIFICATION AND STATISTICAL ANALYSIS

In this study, unpaired *t* tests (Student’s *t* test and Welch’s *t* test), paired *t* tests, and Mann-Whitney U tests were used for comparisons between two groups. The Gehan-Breslow-Wilcoxon test was used for survival analyses. Statistical analyses were performed using Microsoft Excel and GraphPad Prism software. Details of the statistical tests used, exact *n* values, the definition of *n*, measures of central tendency, and measures of variability are provided in the corresponding figure legends. Detailed descriptions of each analysis are provided in the relevant Methods sections. All experiments presented in the manuscript were repeated as technical or biological replicates.

