## Supplemental Figures 1-19 for "YAP1-2α–TAZ heterodimerization promotes BRD4-associated LLPS-like assembly at candidate super-enhancers and cancer cell malignancy"

### Supplemental Figure 1

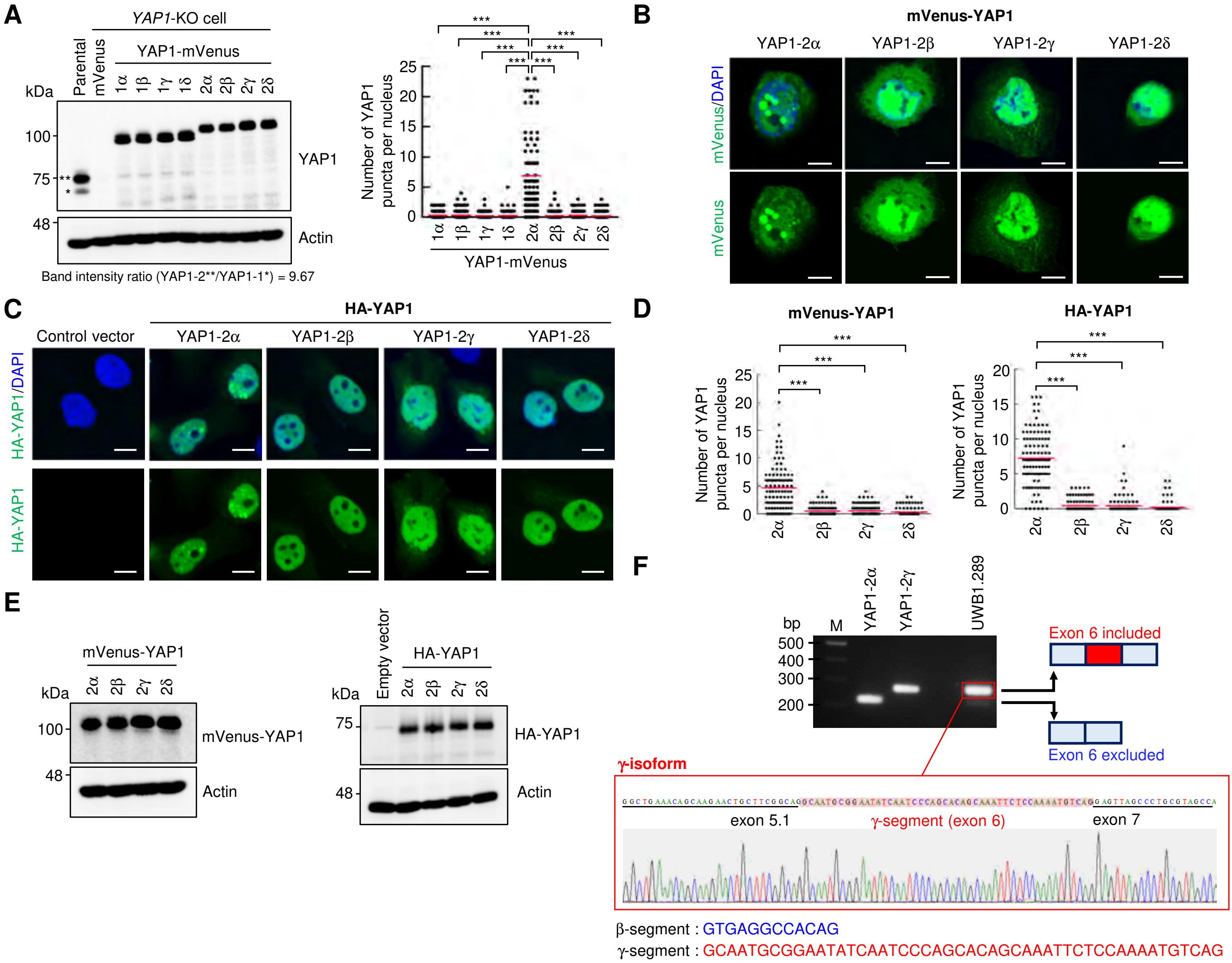


**Figure S1. Isoform-dependent nuclear YAP1 condensate formation (Related to Figure 1)**

(**A**) Related to Figure 1B. Immunoblot analysis of endogenous YAP1-1 (*) and YAP1-2 (**) in parental UWB1.289 cells, and C-terminally mVenus-tagged YAP1 (YAP1-mVenus) isoforms in *YAP1*-KO UWB1.289 cells. In parental UWB1.289 cells, the band density ratio of YAP1-2 (**) to YAP1-1 (*) is 9.87 (left). Number of YAP1-mVenus puncta per nucleus. ***$p < 0.001$ (Mann-Whitney U test). $n = 100$ nuclei per group (right).

(**B** and **C**) Fluorescence imaging analysis of *YAP1*-KO UWB1.289 cells expressing the indicated YAP1 isoforms with an N-terminal mVenus tag (B; mVenus-YAP1) or an N-terminal HA tag (C; HA-YAP1). mVenus fluorescence is shown in green.

(**D**) Number of nuclear YAP1 puncta in Figure S1B (left panel) or Figure S1C (right panel). $n = 100$ nuclei per group. ***$p < 0.001$ (Mann-Whitney U test). Red lines indicate the mean.

(**E**) Immunoblot analysis with the indicated antibodies (related to Figure S1B; left, and Figure S1C; right).

(**F**) RT-PCR analysis of *YAP1* mRNA isoforms with or without exon 6 in UWB1.289 cells (upper). The upper and lower bands correspond to *YAP1* transcripts containing and lacking exon 6, respectively. The upper band was purified for Sanger sequencing (bottom). Sequencing showed that the exon 6-containing transcript, which predominates in parental UWB1.289 cells, encodes the YAP1-1/2γ isoforms that contain the γ-segment but lack the β-segment (lower). M indicates the DNA size marker.

(**A** and **D**) YAP1 puncta were quantified using the predefined criteria for mVenus-fused YAP1, as described in the STAR Methods (Immunofluorescence Staining section).

All scale bars represent 10 µm.

Supplemental Figure 2

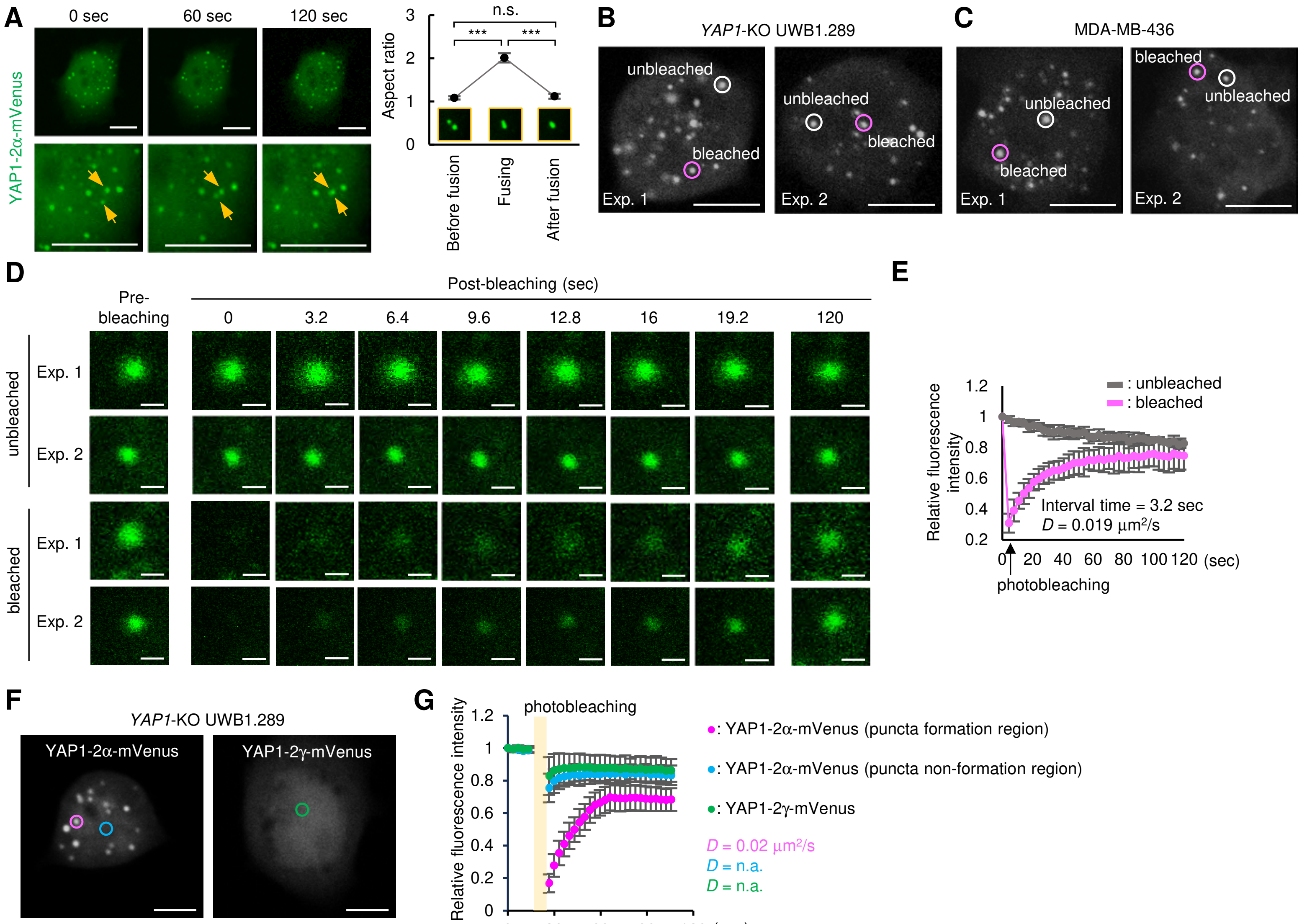


**Figure S2. Live-cell imaging of YAP1-2α nuclear condensates (Related to Figure 1)**

(**A**) Fusion of YAP1-2α puncta. Yellow arrows indicate puncta undergoing fusion. Fusion dynamics were assessed by measuring the aspect ratio of YAP1-2α puncta before, during, and after fusion. $^{***}p < 0.001$; n.s., not significant (Welch's *t* test). $n = 5$ fusion events.

(**B**) Full nuclear images of the cells used for FRAP analysis (related to Figure 1H). The cells are labeled Exp. 1 and Exp. 2. White circles indicate non-photobleached puncta, while pink circles indicate photobleached puncta.

(**C–E**) FRAP analysis of MDA-MB-436 cells expressing YAP1-2α-mVenus. Full nuclear images of the cells used for FRAP analysis are shown in (C), along with representative time-lapse images in (D). Scale bars, 1 μm. Fluorescence intensities of puncta in a series of time-lapse images were analyzed (photobleached, $n = 5$; unbleached, $n = 5$; technical replicates). *D* indicates the diffusion coefficient. Error bars show the mean ± SD (E).

(**F** and **G**) FRAP analysis of *YAP1*-knockout UWB1.289 cells expressing YAP1-2α-mVenus or YAP1-2γ-mVenus. Representative whole-nucleus images of cells used for FRAP analysis (F). Fluorescence recovery curves generated from post-photobleaching time-lapse images. FRAP was performed on YAP1-2α puncta and non-punctate nuclear regions in cells expressing YAP1-2α-mVenus or YAP1-2γ-mVenus ($n = 10$ cells per group). Fluorescence in non-punctate nuclear regions of cells expressing YAP1-2α-mVenus or YAP1-2γ-mVenus, which do not form nuclear YAP1 puncta, recovered too rapidly after photobleaching to allow reliable estimation of diffusion coefficients. By contrast, YAP1-2α-mVenus puncta showed substantially slower fluorescence recovery, allowing quantitative FRAP analysis. These results indicate that YAP1-2α puncta have dynamics distinct from diffusely distributed nuclear YAP1-2α and YAP1-2γ. *D*, diffusion coefficient; n.a., not applicable. Error bars indicate mean ± SD (G).

(A–C, and F) Scale bars represent 10 μm.

Supplemental Figure 3

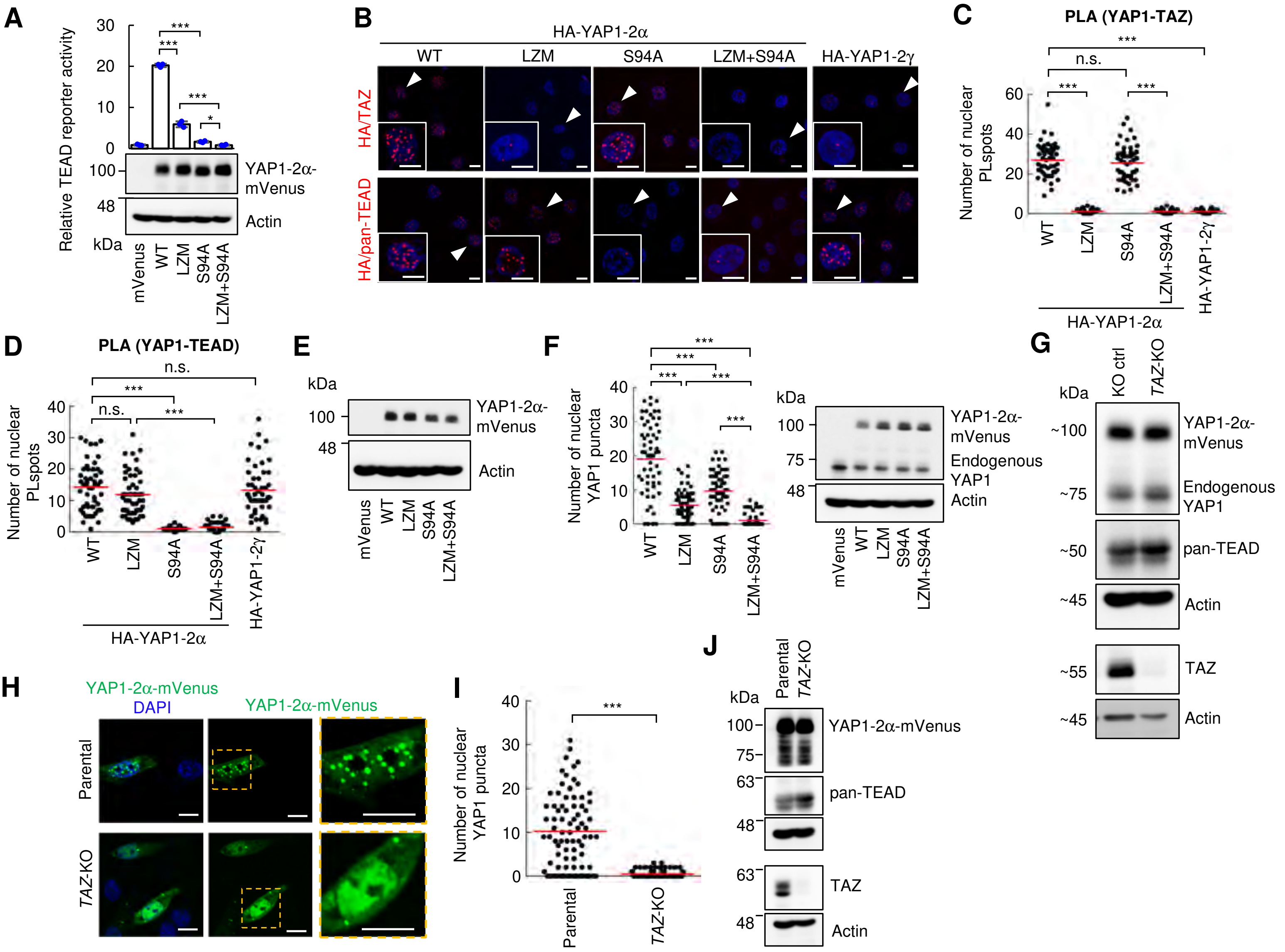


**Figure S3. YAP1-2α–TAZ heterodimerization is required for nuclear condensate formation (Related to Figure 2)** (**A**) TEAD reporter assay in *YAP1*-KO UWB1.289 cells expressing the indicated YAP1-2α variants fused to mVenus at the C terminus. Error bars indicate the mean ± SD ($n$ = 3 technical replicates; top). Expression of the mVenus-fused YAP1-2α variants (middle). Actin served as the protein-loading control (bottom). (**B**) Interaction of YAP1-2α with endogenous TAZ (top) or TEAD (bottom) in UWB1.289 cells, visualized by PLA as red PL spots. Nuclei indicated by white arrows are magnified. (**C** and **D**) The number of nuclear PL spots is shown in the graph (C: interaction between HA-YAP1-2α and TAZ; D: interaction between HA-YAP1-2α and TEAD). $n$ = 50 nuclei in each group. (**E**) Related to Figure 2B. Expression of the indicated YAP1-2α variants in *YAP1*-KO UWB1.289 cells, confirmed by immunoblotting. (**F**) Nuclear puncta formation in MDA-MB-436 cells expressing mVenus-fused YAP1-2α mutants. The number of YAP1 puncta in the nucleus is shown on the left. $n$ = 60 nuclei per group across three independent biological replicates. Protein expression was confirmed by immunoblotting (right). (**G**) Related to Figure 2C. Immunoblot analysis of *TAZ*-KO UWB1.289 cells expressing YAP1-2α-mVenus, using the indicated antibodies. (**H–J**) *TAZ*-KO MDA-MB-436 cells were transfected with a YAP1-2α-mVenus vector. Fluorescence images (H). Number of YAP1-2α-mVenus puncta in the nucleus. $n$ = 80 cells per group from three independent biological replicates (I). Confirmation of *TAZ* knockout, YAP1-2α-mVenus expression, and TEAD expression by immunoblotting (J). (A, C, D, F, and I) ***$p$ < 0.001; *$p$ < 0.05; n.s., not significant (A: Student's $t$ test; C, D, F, and I: Mann-Whitney U test). Red lines indicate the mean. (F and I) YAP1 puncta were quantified using the predefined criteria for mVenus-fused YAP1, as described in the STAR Methods. All scale bars represent 10 μm. (G and J) The same cell lysates were resolved on separate (G) or the same (J) SDS-PAGE gel(s). After transfer, the membrane was cut, and the indicated proteins were detected separately.

Supplemental Figure 4

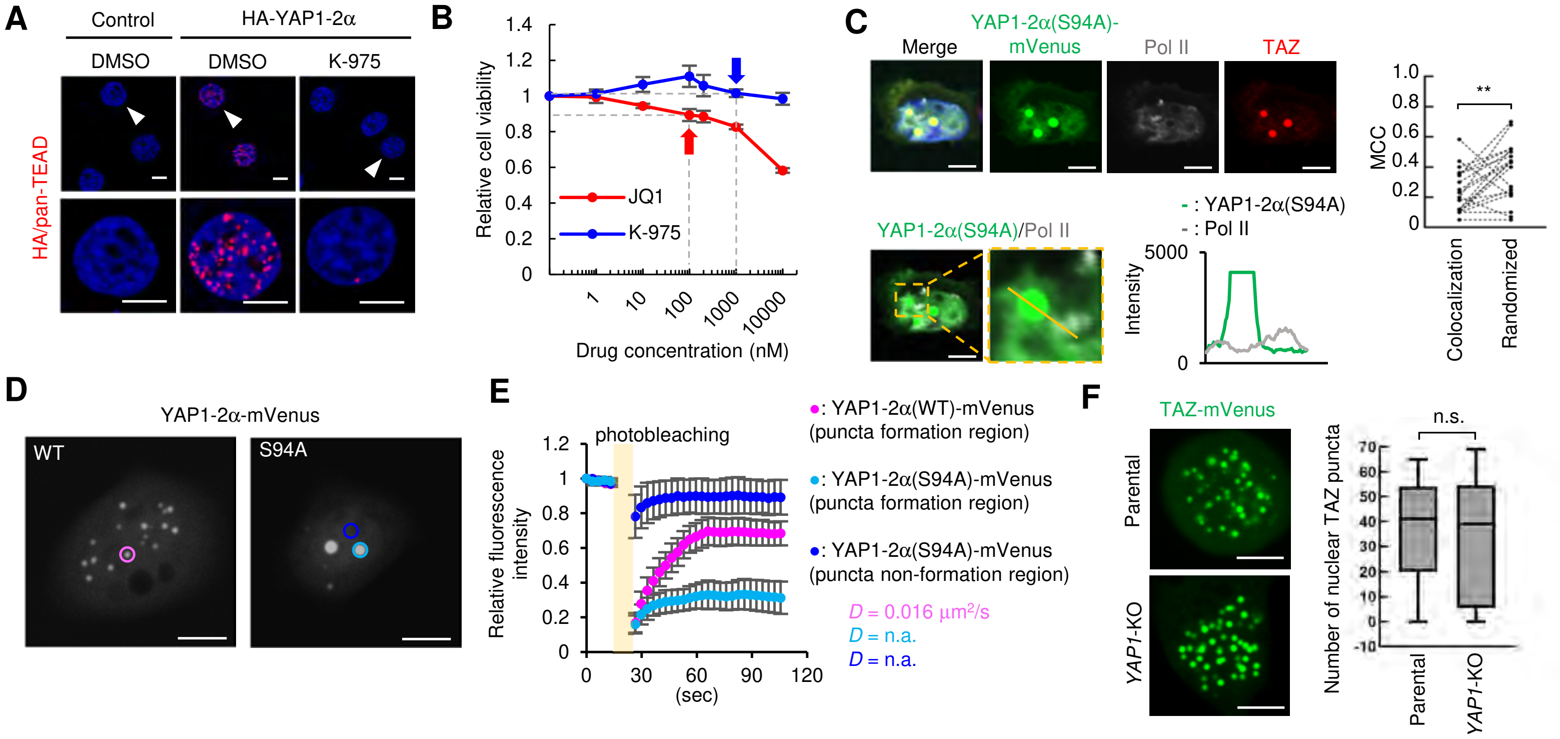


**Figure S4. YAP1-2α/TAZ/TEAD association is required for Pol II-positive nuclear condensate formation (Related to Figure 2)**

(**A**) PLA was used to assess whether K-975 inhibits the physical interaction between HA-YAP1-2α and endogenous TEAD in UWB1.289 cells.

(**B**) Drug sensitivity of UWB1.289 cells to K-975 and JQ1. Cell viability was assessed by an MTS assay 72 h after treatment. Arrows indicate the drug concentrations used in this study, at which no detectable cytotoxicity was observed.

(**C**) Immunofluorescence images of YAP1-2α(S94A)-mVenus (green), TAZ (red), and Pol II (gray). Overlap of fluorescence intensities between YAP1-2α(S94A) puncta and Pol II puncta in the region indicated by the yellow line (left). MCC was quantified in 20 cells. The observed colocalization of Pol II puncta with YAP1-2α(S94A) puncta (M1) was significantly lower than expected under random distribution, as determined by Costes randomization analysis (randomized M1) (right), indicating that Pol II was not significantly colocalized with YAP1-2α(S94A) puncta. $^{**}p < 0.01$ (paired *t* test).

(**D** and **E**) FRAP analysis of *YAP1*-KO UWB1.289 cells expressing YAP1-2α(WT)-mVenus or YAP1-2α(S94A)-mVenus. Representative whole-nucleus images of cells used for FRAP analysis are shown in (D). Fluorescence recovery curves were generated from time-lapse images acquired after photobleaching. FRAP was analyzed in YAP1-2α(WT) puncta, YAP1-2α(S94A) puncta, and non-punctate regions of YAP1-2α(S94A)-expressing cells ($n$ = 10 cells per group). *D*, diffusion coefficient. n.a., not applicable. Error bars indicate the mean ± SD (E).

(**F**) Effect of *YAP1* knockout on TAZ homodimer-mediated LLPS in UWB1.289 cells. TAZ/TAZ LLPS (left). Number of nuclear TAZ puncta in parental or *YAP1*-KO cells (right). Boxes include the 25th–75th percentiles with bars indicating medians. n = 13 cells for each group. n.s., not significant (Mann-Whitney U test). TAZ puncta were quantified using the predefined criteria for mVenus-fused YAP1, as described in the STAR Methods.

All scale bars represent 10 μm.

Supplemental Figure 5

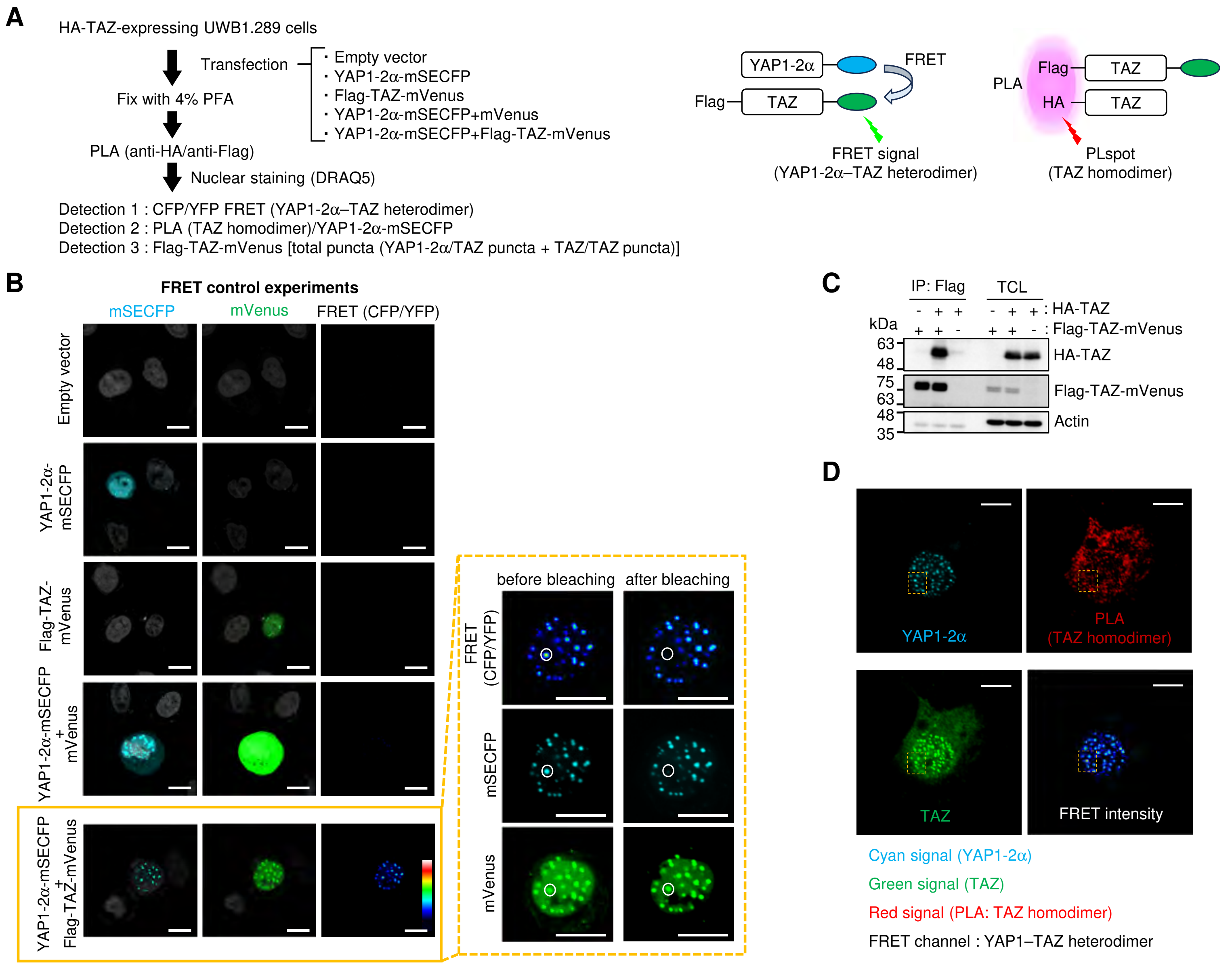


**Figure S5. Combined FRET and PLA analysis of nuclear YAP1-2α condensates (Related to Figure 2)**
(**A–D**) Related to Figures 2F-2H. Schematic overview of the visualization of YAP1-2α–TAZ heterodimers and TAZ homodimers using a combination of FRET and PLA (A). HA-TAZ-expressing UWB1.289 cells transfected with the indicated plasmids were subjected to FRET control experiments. CFP/YFP FRET signals were detected exclusively in cells co-expressing YAP1-2α-mSECFP and Flag-TAZ-mVenus. Photobleaching of the circled region in the FRET donor channel abolished the FRET signal in the corresponding region without affecting the FRET acceptor excitation signal (B). Parental or HA-TAZ-expressing UWB1.289 cells transfected with Flag-TAZ-mVenus or an empty vector were subjected to co-immunoprecipitation analysis. Homodimerization between HA-TAZ and Flag-TAZ-mVenus was assessed by immunoblotting of anti-Flag immunoprecipitates. Co-immunoprecipitation analysis was performed once to biochemically confirm the PLA results (C). Low-magnification images corresponding to Figure 2F. Higher-magnification views of the regions indicated by the yellow dashed lines are presented in Figure 2F (D). All scale bars represent 10 μm.

Supplemental Figure 6

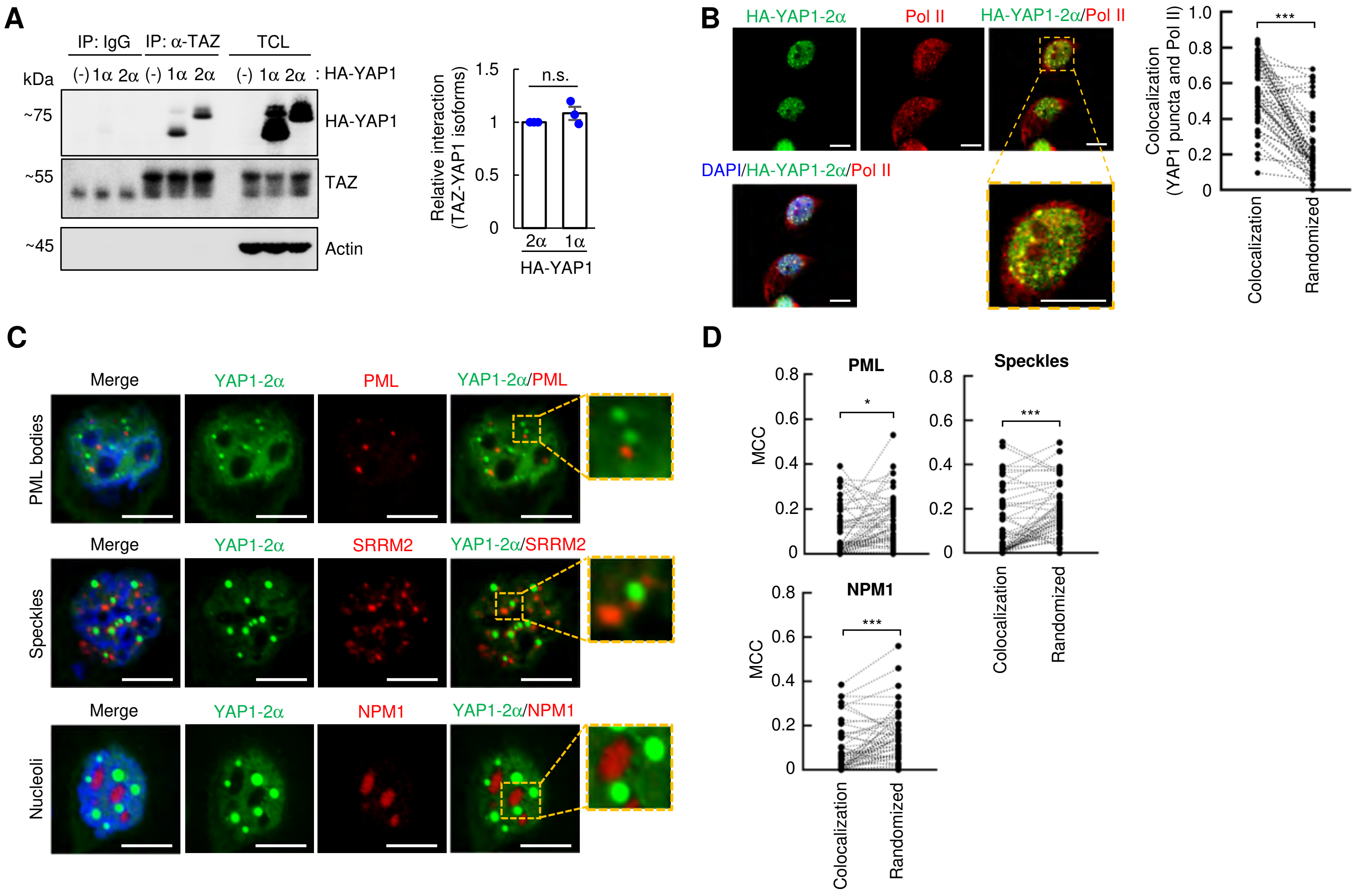


**Figure S6. YAP1-2α condensates are spatially distinct from known nuclear bodies but overlap with Pol II (Related to Figure 3)**

(**A**) Interactions between TAZ and YAP1-1α or YAP1-2α in MDA-MB-436 cells were assessed by anti-HA immunoblotting of anti-TAZ immunoprecipitates (left). The relative interaction between YAP1-1α and TAZ was calculated by setting the YAP1-2α intensity relative to TAZ intensity in the IP sample to one (right). $n$ = 3 biological replicates; n.s., not significant (Student's $t$ test). Error bars indicate the mean ± SEM.

(**B**) Immunofluorescence images of HA-YAP1-2α (green) and Pol II (red) (left panel). Manders' colocalization coefficient (MCC) was quantified in 50 cells. The observed colocalization (Pol II in YAP1-2α puncta) was significantly greater than expected under random distribution as determined by Costes randomization analysis (randomized M1) (right panel). ***$p < 0.001$ (paired $t$ test).

(**C** and **D**) Immunofluorescence images of YAP1-2α-mVenus (green) and nuclear bodies (red), including PML bodies (PML), nuclear speckles (SRRM2), and nucleoli (NPM1) (C). MCC was quantified in 55 cells. The observed colocalization of PML bodies, nuclear speckles, or nucleoli with YAP1-2α puncta was significantly lower than expected under a random distribution, as determined by Costes randomization analysis (randomized M1) (D). ***$p < 0.001$; *$p < 0.05$ (paired $t$ test).

All scale bars represent 10 μm.

Supplemental Figure 7

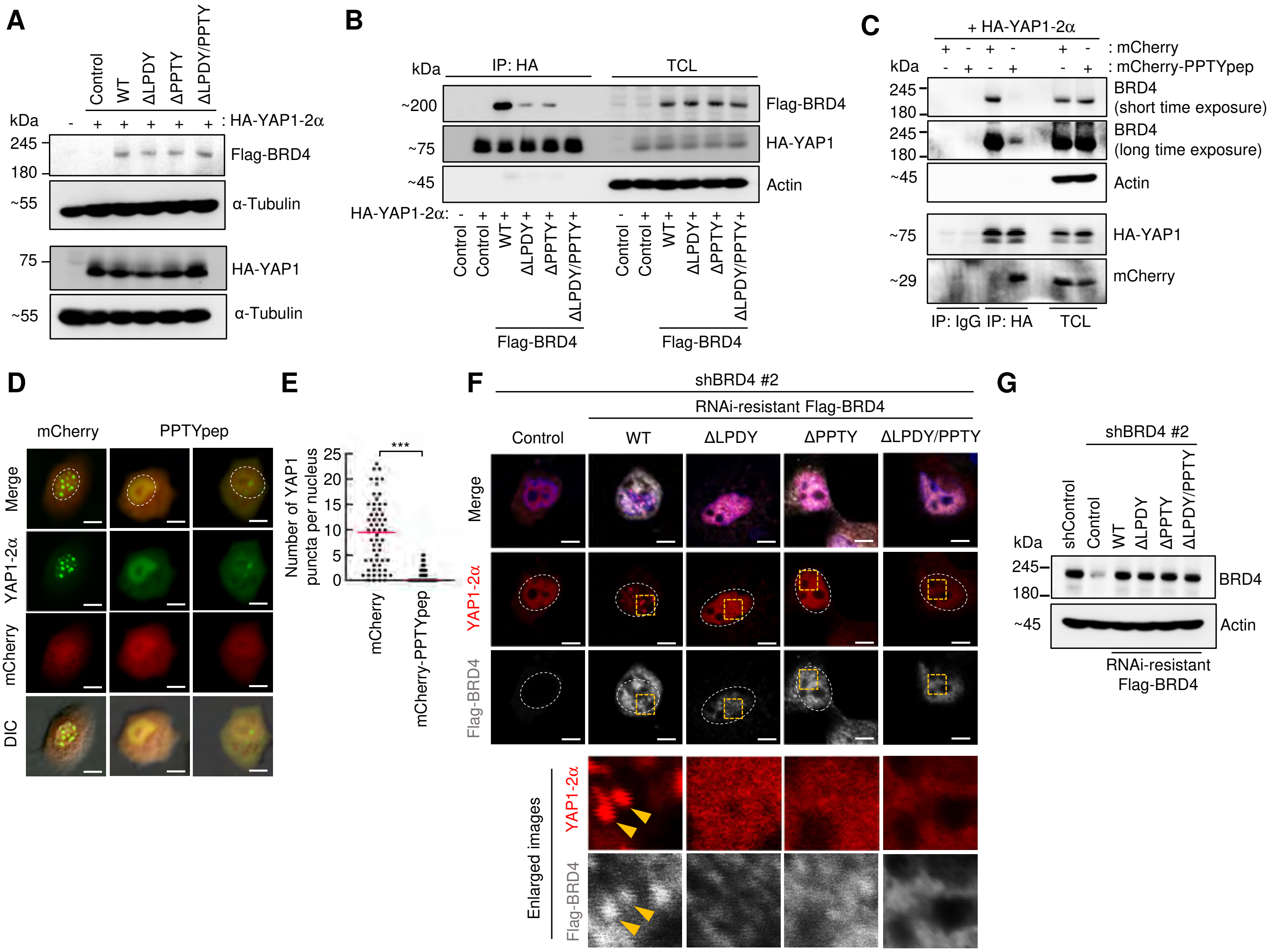


**Figure S7. WW domain-binding motifs in BRD4 are required for YAP1-2α condensate formation (Related to Figure 3)**

(**A**) Related to Figure 3C. Expression levels of HA-YAP1-2α and Flag-BRD4 mutants were assessed by immunoblotting. The same cell lysates were resolved on separate SDS-PAGE gels. After transfer, the membrane was cut and the indicated proteins were detected separately. (**B**) Related to Figure 3C. Interactions between HA-YAP1-2α and the indicated Flag-BRD4 mutants in UWB1.289 cells were assessed by anti-Flag immunoblotting of anti-HA immunoprecipitates. HA-YAP1-2α-expressing UWB1.289 cells were transfected with the indicated Flag-BRD4 mutants and immunoprecipitated with an anti-HA antibody. Total cell lysates (TCL) served as input controls. (**C**) Competitive inhibition of the YAP1-2α–BRD4 interaction with a PPTY peptide derived from the BRD4 sequence. HA-YAP1-2α-expressing cells were transfected with either the mCherry or mCherry-fused PPTY (mCherry-PPTYpep) vector, followed by sequential immunoprecipitation and immunoblot analyses using the indicated antibodies. PPTYpep disrupted the YAP1-2–BRD4 interaction without affecting endogenous BRD4 expression levels. (**D** and **E**) Inhibition of YAP1-2α puncta formation by PPTYpep. UWB1.289 cells were co-transfected with YAP1-2α-mVenus and either the mCherry or mCherry-PPTYpep vector (D). The effect of PPTYpep on YAP1-2α nuclear puncta formation was assessed by immunofluorescence analysis. $n = 60$ cells per group from three independent biological replicates (E). ***$p < 0.001$ (Mann-Whitney U test). (**F** and **G**) Related to Figure 3F. Rescue of the shBRD4-mediated reduction in YAP1-2α puncta formation by an shBRD4-resistant BRD4 construct. Immunofluorescence images of YAP1-2α (red), Flag-BRD4 (gray), and nuclei (blue). Enlarged images of the regions indicated by yellow dashed boxes are shown in the right panels. Yellow arrows indicate colocalization of YAP1-2α puncta and BRD4 (F). Expression of shBRD4 (#2)-resistant Flag-BRD4 and BRD4 knockdown efficiency were confirmed by immunoblotting (G). All scale bars represent 10 μm.

Supplemental Figure 8

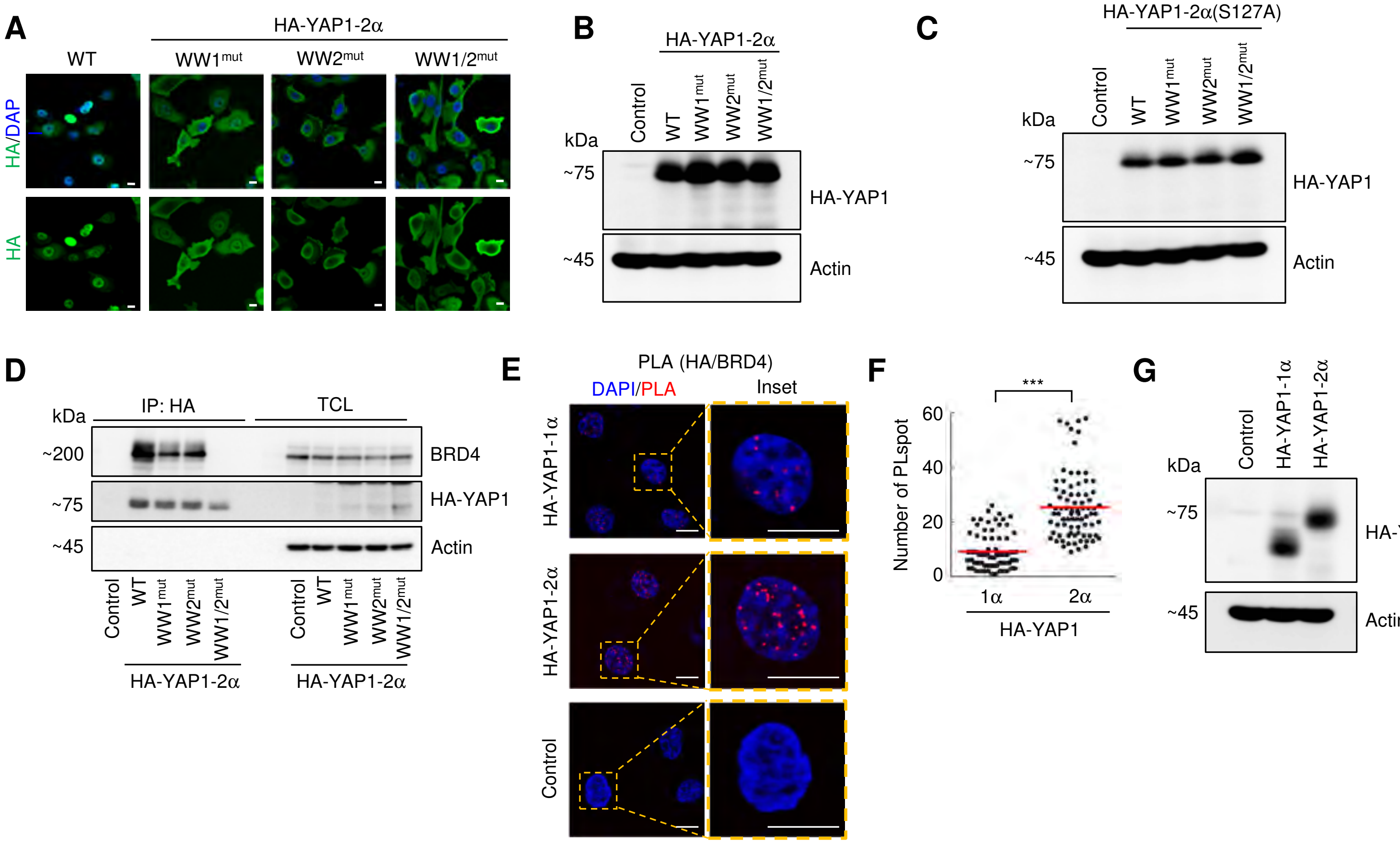


**Figure S8. WW domain-mediated multivalent interactions between YAP1-2α and BRD4 are important for YAP1 condensate formation (Related to Figure 3)**

(**A** and **B**) UWB1.289 cells were infected with lentiviruses encoding HA-YAP1-2α(WT), HA-YAP1-2α(WW1$^{mut}$), HA-YAP1-2α(WW2$^{mut}$), or HA-YAP1-2α(WW1/2$^{mut}$). Immunofluorescence images of HA-YAP1-2α(WT) and the WW domain mutants (green) are shown (A). Expression of HA-YAP1-2α(WT) and the WW domain mutants was confirmed by immunoblotting (B).

(**C** and **D**) Related to Figures 3G and 3H. UWB1.289 cells were infected with lentivirus encoding HA-YAP1-2α(S127A), HA-YAP1-2α(S127A|WW1$^{mut}$), HA-YAP1-2α(S127A|WW2$^{mut}$), or HA-YAP1-2α(S127A|WW1/2$^{mut}$). Expression of HA-YAP1-2α(S127A) and the WW domain mutants was confirmed by immunoblotting (C). Interactions between endogenous BRD4 and the indicated HA-YAP1-2α(S127A) mutants were assessed by immunoblotting anti-HA immunoprecipitates with anti-BRD4. Co-immunoprecipitation analysis was performed once to biochemically confirm the PLA results (D).

(**E–G**) *YAP1*-KO UWB1.289 cells were infected with lentiviruses encoding the indicated HA-tagged YAP1 isoforms. Interactions between HA-YAP1-1α or HA-YAP1-2α and BRD4 were analyzed by PLA. Red dots indicate PLA signals representing HA-YAP1/BRD4 interactions, and nuclei were stained with DAPI (blue) (E). Quantification of the PLA signals shown in Figure S8E is presented in the graph (F). $***p < 0.001$ (Mann-Whitney U test), $n = 80$ nuclei per group. Red lines indicate the mean. Expression of HA-YAP1-1α and HA-YAP1-2α was confirmed by immunoblotting (G).

All scale bars represent 10 μm.

### Supplemental Figure 9

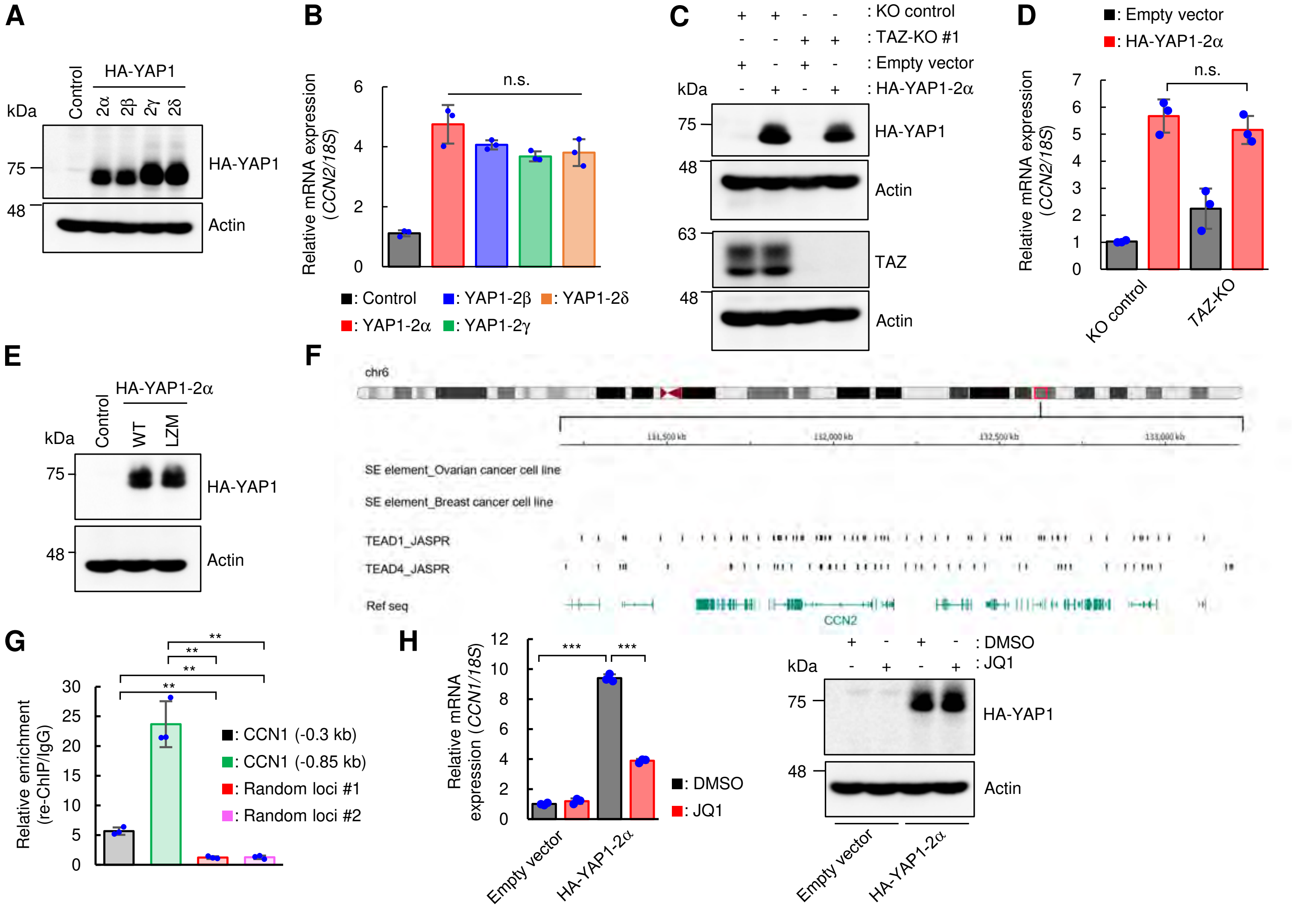


**Figure S9. YAP1-2α–TAZ heterodimerization drives gene expression programs (Related to Figure 4)**

(**A** and **B**) Related to Figure 4A. Expression of the HA-YAP1 isoforms was confirmed by immunoblotting (A). Expression of the canonical TEAD target gene *CCN2* was quantified by qPCR (B).

(**C** and **D**) Related to Figure 4B. Expression of HA-YAP1-2α and loss of TAZ expression after *TAZ* knockout were confirmed by immunoblotting (C). Duplicate aliquots of the same cell lysates were resolved on the same SDS-PAGE gel, and the transferred membrane was cut to allow separate detection of endogenous TAZ and the HA-tagged YAP1-2α protein. Expression of the canonical TEAD target gene *CCN2* was analyzed by qPCR (D).

(**E**) Related to Figure 4C. Expression of HA-YAP1-2α(WT) and HA-YAP1-2α(LZM) was confirmed by immunoblotting.

(**F**) Prediction of TEAD-positive candidate super-enhancer-associated regions near the *CCN2* gene. Super-enhancer elements identified by ROSE in serous ovarian cancer cells (GSM3328052: PEO1 cells) and triple-negative breast cancer cells (GSM6132824: MDA-MB-436 cells) are shown in the first and second tracks from the top, respectively. Putative TEAD-binding motifs were predicted using JASPAR. No putative TEAD-binding motifs were identified within candidate super-enhancer elements near the *CCN2* gene.

(**G**) Related to Figure 4H. Enrichment of the YAP1-2α–TAZ complex at candidate super-enhancer-associated regions upstream of the *CCN1* gene (-0.3 kb and -0.85 kb from the TSS), as well as at genomic regions lacking TEAD-binding motifs (random loci #1 and #2) was assessed by qPCR using ChIP-re-ChIP DNA samples from Figure 4H. Relative enrichment was calculated by dividing the qPCR value for the ChIP-re-ChIP DNA by the value for the corresponding IgG control.

(**H**) The effect of JQ1 treatment on the YAP1-2α-mediated increase in *CCN1* expression was assessed by qPCR (left). Expression of HA-YAP1-2α was confirmed by immunoblotting (right).

(B, D, G and H) ***$p < 0.001$; **$p < 0.01$; n.s., not significant (Welch's $t$ test). Error bars indicate the mean ± SD ($n = 3$ technical replicates). Plots show relative quantification (RQ) values for each sample.

Supplemental Figure 10

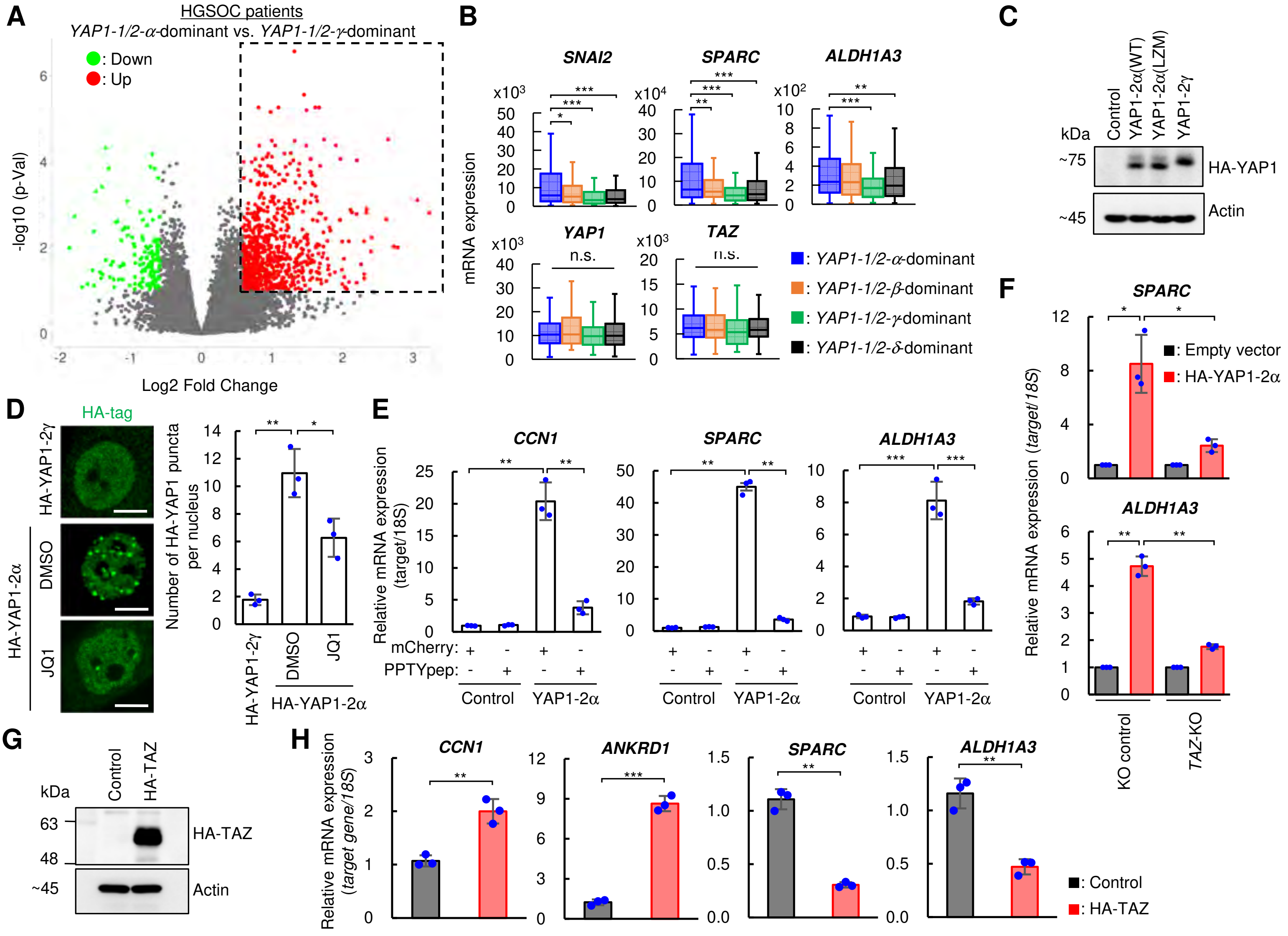

**Figure S10. YAP1-2α activates EMT-related genes in a TAZ- and BRD4-dependent manner (Related to Figure 5)** (**A** and **B**) Related to Figure 5A. HGSOC patients ($n$ = 425) were divided into four groups: "YAP1-1/2α dominant" ($n$ = 131), "YAP1-1/2β dominant" ($n$ = 83), "YAP1-1/2γ dominant" ($n$ = 78), and "YAP1-1/2δ dominant" ($n$ = 133). A volcano plot shows differentially expressed genes between the YAP1-1/2α dominant and YAP1-1/2γ dominant groups (FDR < 0.1, FC > 1.5 or FC < 0.67) (A). Box-and-whisker plots show mRNA expression levels of the EMT-related genes *SNAI2*, *SPARC*, and *ALDH1A3*, as well as *YAP1* and *TAZ*, in HGSOC patients grouped by YAP1 isoform. Boxes represent the 25th-75th percentiles, bars indicate the means, and whiskers indicate the data range (B). (**C**) Related to Figure 5B. Expression of the HA-YAP1 isoforms was confirmed by immunoblotting. (**D**) The effect of JQ1 treatment on YAP1-2α puncta formation was assessed by immunofluorescence staining (left). HA-tagged YAP1 is shown in green. Scale bars represent 10 μm. Quantification of nuclear YAP1 puncta is shown on the right. Plots show the mean number of nuclear YAP1 puncta per experiment ($n$ > 100 nuclei; three biological replicates). $^{**}p$ < 0.01; $^{*}p$ < 0.05 (Welch's $t$-test). (**E**) qPCR analysis of *CCN1*, *SPARC*, and *ALDH1A3* expression—the three representative target genes associated with candidate regulatory elements—with or without competitive inhibition of the YAP1-2α–BRD4 interaction using PPTYpep shown in Figures S7C-S7E. (**F**) *TAZ*-KO UWB1.289 cells were transfected with HA-YAP1-2α or an empty control vector. *SPARC* (top) and *ALDH1A3* (bottom) expression levels were assessed by qPCR. Relative expression levels were calculated by setting the empty vector-transfected cells to 1. (**G** and **H**) Ectopic HA-TAZ overexpression in UWB1.289 cells. Immunoblotting analysis (G). The effects of TAZ overexpression were assessed by qPCR analysis of the canonical TEAD target genes *CCN1* and *ANKRD1*, along with *SPARC* and *ALDH1A3*, representative genes associated with the candidate regulatory elements examined in this study (H). (B, E, F, and H) $^{***}p$ < 0.001; $^{**}p$ < 0.01; $^{*}p$ < 0.05 (Welch's $t$ test). Error bars show the mean ± SD. $n$ = 3 (technical replicates).

Supplemental Figure 11

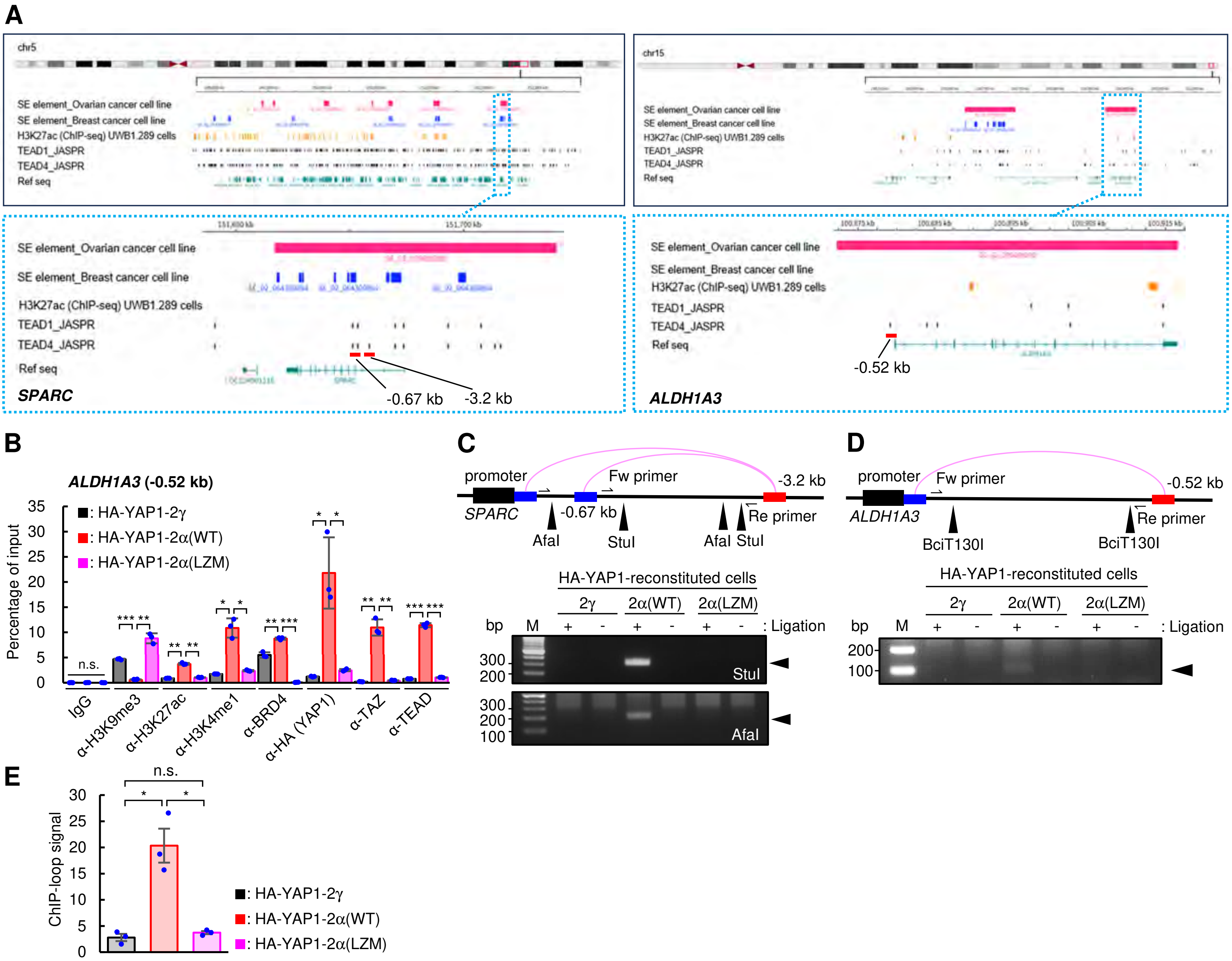


**Figure S11. YAP1-2α/TAZ/TEAD-associated candidate regulatory elements and EMT-related gene activation (Related to Figure 5)**

(**A**) Related to Figure 5C. Super-enhancer regions identified by ROSE in serous ovarian cancer cells (GSM3328052, PEO1 cells) and triple-negative breast cancer cells (GSM6132824, MDA-MB-436 cells) are shown in the first and second tracks of each panel, respectively. H3K27ac ChIP-seq data from UWB1.289 cells (GSM4728119) are shown in the third track of each panel. Candidate super-enhancer-associated regions upstream of *SPARC* and *ALDH1A3* are indicated by red lines in the enlarged panels enclosed by blue dashed boxes (left, *SPARC*; right, *ALDH1A3*). These annotations were used to nominate candidate regions for locus-specific analyses and do not represent de novo, genome-wide regulatory-element calling in UWB1.289 cells. (**B**) Related to Figure 5C. CUT&RUN-qPCR analysis of the indicated chromatin marks and factor enrichment at the region located 0.52 kb upstream of the *ALDH1A3* transcription start site in UWB1.289 cells reconstituted with the indicated YAP1 variants. ***$p$ < 0.001; **$p$ < 0.01; *$p$ < 0.05; n.s., not significant (Welch's *t*-test). Error bars indicate the mean ± SD. $n$ = 3 (technical replicates). (**C**) Related to Figure 5D. Representative agarose gel electrophoresis images of PCR amplicons from ChIP-loop DNA templates, showing interactions between the *SPARC* (-3.2 kb) region and the *SPARC* (-0.67 kb) region (top), and between the *SPARC* (-3.2 kb) region and the *SPARC* promoter (bottom). (**D** and **E**) ChIP-loop analysis was performed to detect interactions between the *ALDH1A3* (-0.52 kb) region and its promoter in YAP1-reconstituted UWB1.289 cells. Representative agarose gel electrophoresis images (D). Quantification of ChIP-loop signals (E). *$p$ < 0.05; n.s., not significant (Welch's *t*-test). Error bars indicate the mean ± SEM. $n$ = 3 (biological replicates).

Supplemental Figure 12

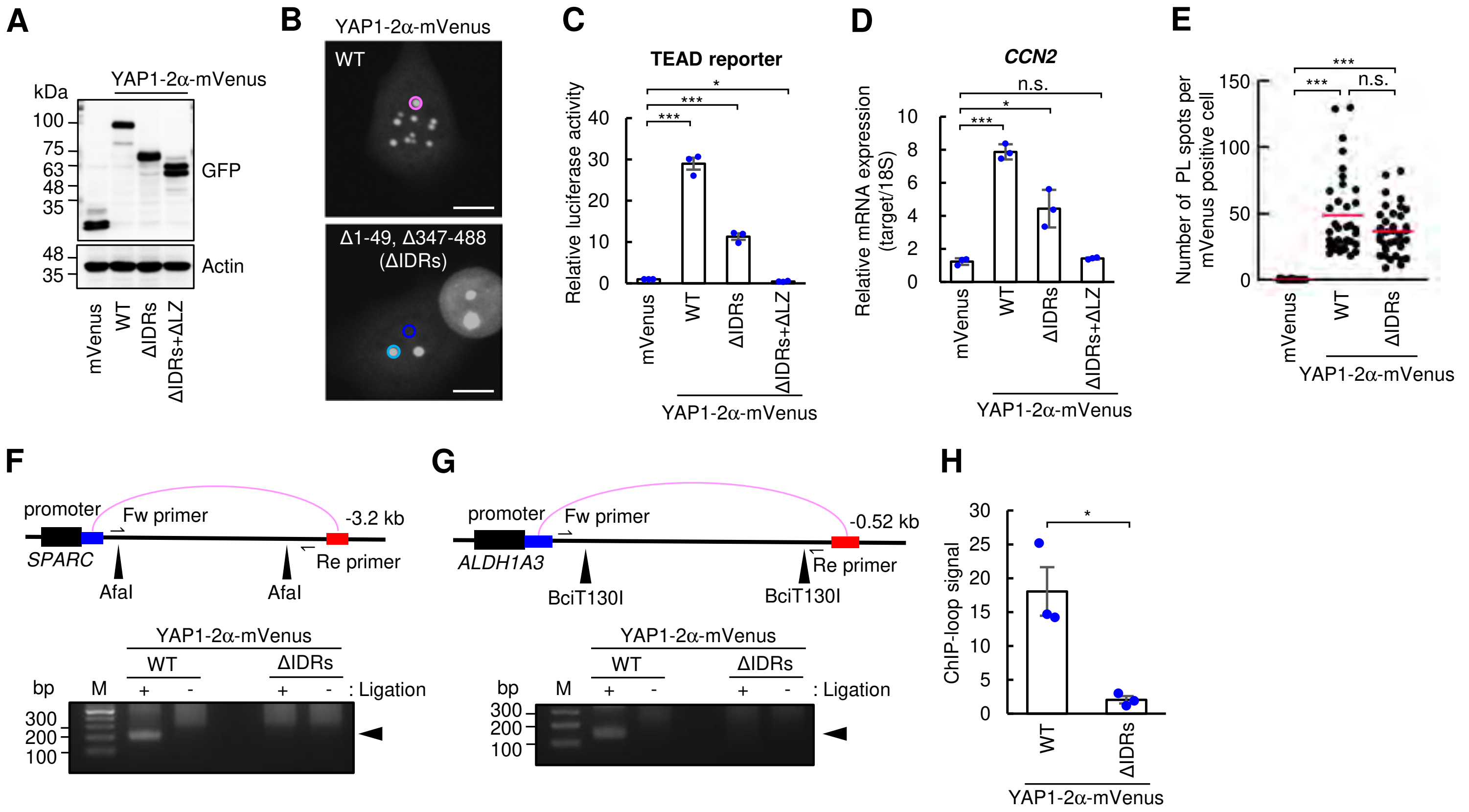


**Figure S12. IDRs promote YAP1-2α nuclear condensate formation and activation of YAP1-2α/TAZ/TEAD-associated regulatory elements (Related to Figure 5)**

(**A**-**E**) Related to Figures 5E-5J. Analysis of IDR-deficient YAP1-2α mutants in *YAP1*-KO UWB1.289 cells. Expression of the IDR-deletion mutants was confirmed by immunoblotting (A). Representative whole-nucleus images of cells used for FRAP analysis (B). TEAD reporter assay (C). qPCR analysis of *CCN2* expression (D). Interactions between BRD4 and mVenus, YAP1-2α(WT)-mVenus, or YAP1-2α(ΔIDRs)-mVenus were assessed using PLA. Quantification of PLA signals is shown in the graph. Red bars indicate the mean (E).

(**F**-**H**) Related to Figure 5J. ChIP-loop analysis was performed to detect interactions between the *SPARC* (-3.2 kb) region and its promoter (F), and between the *ALDH1A3* (-0.52 kb) region and its promoter (G) in *YAP1*-KO UWB1.289 cells transfected with the indicated YAP1-2α constructs. Quantification of ChIP-loop signals shown in Figure S12G is presented in (H).

(C and D) ***$p$ < 0.001; *$p$ < 0.05; n.s., not significant (Welch's $t$ test). Error bars indicate the mean ± SD. $n$ = 3 (technical replicates).

(E) ***$p$ < 0.001; n.s., not significant (Mann-Whitney U test), $n$ = 35 mVenus-positive cells per group.

(H) *$p$ < 0.05 (Welch's $t$ test). Error bars indicate the mean ± SEM. $n$ = 3 (biological replicates).

Supplemental Figure 13

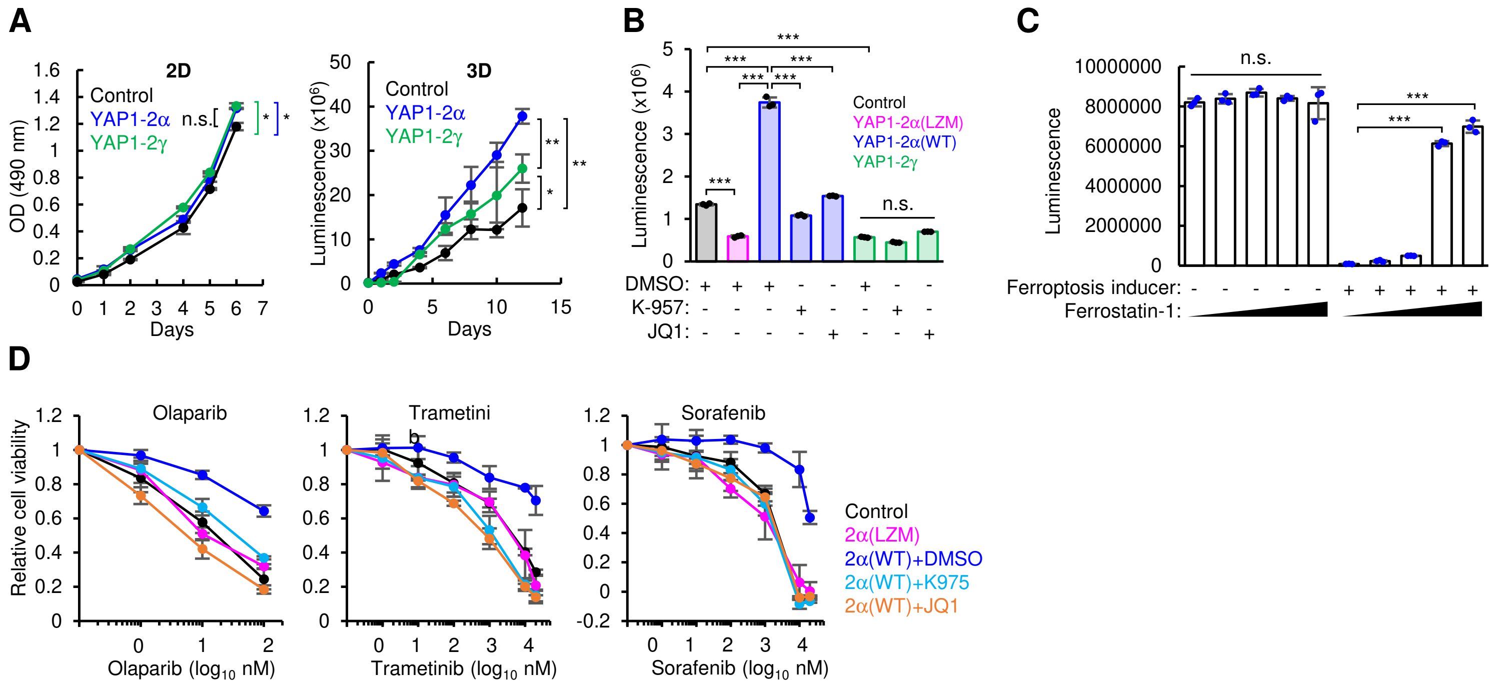


**Figure S13. YAP1-2α promotes cancer-associated phenotypes in vitro (Related to Figure 6)**

(**A**) Two-dimensional (2D) or three-dimensional (3D) cell proliferation of YAP1-2α-overexpressing, YAP1-2γ-overexpressing, and control UWB1.289 cells. 2D proliferation was measured by an MTS assay (left), whereas anchorage-independent 3D growth was assessed by measuring ATP content (right).

(**B**) Cell chemotaxis was evaluated with a Transwell assay, and migrated cells were quantified by measuring ATP levels.

(**C**) Related to Figure 6D. Chlorido[N, N'-Disalicylidene-1,2-Phenylenediamine]Iron(III)-induced ferroptosis was validated using the ferroptosis inhibitor ferrostatin-1 (final concentrations: 0, 1, 10, 100, and 1,000 nM).

(**D**) The effects of the YAP1-2α/TAZ/TEAD transcriptional axis on the sensitivity of UWB1.289 cells to various anticancer drugs. Cells were treated with JQ1 or K-975 in combination with the indicated anticancer drugs, or with DMSO as a vehicle control. Olaparib (PARP inhibitor), trametinib (MEK inhibitor), and sorafenib (tyrosine kinase inhibitor) were used.

(A-C) ***$p < 0.001$; **$p < 0.01$; *$p < 0.05$; n.s., not significant (Welch's $t$ test). $n = 3$ (technical replicates).

All error bars indicate the mean ± SD.

Supplemental Figure 14

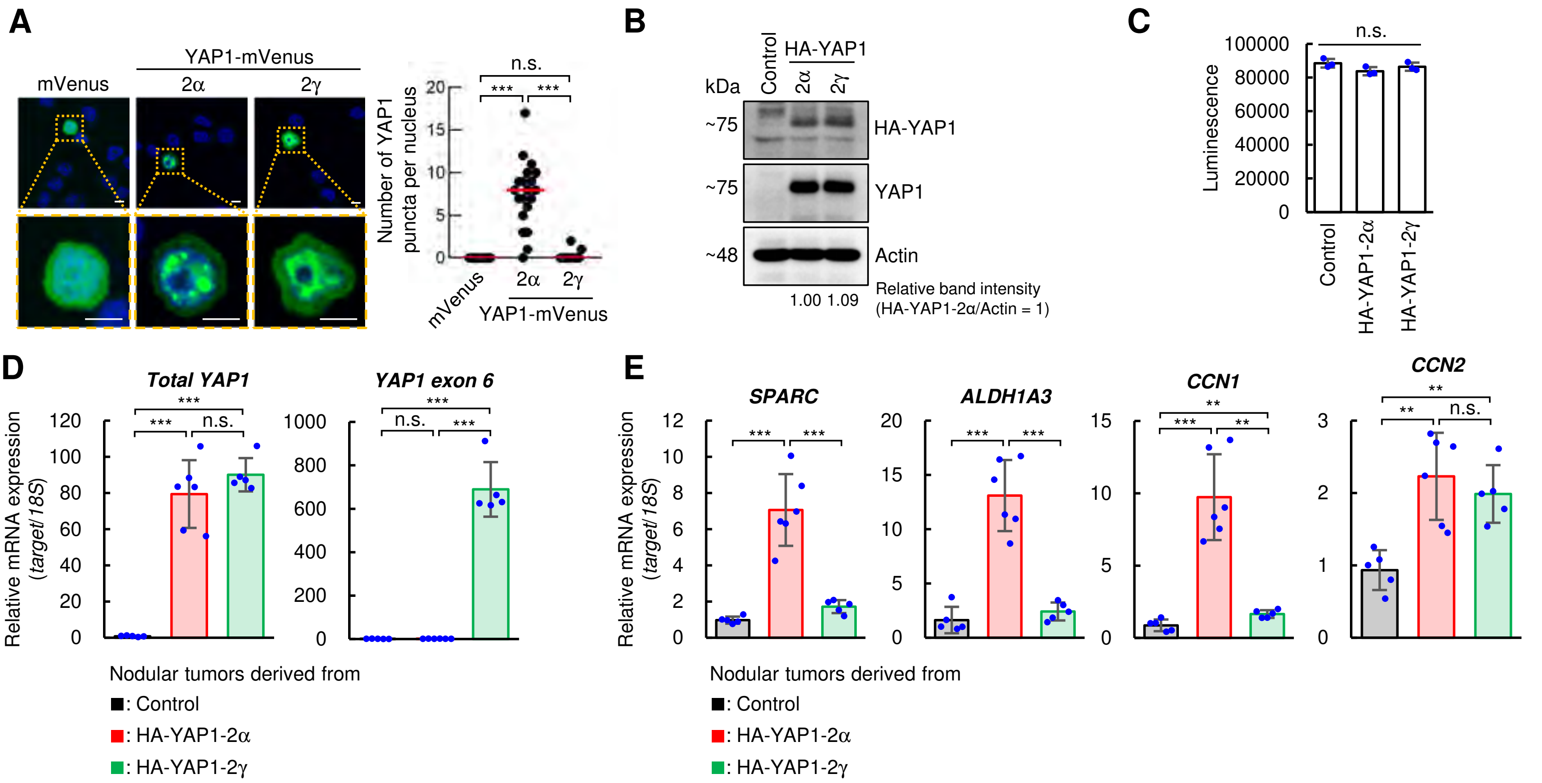


**Figure S14. YAP1-2α promotes peritoneal dissemination in diffuse-type gastric cancer cells (Related to Figure 6)**

(**A**) Representative fluorescence images of MKN45 cells that stably express firefly luciferase (MKN45-Fluc) after transfection with the indicated mVenus-fused YAP1 isoforms (left), and quantification of nuclear YAP1 puncta (right). $n$ = 20 nuclei per group. ***$p$ < 0.001; n.s., not significant (Mann-Whitney U test). Red lines indicate the mean. YAP1 puncta were quantified using predefined criteria for mVenus-fused YAP1 as described in the STAR Methods (Immunofluorescence Staining section). Scale bars represent 10 μm.

(**B** and **C**) Related to Figures 6F-6I. MKN45-Fluc cells were transduced with lentiviral vectors encoding the indicated HA-tagged YAP1 isoforms. Expression of the HA-YAP1 isoforms was assessed by immunoblotting. Band intensities were normalized to Actin and expressed relative to the HA-YAP1-2α band intensity, which was set to 1 (B). Luciferase activity was measured in the corresponding cell lines. $n$ = 3 (technical replicates). n.s., not significant (Student's $t$ test) (C).

(**D** and **E**) Related to Figures 6F-6I. Ectopic expression of HA-YAP1 isoforms in tumor nodules was verified by qPCR analysis of total *YAP1* and exon 6-containing *YAP1* transcripts (D). qPCR analysis of canonical TEAD target genes and selected genes linked to candidate regulatory elements in tumor nodules (E). $n$ = 5 (tumor nodules derived from Control and HA-YAP1-2γ-expressing cells) or $n$ = 6 (tumor nodules derived from HA-YAP1-2α-expressing cells) technical replicates. ***$p$ < 0.001; **$p$ < 0.01; n.s., not significant (Welch's $t$ test). Plots show relative quantification (RQ) values for each sample.

All error bars indicate the mean ± SD.

Supplemental Figure 15

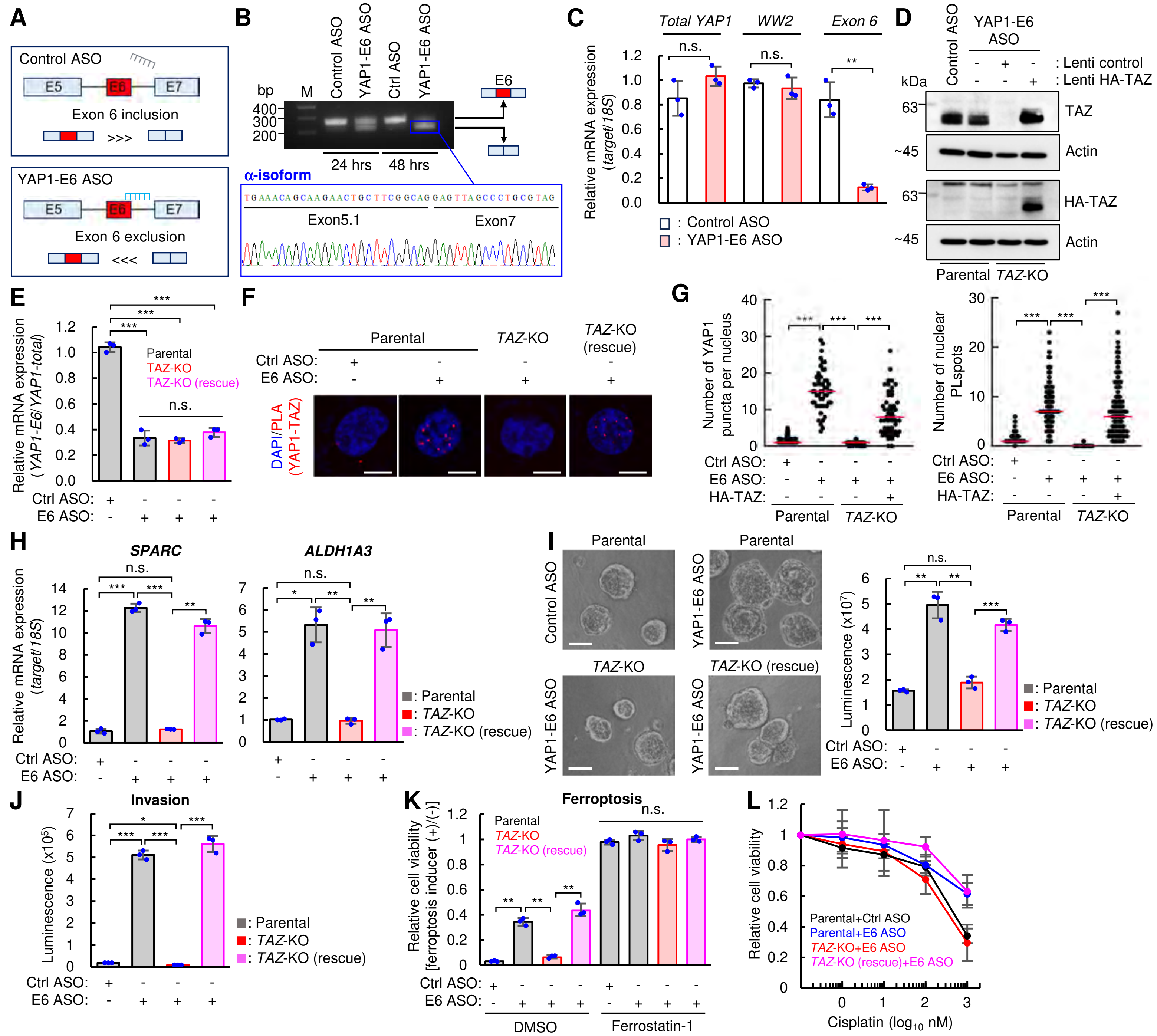


**Figure S15. Endogenous YAP1 exon 6 skipping promotes YAP1-2α-associated phenotypes (Related to Figure 7)** (**A-C**) Regulation of endogenous *YAP1* pre-mRNA splicing in UWB1.289 cells using an antisense oligonucleotide targeting *YAP1* exon 6. Schematic illustration of the ASO targeting *YAP1* exon 6 (A). ASO-induced skipping of *YAP1* exon 6 was confirmed by RT-PCR (top panel). The upper and lower bands represent exon 6 inclusion and exclusion, respectively. To identify the exon 6-skipping transcript, the lower-band RT-PCR amplicon was excised, purified, and subjected to Sanger sequencing (bottom panel) (B). Relative expression levels of total *YAP1*, WW2-containing *YAP1* transcripts, and exon 6-containing *YAP1* transcripts were quantified by qPCR (C). (**D-L**) Related to Figure 7A. Analysis of parental UWB1.289 cells, *TAZ*-KO cells, and HA-TAZ-rescued *TAZ*-KO cells after ASO-induced *YAP1* splice switching via exon 6 skipping. Validation of *TAZ* knockout and HA-TAZ expression by immunoblotting (D). Confirmation of *YAP1* exon 6 exclusion by qPCR (E). Visualization of the interaction between endogenous YAP1 and TAZ proteins by PLA (F). Quantification of results shown in panel F (left) and Figure 7A (right) (G). qPCR analysis of YAP1-2α target genes associated with candidate super-enhancers (H). Analyses of spheroid-forming capacity (I), Matrix invasion (J), ferroptosis (K), and cisplatin sensitivity (L). Data are mean ± SD; red bars indicate means. *$p < 0.05$, **$p < 0.01$, ***$p < 0.001$; n.s., not significant; Welch's *t*-test (C, E, and H-K) or Mann-Whitney U test (G). Three technical replicates (C, E, and H-K), $n = 55$ (left) or 95 (right) nuclei per group (G).

Supplemental Figure 16

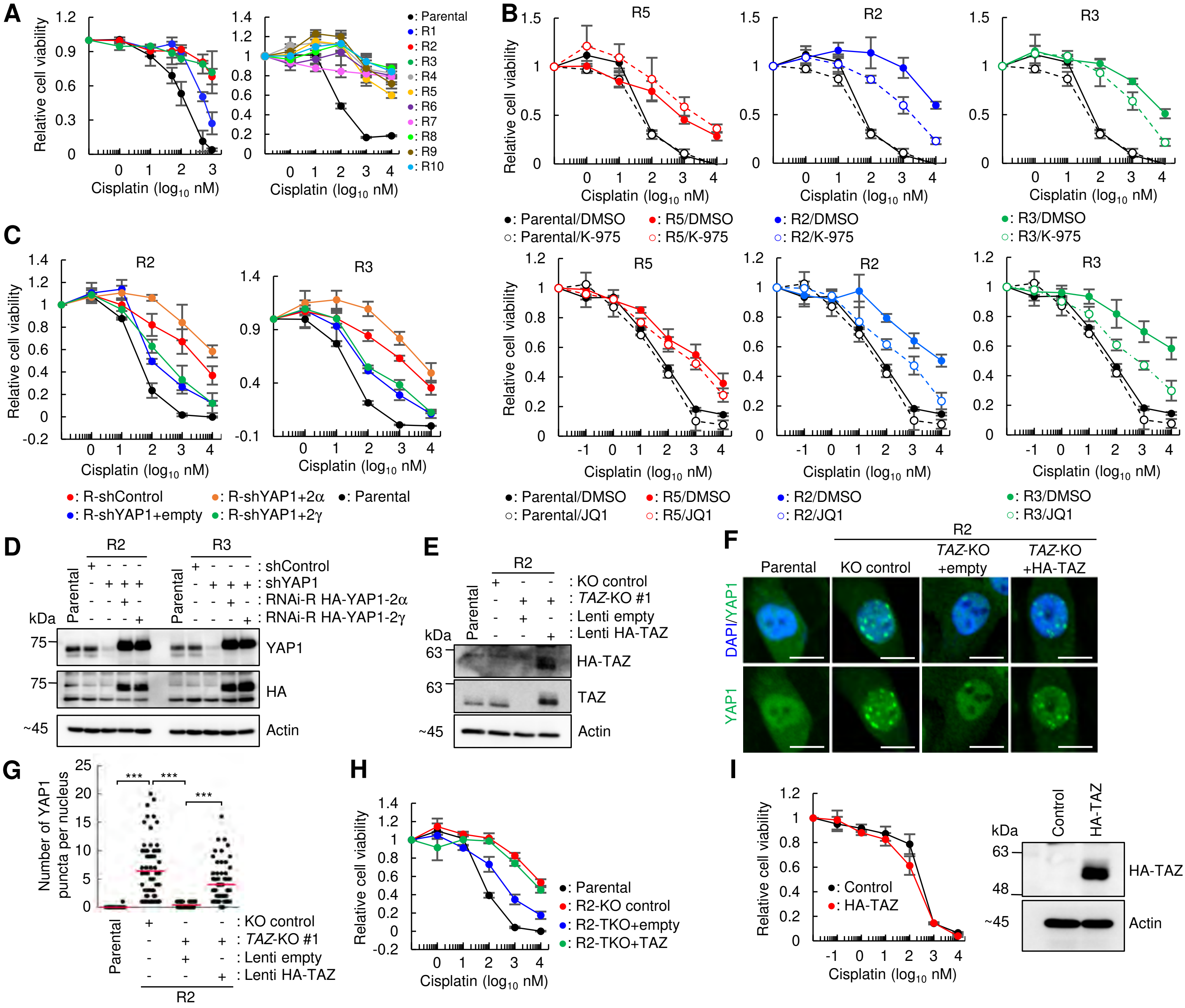


**Figure S16. A subset of cisplatin-resistant clones exhibits an endogenous YAP1 splicing switch (Related to Figure 7)**

(**A**) Sensitivity to cisplatin in independently established cisplatin-resistant clones was assessed using an MTS assay. (**B**) The effect of K-975 (top) or JQ1 (bottom) treatment on cisplatin sensitivity in the indicated cisplatin-resistant clones was assessed by MTS assay. (**C** and **D**) UWB1.289 cells transduced with lentiviral shRNA targeting *YAP1* or non-targeting shRNA were infected with the indicated RNAi-resistant HA-YAP1 isoforms. YAP1 expression levels were confirmed by immunoblot analysis (C). Cisplatin sensitivity was assessed by an MTS assay (D). (**E-H**) The *TAZ*-KO cisplatin-resistant UWB1.289 cell clone R2 was transduced with lentiviruses encoding HA-TAZ. TAZ-KO and HA-TAZ expression were confirmed by immunoblot analysis (E). The effect of *TAZ* knockout on endogenous YAP1 puncta formation in R2 was assessed. Representative immunofluorescence images of YAP1 (green) are shown in (F), with quantification of nuclear YAP1 puncta in parental and R2 cells in (G). Cisplatin sensitivity in the indicated cells was measured using an MTS assay (H). (**I**) The effect of ectopic TAZ expression on cisplatin sensitivity in parental UWB1.289 cells was assessed by an MTS assay (left). HA-TAZ expression was confirmed by immunoblotting (right).

Data are shown as mean ± SD; three technical replicates (A-C, H, and I). Red bars indicate means; ***$p < 0.001$; Mann-Whitney U test; $n = 60$ nuclei per group (G). All scale bars represent 10 μm.

Supplemental Figure 17

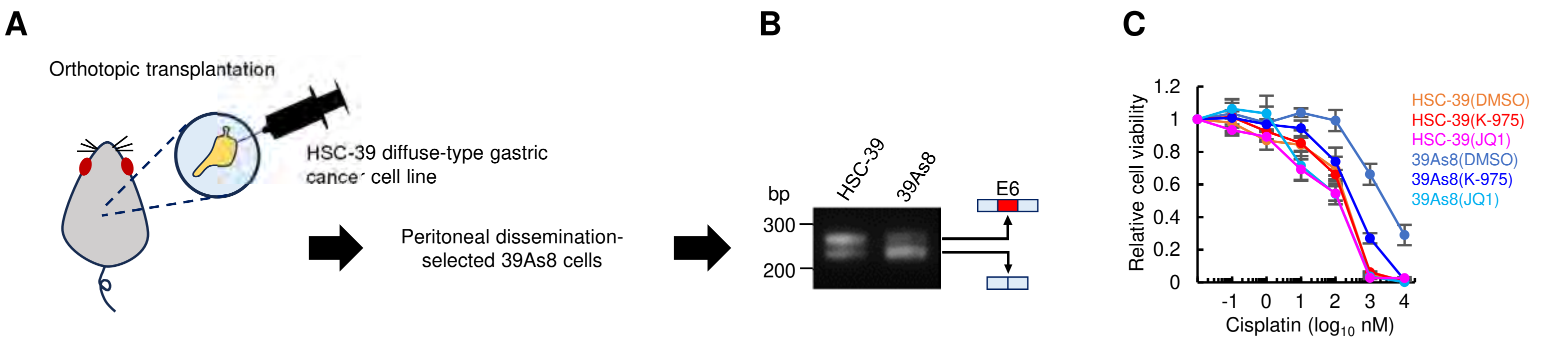


**Figure S17. Peritoneal dissemination selection is associated with an endogenous YAP1 splicing switch (Related to Figure 7)**

(**A–C**) Analysis of a peritoneal dissemination-derived cell line (39As8) established from peritoneal metastatic lesions generated after orthotopic transplantation of the diffuse-type gastric cancer cell line HSC-39 into nude mice. Experimental schematic (A). RT-PCR analysis of *YAP1* exon 6 splicing (B). The effects of JQ1 or K-975 treatment on cisplatin sensitivity in the indicated cells were assessed using an MTS assay (C). Data are mean ± SD; three technical replicates.

### Supplemental Figure 18

A

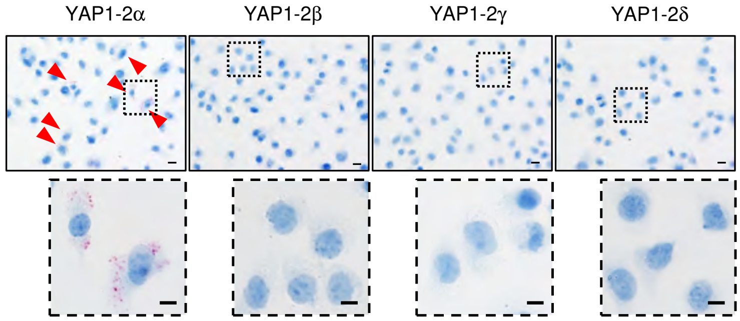


B

Stage

| YAP1-E5.1/E7 staining score | NAT/normal | I | II | III | IV |
|---|---|---|---|---|---|
| 0 | 10 | 11 | 6 | 2 | 0 |
| 1+ | 0 | 3 | 4 | 3 | 0 |
| 2+ | 0 | 2 | 2 | 2 | 1 |
| 3+ | 0 | 0 | 3 | 6 | 0 |

C D E

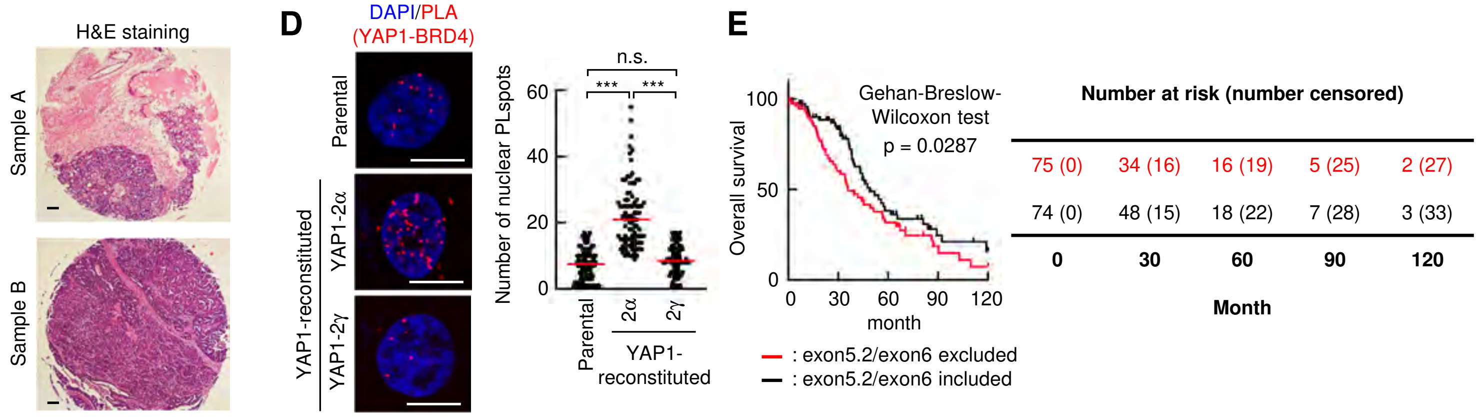


F

**Multivariable Cox proportional hazards regression analysis of OS**

| Covariates | HR | 95% CI | P value |
|---|---|---|---|
| YAP1 exon 5.2 (PSI) | 0.343 | 0.095 - 1.266 | 0.1078 |
| YAP1 exon 6 (PSI) | 0.484 | 0.190 - 1.271 | 0.1388 |
| YAP1 expression (TPM) | 0.980 | 0.626 - 1.538 | 0.9309 |
| Age | 1.024 | 1.011 - 1.036 | 0.0002 |
| Stage III vs. Stage II | 2.049 | 1.040 - 4.824 | 0.0369 |
| Stage IV vs. Stage II | 2.382 | 1.142 - 5.804 | 0.0189 |

Stage I (n = 1), Stage II (n = 24), Stage III (n = 318), Stage IV (n = 62), N.A. (n = 11).

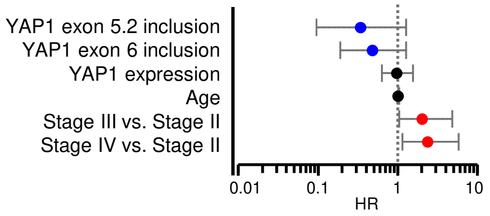


**Figure S18. *YAP1-E5.1/E7* expression in HGSOC correlates with advanced FIGO stage (Related to Figure 7)**

(**A**) Confirmation of BaseScope probe specificity for *YAP1-E5.1/E7* transcripts, which encode either YAP1-1α or YAP1-2α. BaseScope was performed on cells transfected with the indicated YAP1 isoforms. Red arrows mark BaseScope signal-positive cells. Scale bars represent 10 μm.

(**B**) Table of FIGO stage and *YAP1-E5.1/E7* staining score.

(**C**) Related to Figures 7G–7K. H&E staining of clinical HGSOC specimens, sample A and sample B, used for the analysis shown in Figure 7G–7L.

(**D**) PLA analysis of the YAP1–BRD4 interactions in parental and YAP1-reconstituted UWB1.289 cells. Representative PLA signals (left) and quantification (right). $n$ = 90 nuclei per group; $***p < 0.001$; n.s., not significant (Mann-Whitney U test). Red lines indicate the mean.

(**E**) Kaplan-Meier analysis of overall survival (OS) in the TCGA-HGSOC cohort. HGSOC patients were stratified into *YAP1* splicing subgroups based on inclusion or exclusion of exons 5.2 and 6: exon 5.2/exon 6 exclusion (exon 5.2 PSI < 0.35 and exon 6 PSI < 0.65, $n$ = 75) and exon 5.2/exon 6 inclusion (exon 5.2 PSI > 0.35 and exon 6 PSI > 0.65, $n$ = 74).

(**F**) Multivariable Cox proportional hazards regression analysis of overall survival (OS) in the TCGA-HGSOC cohort with available TCGA SpliceSeq data ($n$ = 416). Stage I cases were excluded because only one specimen was available (n = 1), precluding reliable statistical evaluation. Covariates included *YAP1* exon 5.2 and exon 6 inclusion levels (PSI), *YAP1* mRNA expression (TPM), age at diagnosis, and FIGO stage (stage III or IV versus stage II). Hazard ratios (HRs) are shown as points, and 95% confidence intervals (CIs) as bars. In the right panel, red and blue indicate variables associated with increased and decreased risk, respectively. The left panel summarizes the multivariable Cox regression results.

Supplemental Figure 19

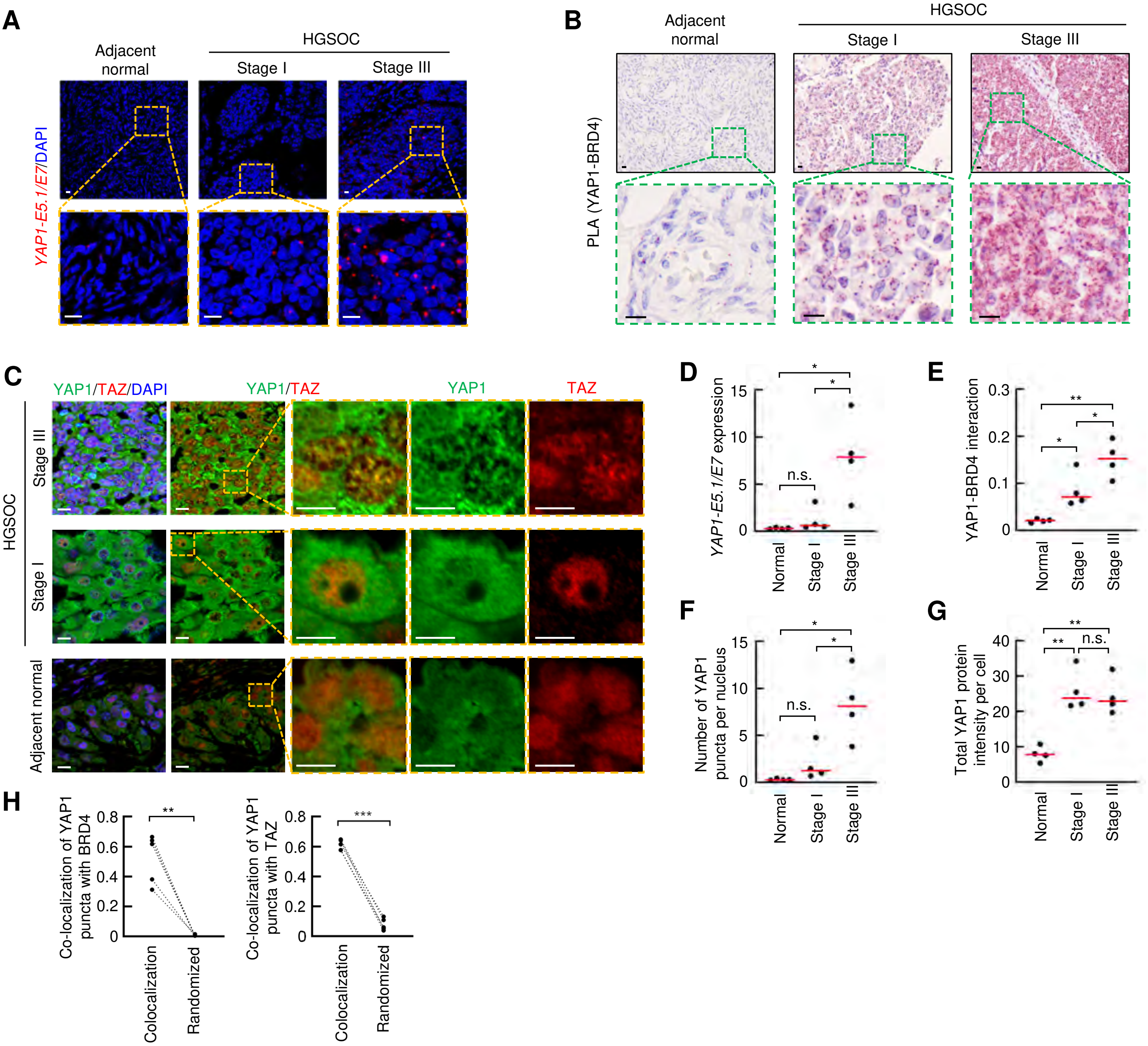


**Figure S19. BRD4 and TAZ colocalize with nuclear YAP1 condensates in HGSOC specimens (Related to Figure 7)**

(**A–H**) Independent validation analyses were performed using adjacent normal tissues ($n$ = 4) and HGSOC specimens from patients with FIGO stage I ($n$ = 4) or stage III ($n$ = 4). These samples were subjected to the analyses described below. *YAP1-E5.1/E7* transcripts were detected by BaseScope (A). YAP1–BRD4 interactions were detected by PLA (B). Immunofluorescence analysis of YAP1 (green) and TAZ (red) was performed (C). Correlations between FIGO stage and *YAP1-E5.1/E7* expression levels (D), YAP1–BRD4 interactions (E), the number of YAP1 puncta per nucleus (F), and total YAP1 protein intensity per cell (G) are shown. Colocalization analysis of YAP1 puncta with BRD4 (left) and TAZ (right) was performed using the Coloc2 plugin. $n$ = 5 YAP1 puncta-positive specimens (FIGO stage I, $n$ = 1; FIGO stage III, $n$ = 4), defined as specimens with >3 YAP1 puncta per nucleus (H).

Red bars indicate the mean. $^{***}p < 0.001$; $^{**}p < 0.01$; $^{*}p < 0.05$; n.s., not significant. Welch's $t$ test (D–G) or paired $t$ test (H). $n$ = 4 specimens per group (D–G). All scale bars represent 10 μm.
